# Integrative genomic and epigenomic analyses identified IRAK1 as a novel target for chronic inflammation-driven prostate tumorigenesis

**DOI:** 10.1101/2021.06.16.447920

**Authors:** Saheed Oluwasina Oseni, Olayinka Adebayo, Adeyinka Adebayo, Alexander Kwakye, Mirjana Pavlovic, Waseem Asghar, James Hartmann, Gregg B. Fields, James Kumi-Diaka

## Abstract

The impacts of many inflammatory genes in prostate tumorigenesis remain understudied despite the increasing evidence that associates chronic inflammation with prostate cancer (PCa) initiation, progression, and therapy resistance. The overarching goal of this study was to identify dysregulated inflammatory genes that correlate with PCa progression and decipher their molecular mechanisms as well as clinical significance in PCa using integrative genomics, transcriptomics, and epigenomics approach. Our Weighted Gene Co-expression Analysis (WGCNA) and multivariate analysis identified 10 inflammatory genes: IRAK1, PPIL5/LRR1, HMGB3, HMGB2, TRAIP, IL1F5/IL36RN, ILF2, TRIM59, NFKBIL2/TONSL, and TRAF7 that were significantly associated with PCa progression. We explored the potentials of IRAK1 and other inflammatory genes as diagnostic and/or prognostic biomarkers by performing both KM survival and AUROC curve analyses. Our results indicate the clinical significance of these inflammatory genes in predicting the development and progression of PCa. IRAK1 was found to be overexpressed and hypomethylated in most PCa samples. A significantly high percentage of castration-resistant PCa (CRPC) and neuroendocrine PCa (NEPC) samples display copy number variations, especially amplification of the IRAK1 gene compared to the indolent prostate adenocarcinoma (PRAD) samples. Furthermore, we identified missense and frameshift mutations of IRAK1 in a few PRAD samples with potential functional implications. In conclusion, the results from this study suggest that IRAK1 dysregulation may be an important contributor to chronic prostatitis (inflammation) and PCa progression.

## Introduction

Prostate cancer (PCa) is the number one cause of cancer-related deaths and the second most diagnosed cancer in US men (Siegel, et al., 2021). One of the major drawbacks to the accurate diagnosis and treatment of inflammation-driven PCa progression is our limited understanding of the molecular mechanisms underlying aberrant inflammation signaling in PCa patients (Pezaro et al., 2014; Sciarra et al., 2008). Since asymptomatic chronic inflammation is hard to diagnose, we do not have adequate knowledge of how specific genes within the inflammatory cascade drive PCa initiation and progression (Schiller & Parikh, 2011; Wagenlehner et al., 2007). Integrative bioinformatics analysis involves the integration of different software, tools, and databases to identify anomalies, patterns, and correlations within large datasets to answer various biological questions and predict disease outcomes or future trends (Hasin et al., 2017). In this study, we adopted an integrative bioinformatics pipeline to analyze genomic and epigenomic data to identify inflammatory genes involved in prostate tumorigenesis.

Omics data contain vital information, such as the genetic variation, gene expression, and DNA methylation of various oncogenes and tumor suppressor genes associated with clinical traits (Heindl et al., 2015). These data are usually disconnected; hence, we took an integrative bioinformatics approach that combined genetic, gene expression, and epigenetic data to explore inflammatory genes that are associated with PCa progression. This approach is expected to provide effective diagnostic and prognostic models to predict, manage, and prevent inflammation-driven PCa progression (Yang et al. 2015; Robinson et al. 2015). DNA methylation is an epigenetic modification that has been suggested to play an important role in regulating gene expression and therefore could be adopted as an alternative biomarker to gene expression (genomic) profiling (Koch et al., 2015; Héninger et al., 2015). Despite this knowledge, we still do not have an adequate understanding of the impacts of DNA methylation of inflammatory genes in PCa development and progression.

Specifically, this study aims to characterize inflammatory genes as predictors of PCa progression, with a particular focus on the impact of aberrant signaling of interleukin-1 receptor-associated kinases (IRAKs) on prostate tumorigenesis. We harnessed the richness of the large-scale genomic, transcriptomic, and methylation data available via The Cancer Genome Atlas (TCGA) and the cBioPortal for Cancer Genomics (cBioPortal) databases to identify the gene alterations, expression, and methylation patterns of inflammatory genes associated with PCa progression like IRAK1 (Cerami et al., 2012; Gao et al., 2012; Weinstein et al., 2013). We also assessed the correlation between inflammation-associated genes and clinical features/traits such as progression and biochemical recurrence/cancer relapse.

We applied weighted gene co-expression network analysis (WGCNA) to study and construct biological networks and generated clusters or modules containing genes that are highly correlated with PCa progression and relapse using a set of criteria designed into our bioinformatics analysis pipeline (Langfelder & Horvath, 2008). We then identified inflammation-associated genes within intramodular “hubs” of genes for further downstream analysis and investigative studies. The WGCNA approach deployed in this study identified significant modules and co-expression patterns among genes associated with PCa progression and chronic inflammation. We have identified an inflammatory signature of 10 genes (located in the magenta module) associated with PRAD progression and focused on IRAK1 for further downstream explorative analysis. Our study provides a framework for future mechanistic studies into the role of inflammatory genes in PCa progression (Xu et al., 2020). Understanding the pattern of genomic, epigenomic, and proteomic alterations of chronic inflammatory genes associated with PCa progression may provide us with effective prognostic and diagnostic tools to manage chronic inflammation-associated PCa (Tan et al., 2018).

## Materials and Methods

### Data sourcing from databases and bioinformatics pipeline design

R studio Desktop v1.4.1106 (https://www.rstudio.com/), an open-access R programming language software platform for statistical computing was downloaded and installed on a MacBook computer to perform many of our integrated bioinformatics and statistical analyses (R studio Core Team, 2020). R packages and their dependencies needed for each analysis were also installed from Bioconductor software v3.14 (https://www.bioconductor.org/install/) and the Comprehensive R Archive Network (CRAN) repository (https://cran.r-project.org/). The data used in this study were obtained from the TCGA (https://portal.gdc.cancer.gov/) and the cBioPortal (http://cbioportal.org) databases. The cBioPortal database contains about 20 PCa cohort studies that we explored for the genetic alteration analysis portion of this study.

For most of our analyses, the PanCancer Whole Exome sequencing (WES) and RNAseq datasets (n = 494) acquired from TCGA were utilized. The RNAseq dataset from matched normal prostate samples (n = 52) from PRAD patients as well as unmatched from GTEx (n = 245) were also added as the control group during the differential gene expression analysis (DGEA) to evaluate the clinical significance of gene expression between the primary tumor samples vs normal samples. The mRNA expression reads were derived from an Illumina HiSeq 2000 sequencer and aligned to Gencode hg19/GRCh37 to generate gene counts (overlapping reads), which are then Batch normalized to RSEM (RNASeqV2 by Expectation Maximization) for further analysis based on predetermined bioinformatics pipeline. An advantage of RSEM is that it can be used to estimate expression levels of genes and their isoforms by performing RSEM-prepare-reference and RSEM-calculate-expression analysis. In addition to the genomic (RNAseq, WES, and methylation) datasets, TCGA and cBioPortal also provide metadata of clinical traits associated with the patients or tumor samples. After downloading all these documents, we designed a bioinformatics workflow or pipeline as briefly summarized in **Figure 1**. The Transcript counts were calculated using the Flux Capacitor program. Normalized and unnormalized RSEM counts were extracted for each prostate adenocarcinoma (PRAD) patient and developed into a matrix for further analysis.

**Figure 1:**
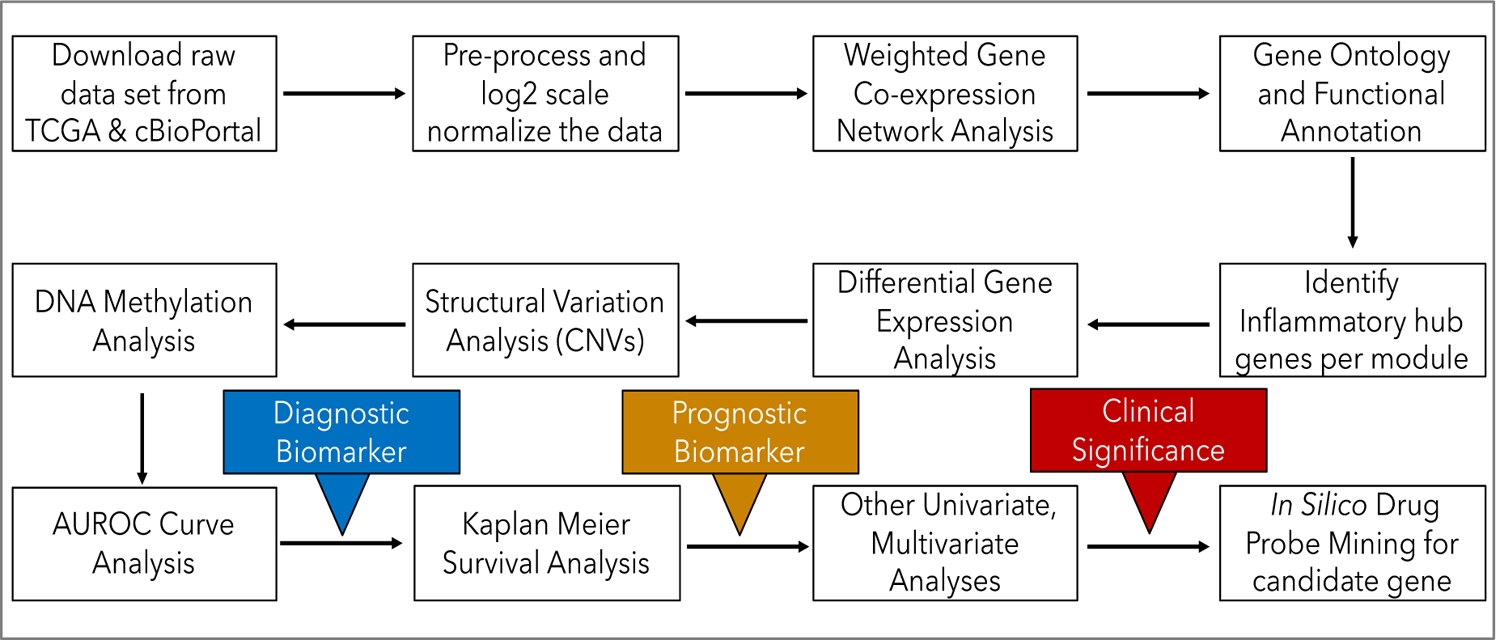
Flow chart summarizing the bioinformatics pipeline designed for this study.

### Batch effect identification and correction

The protocols employed by TCGA for data collection and processing include transportation of biospecimens of Tumor and Normal tissue samples and their clinical data from patient donors through TCGA Biospecimen Core Resources (BCR) laboratories to various Tissue Source Sites (TSS) to ensure proper ethics and ensure that specimens meet the TCGA biospecimen criteria. Upon receiving these biospecimens, those that meet the required quality were cataloged, processed, and stored for further analysis. Any patient identifying information was removed during the biospecimen processing. Thereafter, biospecimens were grouped into batches based on the type of cancer and each batch was identified by analytes and transported to sequencing centers for sequencing and processing into usable data. The output data were transferred electronically to the TCGA Genome Characterization Centers and Genome Sequencing Centers (CGCC and GSC) for interpretation and meta-analysis. The CGCC was involved in analyzing genetic changes involved in cancer while high-throughput TCGA Genome Sequencing Centers identify the changes in the DNA sequence associated with specific cancers before sharing it with the public through the TCGA Data Portal.

Before performing any analysis, MBatch, an R programming package (https://bioinformatics.mdanderson.org/public-software/mbatch/) and Batch Effect Viewer (https://bioinformatics.mdanderson.org/public-software/tcga-batch-effects/) were used to assess, diagnose, and correct for any batch effects and data variance from the different sequencing centers in which the downloaded TCGA data were initially acquired or processed. MBatch has a statistical capability to analyze and quantify the presence of batch effects in the dataset. It does this by using algorithms such as Hierarchical Clustering and Principal Component Analysis. Dispersion Separability Criterion (DSC) quantifies the amount of batch effect in data similar to the Scatter Separability Criterion. DSC is a measure of the ratio of dispersion between batches (Db) and within dispersion (Dw) (i.e., DSC = D_b_/D_w_). A higher DSC value means that there is a greater dispersion between batches than within batches. This means there is a higher probability of the samples within batches to be more identical to each other than the batches themselves are to each other. When the DSC values are significantly below 0.5, it means a weak batch effect, but the batch effect will have to be considered when the DSC is higher than 0.5 with a significant p-value (Dy et al., 2004, Wodarz et al., 2014). DSC p-value provides information on the presence of outliers that may skew DSC values. The null hypothesis stating that there are no batch effects in the data set will be rejected if the p-value is found to be less than the 0.05 significance threshold.

### Data cleaning, outliers removal, and detecting low-quality counts

Following batch correction and effect viewing using MBatch and Batch Effect Viewer, TAMPOR was used to perform deep data cleaning to ensure that our final input data is devoid of any incomplete data or outlier that may skew our outputs. TAMPOR (https://github.com/edammer/TAMPOR) helps to maintain the integrity of our data by performing robust batch effect correction through the removal of batch artifacts and batch-wise variance as well as the removal of genes or samples with ≥ 50 percent missing or zero values, and removal of replicates or cluster outliers. At the end of TAMPOR analyzes, our total gene has been reduced from 20,506 to 17,794 genes, and the number of patients/samples from 494 to 472.

### Logarithm transformation and normalization of PRAD dataset

Following removal of outliers and low-quality data, we log_2_ transformed (i.e., log_2_ (expression value +1)) and quantile normalized our data using the Limma package or DESeq package in R. Read Counts were further converted into expression values such as Counts Per Million, Reads Per Million, Reads Per Kilobase Millions (RPKM), Fragment Per Kilobase Million (FPKM), and Transcripts Per Kilobase Million (TPM), depending on the processing requirement of the analysis being performed.

### Gene clustering and networking analysis using Weighted Gene Co-expression Network Analysis (WGCNA)

Before running the WGCNA, the clinical data containing phenotypic information about the clinical traits of each sample was downloaded and curated. The non-numeric variables were converted to numeric or binary units (**Supplementary Method S1**). The WGCNA package was installed from the Comprehensive R Archive Network (CRAN), the standard repository for R add-on packages. To run the WGCNA package, the installation tools/algorithms for the needed packages were obtained from Bioconductor and installed in R (**Supplementary Method 2**).

WGCNA algorithm was used to identify gene co-expression networks in our dataset by robustly calculating the eigengene, bicor rho, and p values for each module and then correlating the first principal component of each module (module eigengenes) with clinical traits or phenotypes of interest. WGCNA does this by constructing a sample dissimilarity matrix (1-topology overlap) and grouping genes that have similar expression patterns within the patient cohort together. The network construction was performed using WGCNA blockwiseModules function with parameters as follows: WGCNA dynamic tree-cutting algorithm, CutreeHybrid, Power achieving scale-free topology=10, Sd out=3, Adjust Threshold=0.002, mergeHeight=0.15, MaxBlocksize=25000. PAMstage=True, DeepSplit=2, minModuleSize=100, Net=blockwiseModules(t(cleanDat), Power=power, DeepSplit=ds, Verbose=3, PamStage=PAMstage, SaveTOMs=FALSE, CorType=”bicor”, NetworkType=”signed”, MergeCutHeight=mergeHeight, PamRespectsDendro=TRUE, TOMDenom=”mean, ReassignThresh=0.05, Biweight midcorrelation “bicor”, biweight midcorrelation was used as opposed to Pearson correlation to robustly correlate with less weight given to outlier measures (Ohandjo et al., 2019). The WGCNA R-script and outputs for this study can be visualized or downloaded from GitHub via https://github.com/soseni2013/WGCNA-for-Cancer.

### Gene ontology and upstream/downstream regulator analysis

The biological functions, cellular processes, and molecular functions of each module were predicted using WebGestalt (http://www.webgestalt.org/), an R package, and GO-Elite, a Python program (http://www.genmapp.org/go_elite/help_main.htm). Visualization of the outputs was performed using a custom R script. WebGestalt (WEB-based Gene SeT AnaLysis Toolkit) is a functional enrichment analysis web tool that supports three well-established and complementary methods for enrichment analysis, including Over Representation Analysis (ORA), Gene Set Enrichment Analysis (GSEA), and Network Topology-based Analysis (NTA) (Wang et al., 2017; Zhang et al., 2005). GSEA involves using predefined lists for classifying genes of interest, such as biological processes, cellular component categories, and molecular functions, and testing for statistical overrepresentation of the category members, in this case to gene lists based on module membership. A gene set is defined as any group of genes that share a common biological function while enrichment analysis is an important step in computational biology for inferring knowledge about an input gene set by matching it to annotated or known gene sets from prior experimental or computational studies.

Enrichment analysis checks whether the input set of genes significantly overlaps with annotated gene sets. WebGestalt was supplemented with GO Elite to perform a second itinerary gene ontology enrichment analysis on our module of interest to identify the functional annotations and biological functions of genes located in the module (Zambon et al., 2012). In addition to the standard ensemble v62 database with 3 standard ontology categories, the GSEA molecular signature C2 database (v6.2) was used as a reference to identify the association of network modules with the curated lists related to published studies with a varying focus on health and disease, particularly cancer-dysregulated gene lists. The GSEA C3 database was also used to identify enriched upstream and downstream regulators among the members of each module.

### Module-based differential gene expression analysis

Differential gene expression analysis was performed to compare PRAD patients who progressed (n = 93) versus those who did not progress (n = 401). A non-parametric ANOVA (Kruskal-Wallis) was used to remove noisy/skewed RNAseq data and to accommodate the unequal sample sizes in each group. Fold change (FC) between the two groups was calculated for genes located in each module. A volcano plot of the -log p-value on the y-axis vs the FC (the difference between the gene expression values of the progressed and non-progressed) was made for each module.

Furthermore, differential gene expression analysis was performed for the normalized data to show inflammatory gene expression profiles between prostate tumor and paired adjacent normal or unpaired non-cancer normal samples using DESeq2 package in R (https://bioconductor.org/packages/release/bioc/html/DESeq.html) and the expression pattern among cancer patients/samples visualized using TNM plotter (Bartha & Győrffy, 2021) as well as Heatmap3 package in R studio interface. Correlations between the inflammatory genes of interest and known PCa-associated genes were determined using the CorrPlot package in R to plot correlograms for the genes in indolent, metastatic/castration-resistant, and neuroendocrine/lethargic PCa patients. Also, principal component analysis for dimensionality reduction was used to identify the pattern of expression between the PRAD and normal/non-cancer samples (Tang et al., 2017).

### Functional enrichment analysis of inflammatory genes in the module of interest

After identifying the differential gene expression pattern between patients with progressive and non-progressive tumors, we decided to determine if the deregulation of chronic inflammation genes in this module can contribute to progression. Enrichr (https://maayanlab.cloud/Enrichr/) was used to validate enrichment analysis performed using WebGestalt and Go-Elite. Enrichr allows for profiling of gene ontology of interest whereby we selected those associated with the inflammatory super pathway axis (Kuleshov et al., 2016). Using this method, Enrichr provides the functional annotation of the genes that are heavily involved in inflammatory signal transduction. We then created a set of criteria that each gene must meet to be included in the inflammatory signature, followed by the ranking of genes according to how they perform and then selecting the most significant gene in the gene set based on our ranking system. Some of the criteria used for ranking include kME (Eigengene) value, inflammation score, oncoscore, and whether signaling through the TLR/IL-1R/NF-κB super pathway.

### Upstream and downstream regulator analysis using Ingenuity Pathway Analysis (IPA), Cytoscape, and String_db software

To identify the biological function of the significantly associated modules to traits of interest, we investigated the genes within the module of interest participating in the same biological process, such as inflammation regulation. Since we have identified IRAK1 as our preferred candidate, we conducted upstream regulator analysis using IPA software (QIAGEN; https://www.qiagenbioinformatics.com/products/ingenuitypathway-analysis). IRAK1 was found to be regulated or interactive with genes in the TLR/IRAK/NF-κB pathway while other non-canonical pathways were also identified. We used Enrichr and String_db to enrich cancer-associated and inflammation-associated genes within our gene set which was curated and filtered to select a few that have a stronger confidence score and significant adjusted p-value. String_db network analysis tool was then used to predict their regulatory interaction with IRAK1, which provided us a better insight into the biological and molecular functions of IRAK1 with regards to tumorigenesis (Szklarczyk et al., 2019). Cytoscape is open-source software for integrating biomolecular interaction networks with high-throughput expression data into a unified conceptual framework. Cytoscape was also used to identify the co-expression or regulatory interaction between IRAK1 and other neighboring genes in the magenta module (Shannon et al., 2003).

### Genetic alteration analyses of IRAKs in PRAD patients/samples

To determine whether the overexpression in IRAK1 was caused due to genetic alterations, the MutSigCV and GISTIC (v. 2.0) algorithmic tools were used to identify somatic mutations and copy number alterations/variations (CNVs/CNAs – deep or shallow deletion, gains, diploid, and amplifications), respectively, from a total of 20 PCa cohort studies of 6044 patients (6329 samples) downloaded from cBioPortal database (Lawrence et al., 2013; Mermel et al., 2011; Cerami et al., 2012). The genomic and clinical data sets were preprocessed and analyzed as previously described in our methodology. Variant calling of somatic mutations and copy number variation had been previously performed and the annotations of the mutations standardized using Genome Nexus and Canonical UniProtKB Transcript. Following the validation of genetic alterations, Mutation Assessor (MA; http://mutationassessor.org/r3/), Catalogue Of Somatic Mutations In Cancer (COSMIC, https://cancer.sanger.ac.uk/cosmic), Polymorphism Phenotyping v2 (PolyPhen-2; http://genetics.bwh.harvard.edu/pph2/), and Sorting Intolerant from Tolerant (SIFT; https://sift.bii.a-star.edu.sg/) variant calling and analysis tools were used to predict the functional impact of observed mutations, *in silico* (Adzhubei et al., 2013; Gnad et al., 2013; Ng et al., 2003; Sim et al, 2012; Sondka et al., 2018).

Mutation Assessor can predict the functional impact of amino-acid substitutions in proteins, such as mutations discovered in cancer or missense polymorphisms based on the evolutionary conservation of affected amino acid in protein homologs. SIFT predicts whether an amino acid substitution in naturally occurring nonsynonymous polymorphisms or laboratory-induced missense mutations affects protein function based on sequence homology and the physical properties of amino acids. PolyPhen-2 predicts the possible impact of an amino acid substitution on the structure and function of a human protein using straightforward physical and comparative considerations. The functional impact scores for SIFT, PolyPhen, COSMIC, and Mutational Assessor were integrated and ranked to form a new scoring method (severe, moderate, and mild) for our analysis. Mutual exclusivity and co-occurrence trends between altered gene pairs within a gene set per patient were determined by calculating the odds ratio (OR), which reflects the probability that a gene pair is mutually exclusive or co-occurring. The significance of the relationship between a gene pair is determined by Fisher’s exact test (p < 0.05).

To determine the transcriptomic effects of oncogenic somatic mutations in PRAD samples, the Mann-Whitney U test was performed for 17,794 genes using the “TARGET” analysis module of the muTarget software (http://www.mutarget.com/) to identify differentially expressed genes between mutated and wild-type patient cohorts of PRAD. The output of the analysis generated a list of genes whose mutation status was significantly (p < 0.05) associated with upregulation or downregulation of the IRAK1 gene. Boxplots were generated to show the distribution of the most significant genes using the “ggplot2” package to visualize expression differences (Nagy and Győrffy, 2021).

### Differential methylation analysis of IRAK1 in PRAD dataset

The Illumina Infinium Human Methylation 450k BeadChip (Illumina 450K array) prostate adenocarcinoma dataset was downloaded from the TCGA consortium database. SMART (Shiny Methylation Analysis Resource Tool; http://www.bioinfo-zs.com/smartapp/), DNMIVD (DNA Methylation Interactive Visualization Database; http://www.unimd.org/dnmivd/), Wanderer (http://maplab.imppc.org/wanderer/), and MEXPRESS (https://mexpress.be/), and tools were used to identify, analyze, visualize and compare significant DNA methylation lesions at the CpG promoter and transcriptional regions in IRAK1 between PRAD and normal patients (Díez-Villanueva et al., 2015; Ding et al., 2020; Koch et al., 2015; Li et al., 2019). CpG islands were predicted by searching the IRAK1 sequence one nucleotide base at a time, scoring each dinucleotide (+17 for CG and −1 for others), and then identifying the maximally scoring segments. Each scored segment was then evaluated for the following criteria for GC content of 50% or greater and base-pair length of greater than 200 bp as well as observed/expected CpG ratio greater than 60%. The observed CpG is the number of CpG dinucleotides in a segment while the expected CpG is calculated by multiplying the number of ‘C’ and the number of ‘G’ in a segment and then dividing the product by the length of the segment.

Pairwise correlation analysis was performed to explore the correlation between IRAK1 gene expression or transcript level (log_2_ scale (TPM+1) values) and DNA methylation with significance derived when the beta (b) values or methylation (m) values of CpG probes are less than 0.05. Pearson, Spearman, and Kendall correlation methods were employed for correlation analysis. Both the IRAK1 gene expression and methylation data of PRAD (n = 492) and normal (n = 50) samples were included for the correlation analysis. Also, the mean methylation (aggregation) value for all CpGs was calculated and tested for statistical significance (Wilcoxon test, p < 0.05).

Following the estimation of gene-level CNV of IRAK1 using the GISTIC2 threshold method. The estimated values representing homozygous deletion (−2), single copy deletion (−1), diploid normal copy (0), low-level copy number amplification(+1), and high-level copy number amplification (+2) were correlated with the methylation values of IRAK1.

### Predicting the clinical significance of IRAK1 and other inflammatory genes from the magenta module in PRAD samples/patients

To evaluate the diagnostic and prognostic significance of IRAK1 in PRAD samples or patients, we performed an Area Under the Receiver Operator Characteristic (AUROC) analysis using EasyROC software, which helps to predict the diagnostic potential of our gene as a progression and PRAD biomarker (Goksuluk et al., 2016). Univariate and multivariate proportional hazards regression (Cox regression) model and KM Plotter were used to assess the correlation between IRAK1 gene expression and the overall survival status (OS), the progression-free survival status (PFS), and relapse-free survival (RFS)/disease-free survival status (DFS) (Győrffy et al., 2010). The OS_STATUS refers to the overall survival status (“0” −> “living” or “1” −> “deceased”) and the event of interest is death from PRAD. This provides a very broad sense of the mortality of the groups. The OS_MONTHS indicates the number of months from time of diagnosis to time of death or last follow-up. The event of interest for DFS or RFS is the relapse of a disease rather than death (“0” -> “Alive or dead tumor-free”, “1” -> “Dead or alive with tumor” because patients may have relapsed but are not deceased. DFS curves are usually lower than OS curves. PFS_STATUS uses the progression of a disease as an end-point (i.e., tumor growth or spread). It indicates whether the patient’s disease has recurred/progressed. This is useful in isolating and assessing the effects of our gene of interest on the disease condition. The PFS_MONTHS indicates at what time the disease progressed/recurred or the last time the patient was seen. Disease-specific (DS) survival curves (also known as cause-specific survival) utilize death from the disease of interest as the endpoint. We also performed a similar analysis on genes interacting with IRAK1 in the magenta module as well as for the 10 identified inflammatory genes.

## Results

### Batch effect correction analysis of PRAD dataset in preparation for WGCNA

Genomic and clinical trait data were downloaded from cBioPortal and TCGA databases (**Table 1**), data-mined, and analyzed as highlighted in our bioinformatics pipeline (**Figure 1**). Before WGCNA analysis, a robust preprocessing analysis on the acquired PRAD sample data, including removal of outliers, and detection and removal of low-quality counts using 3 separate methods (TAMPOR, MBatch, and Empirical Bayes (ComBat)) was performed (**Supplementary Figure S1**). As stated in the methods section, these batch effect correction analyses ensured that our final input dataset is devoid of any factors that could skew the outcome of our downstream analyses. Following batch effect correction, we were able to refine our initial data to 472 patients from 494 patients, and 17,794 genes from an initial total of 20,506.

**Table 1:** Distribution of some of the reported clinical traits among prostate adenocarcinoma (PRAD, n = 494) patients.

| Clinical Traits | Classification | Number of Patients/Samples |
| --- | --- | --- |
| Race | Caucasian-American | 147 |
|  | African-American | 7 |
|  | Asian-American | 2 |
|  | NA | 338 |
| Gender | Male | 494 |
| Progression-free Survival (PFS) | Censored | 401 |
|  | Progression | 93 |
| Disease-free Status (DFS) or Relapse-free Status (RFS) | Disease-free | 304 |
|  | Recurred/Progressed | 30 |
|  | NA | 160 |
| Biochemical Recurrence | No | 355 |
|  | Yes | 81 |
|  | NA | 58 |
| Overall Survival (OS) or Vital Status | Living | 484 |
|  | Deceased | 10 |
| Resistance to Radiotherapy | No | 389 |
|  | Yes | 59 |
|  | NA | 46 |
| Cancer Status | Tumor-free | 314 |
|  | With tumor | 94 |
|  | NA | 86 |

NA: Not available

### Module eigengenes and gene relationships in PRAD dataset

WGCNA organizes genes with similar characteristics and expression patterns and associates them with clinical traits of interest. The correlation of the clustered genes with the clinical traits helps to identify modules containing hub genes of clinical significance. Each gene-containing module was correlated with specific clinical outcomes such as progress-free survival (PFS) status, disease-free survival (DFS) status, cancer status, cancer biochemical recurrence, TNM staging pathology, race, gender, radioresistance status, and overall survival (OS) status obtained from TCGA and cBioPortal databases (**Table 1**).

For a better understanding of the genomic and transcriptomic landscape of diagnosed PRAD patients that progressed after initial treatment, we analyzed our data using the WGCNA algorithm in R with some modifications as highlighted in the methods section. The transcriptomes comprising 17,794 gene products across 472 PRAD samples were examined for systems-level relationships by determining co-expression subnetworks (modules) of gene transcripts and gene connectivity (or module membership) using WGCNA with transcript adjacency based on biweight correlation, as described in the methods section.

We generated a similarity co-expression matrix, adjacency matrix (signed), topological overlap matrix (TOM), and dissimilarity matrix (1-TOM) using WGCNA. In total, 9 modules, excluding the junk (grey) module were identified and the module eigengenes were calculated and computed (kMEs, numbered by their size rank from largest to smallest) (**Supplementary Figures S2 – S3**). The grey (M0) module contains junk (not significantly correlated) genes (n = 4739). The other modules, and the number of genes, were turquoise (M1; n = 3245), blue (M2; n=2245), brown (M3; n = 1851), yellow (M4; n = 1469), green (M5; n=1340), red (M6; n = 895), black (M7; n = 825), pink (M8; n=597), and magenta (M9; n = 588) (**Table 2**). After the kMEs were identified, their relatedness was determined using a correlation-based relatedness dendrogram (**Figure 2A**). The relatedness dendrogram showed the proximity clustering between M9 and M1 modules, M3 and M4 modules, and M2 and M8 modules. The relationship between each module and each clinical trait was assessed using the biweight midcorrelation method (**Figure 2B**).

**Figure 2:**
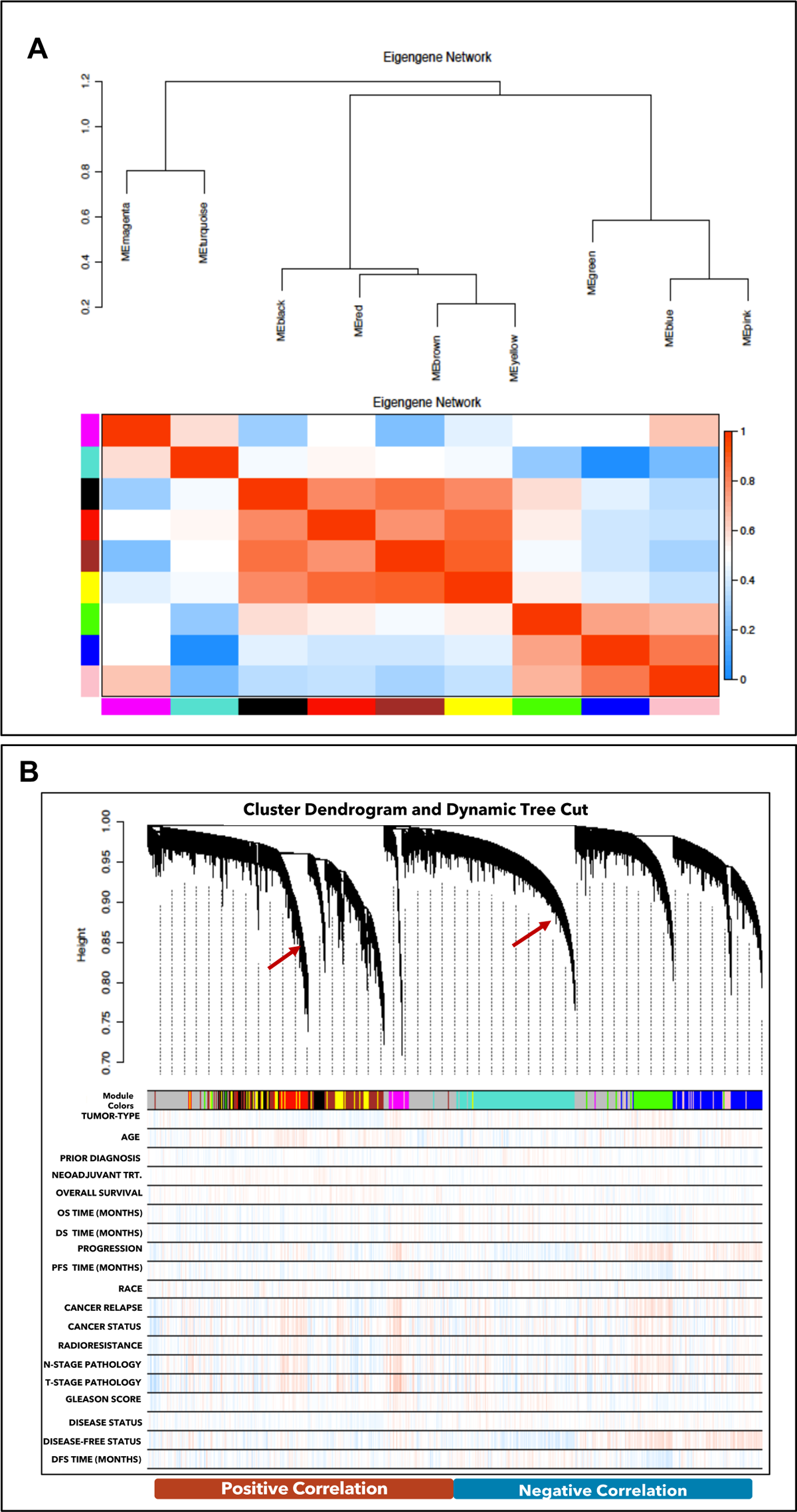
Gene dendrogram of clustered dissimilarity, based on consensus topological overlap, with the corresponding module colors and correlation. **A.** Relatedness dendrogram and correlation heatmap of modules identified by weighted gene co-expression network analysis (WGCNA) showing the dendrogram of consensus module eigengenes obtained by WGCNA on the consensus correlation. **B.** Heatmap plot of the pairwise correlations (adjacency matrix) of module eigengenes. Red boxes represent positive pairwise correlation, and blue boxes represent anti-correlation or negative correlation while the white boxes represent the non-pairwise correlation. Each colored row represents a color-coded module, which contains a group of highly connected genes. A total of 9 modules (excluding grey modules) were identified. The relationship between each relevant clinical trait was evaluated for each color-coded module. Biweight mid-correlation (bicor) was used as a robust alternative to the Pearson correlation coefficient. The red arrows are pointing to branches of the Dynamic Tree Cut which uses a non-constant height cut-off to detect clusters in a dendrogram based on connectedness and correlation coefficient. This plot was generated based on the cutreeHybrid code included in our WGCNA script.

**Table 2:** WGCNA modules output and the corresponding number of genes per module. Table showing the module colors and IDs as well as the number of gene clusters present in each module out of a total of 17, 794 processed genes. 472 PRAD patients/samples analyzed after batch corrections, normalization, and log_2_ transformation. IRAK1, IRAK2, IRAK3, & IRAK4 were located in the magenta, red, brown, and grey (junk) modules, respectively.

|  | **Module Name** | **Module Number** | **Gene Cluster per Module**<br>**Total gene # = 17,794** | **Modules for IRAK Genes** |
| --- | --- | --- | --- | --- |
|  | Magenta | M9 | 588 | IRAK1 |
|  | Green | M5 | 1340 |  |
|  | Red | M6 | 895 | IRAK2 |
|  | Brown | M3 | 1851 | IRAK3 |
|  | Yellow | M4 | 1469 |  |
|  | Blue | M2 | 2245 |  |
|  | Pink | M8 | 597 |  |
|  | Turquoise | M1 | 3245 |  |
|  | Black | M7 | 825 |  |
|  | Grey | Junk Genes | 4739 | IRAK4 |

### Significant modules and co-expressed hub genes positively correlated with PCa progression

The WGCNA generated 9 (color-coded) modules containing various significantly upregulated and downregulated genes. To identify inflammation-associated hub genes associated with PCa progression, we categorized modules based on enriched genes and correlated each module with clinical traits (Module-trait relationship). The consensus kME (Eigengene-based connectivities) for all genes were calculated per their respective modules. Significant highly connected (hub) genes (kME > 3.0) were identified in each module based on their module-dependent connectedness. The magenta (p = 4.0e-6, r = 0.21), green (p = 2.0e-5, r = 0.19), blue (p = 0.02, r = 0.11) and pink (p = 0.03, r = 0.099) modules were found to be positively correlated with PCa progression while only the brown module was negatively correlated (p = 0.02, r= −0.11) with progression (**Figure 3**). Genes were isolated from the most statistically significant and highly correlated module (Magenta) and associated with our clinical trait of interest (progression) (**Table 2**).

**Figure 3:**
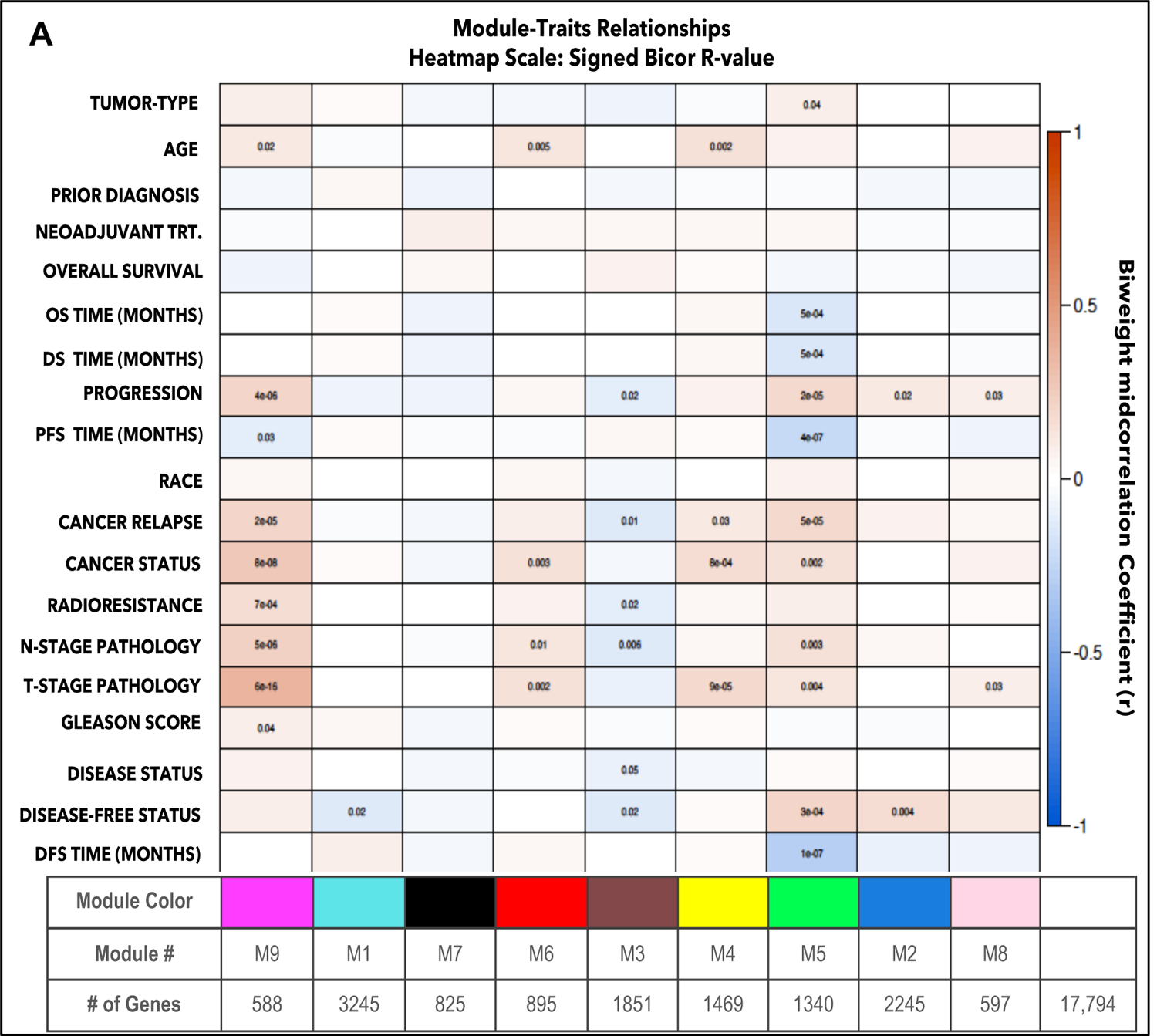
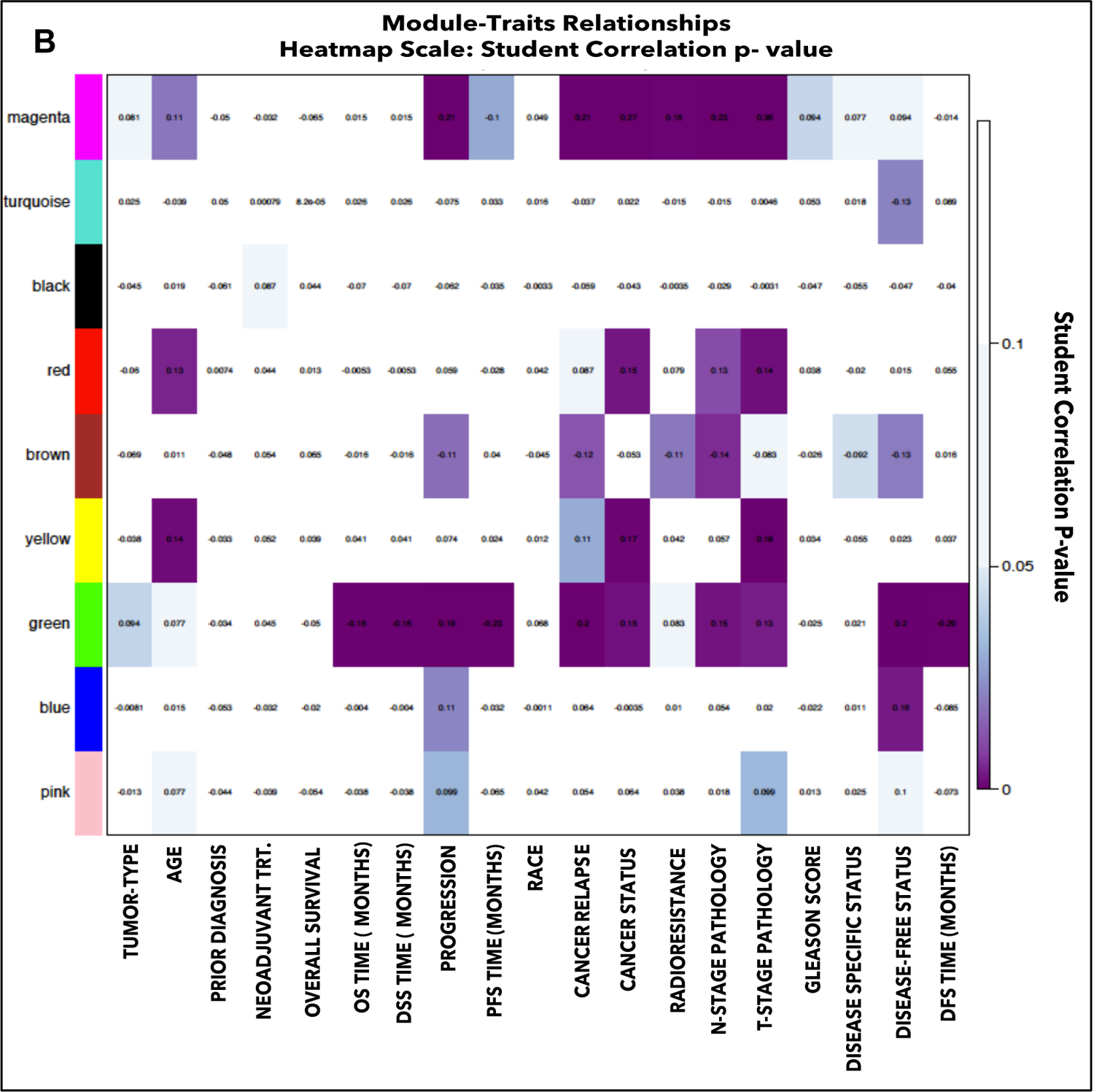
Module-trait relationship based on p-value and bicor correlation coefficient to reveal modules associated with PRAD clinical traits. **A.** showing the heatmap of module-trait relationships based on the bicor (r) value. Each row in the heatmap corresponds to a specific clinical trait and each column to a module. The module colors are shown on the lower side of each column. Boxes colored blue and red signify negative and positive correlations to PRAD clinical traits or phenotypes, respectively. Modules with significant Module-trait biweight correlation (bicor) color scale (−1 = blue; 0 = white; +1 = red) were indicated by including the significant p-value in the boxes. **B.** showing the heatmap of the module-trait relationship based on the p-value significance. Each column corresponds to specific clinical traits and each row to a module. The module colors are shown on the left side of each row. The color gradient of the heatmap boxes from purple to white denotes the correlation p-value with significance from 0 to 0.05 as well as no significance from 0.05 to upward.

The magenta module has 588 genes in it (**Supplementary Figure S11**) and about 326 of them were found to be significantly upregulated and positively correlated with the progression status of PRAD patients (**Figure 3**). Overall, magenta (M9) contains genes that have positive correlation with cancer progression (bicor r = 0.21, p = 4e-06), age (bicor r = 0.11, p = 0.02), cancer relapse (bicor r = 0.21, p = 2e-05), lymph node (N-stage) pathology (bicor r = 0.23, p = 5e-06), tumor (T-stage) pathology (bicor r = 0.36, p = 5e-16), and Gleason score ((bicor r = 0.094, p = 0.04) but negatively correlated with period (months) of progression-free status (bicor r = 0.1, p = 0.03). The kME values for the IRAK family members were also identified per module: IRAK1 (magenta; kME = 0.394), IRAK2 (red: kME = 0.5112), IRAK3 (brown: kME = 0.8043), and IRAK4 (grey: kME = 0.0995).

### Association of IRAK family genes with PRAD progression

Because the IRAK family is made up of 4 genes, we were interested to know if the other members of this family also contribute to PRAD progression (**Table 2**). We first located the modules that the others are located and calculated their kME values as well as their biweight (bicor r) correlation with progression. There was no significant difference (p-value = 0.485) in IRAK1 expression between progressed and non-progressed patients (F-value: 0.488; FDR (BH): 0.662; FC: −0.064). Also, IRAK4 was not significantly associated (p-value = 0.285) with the progression status of PRAD patients (F-value: 1.147; FDR (BH): 0.479; FC: 0.039). Interestingly, IRAK3 was found to be significantly (p-value = 0.038) and negatively correlated with the progression status of PRAD patients (F-value: 4.351; FDR (BH): 0.133; FC: 0.223). This indicates that the downregulation of IRAK3 in PRAD patients may favor PRAD progression.

### Differential gene expression analysis between progressed and non-progressed PRAD patients per module

Differentially expressed genes in all modules including grey modules for all PRAD patients that progressed versus those that did not progress were identified using the ANOVA-Turkey statistical method. There were 3230 significantly upregulated and 2342 downregulated genes in total with fold change (the difference between progressed and non-progressed) ranges of −1.58 to 1.4. Of significance, the volcano plots of -log_10_ p-value vs. log_2_ fold change (FC) of the PRAD progression status showed that magenta (M9) and green (M5) have 326 and 813 significantly upregulated genes and both contain no significantly downregulated genes. The blue module has 813 upregulated and 33 downregulated genes, pink has 192 upregulated genes and 6 downregulated genes while yellow has 328 upregulated and 12 downregulated genes. The red module has 157 upregulated and 14 downregulated genes while the turquoise module has 50 upregulated and 898 downregulated genes. The brown module has 3 upregulated and 647 downregulated genes while the black module has 16 upregulated and 146 downregulated genes. Lastly, the grey has 477 upregulated and 586 downregulated genes (**Figure 4**).

**Figure 4:**
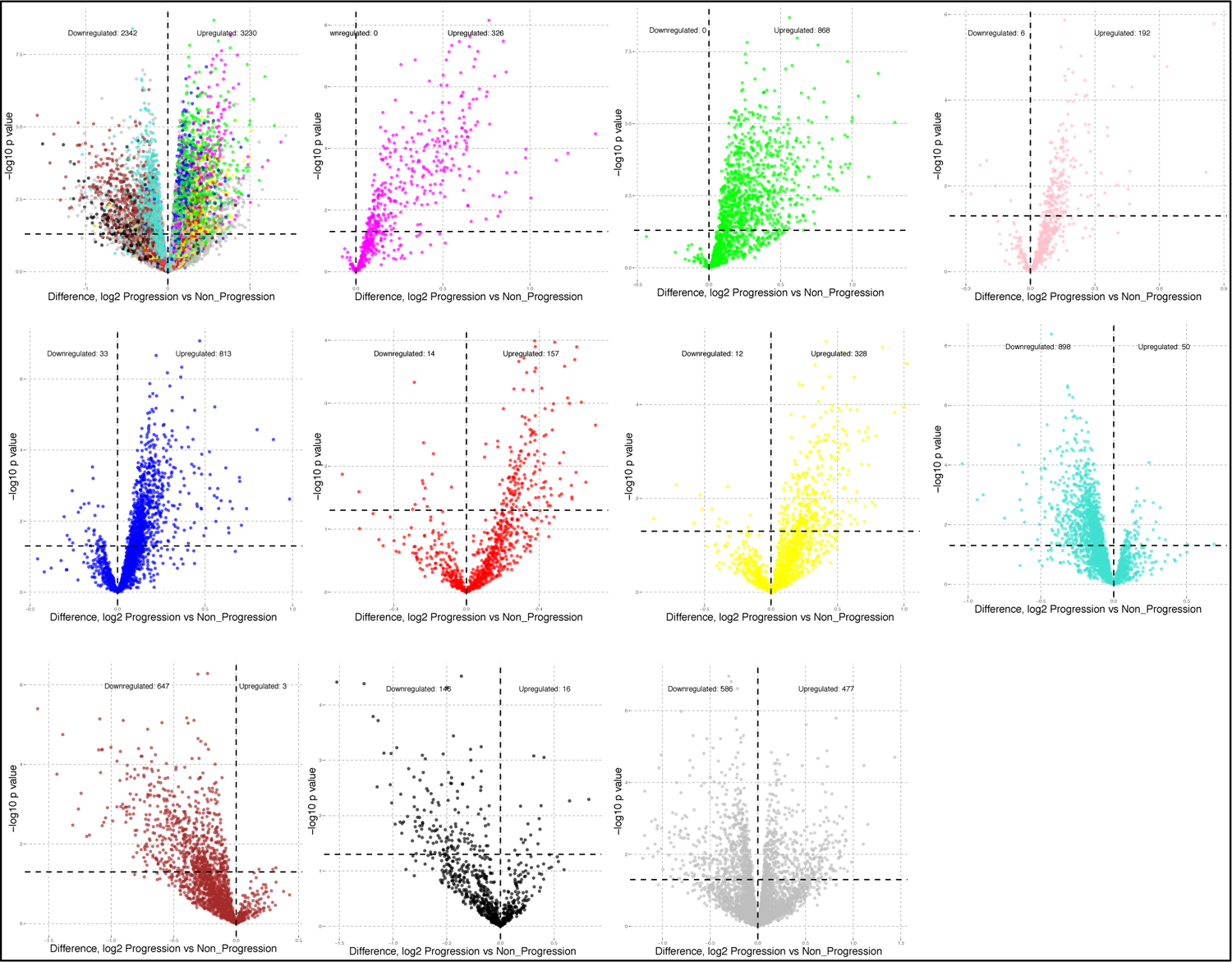
Volcano plots (-log_10_ p-value vs log_2_ fold change) of differentially expressed genes between progressed vs non-progressed PRAD patients in each module. Differential gene expression was performed to compare PRAD patients who progressed (n = 93) with those that do not have progression (n = 401). The first volcano plot on the left is showing the DGE of all genes (n = 17,794) located in all the modules together while others are plots for each module. Magenta, green, pink, blue, red, yellow modules appear to have mostly upregulated genes correlated to progression in PRAD patients while turquoise, brown and black modules are mostly made up of downregulated genes in correlation with progression in PRAD patients. The p-values were computed using ANOVA-Tukey statistical test. On the y-axis of each volcano, the plot is the -log10 of p-value while on the x-axis is the log_2_ of FC. The horizontal dotted lines show the line of significance at p-value < 0.05.

### Functional enrichment analyses of inflammatory genes in the magenta module

Go-Elite was used to analyze and identify the functional annotation and gene ontology for each module based on their co-expression, relatedness, and kME scores (**Figures 5 and** Supplementary **Figure S16**). Based on the significant association of the magenta module with PRAD progression in the WGCNA analysis, we decided to focus on the genes in this module. We used WebGestalt to perform gene set enrichment analysis (GSEA) as well as over-representation analysis (ORA) for the magenta (M9) module which categorized the genes in this module based on their biological processes, cellular components, and molecular functions. Many of these genes in the magenta were found to be associated with cellular processes such as metabolism and biological regulation of cell communication, cell cycle, cell division, cell proliferation, cell reproduction, cell growth, cell response to stimuli, and many others. A greater percentage of the magenta genes were also found to be localized in the nucleus and cytosol. Some of the identified molecular functions of these genes include protein binding, nucleic acid binding, ion binding, enzyme regulator activity, and many others (**Figure 6**).

**Figure 5:**
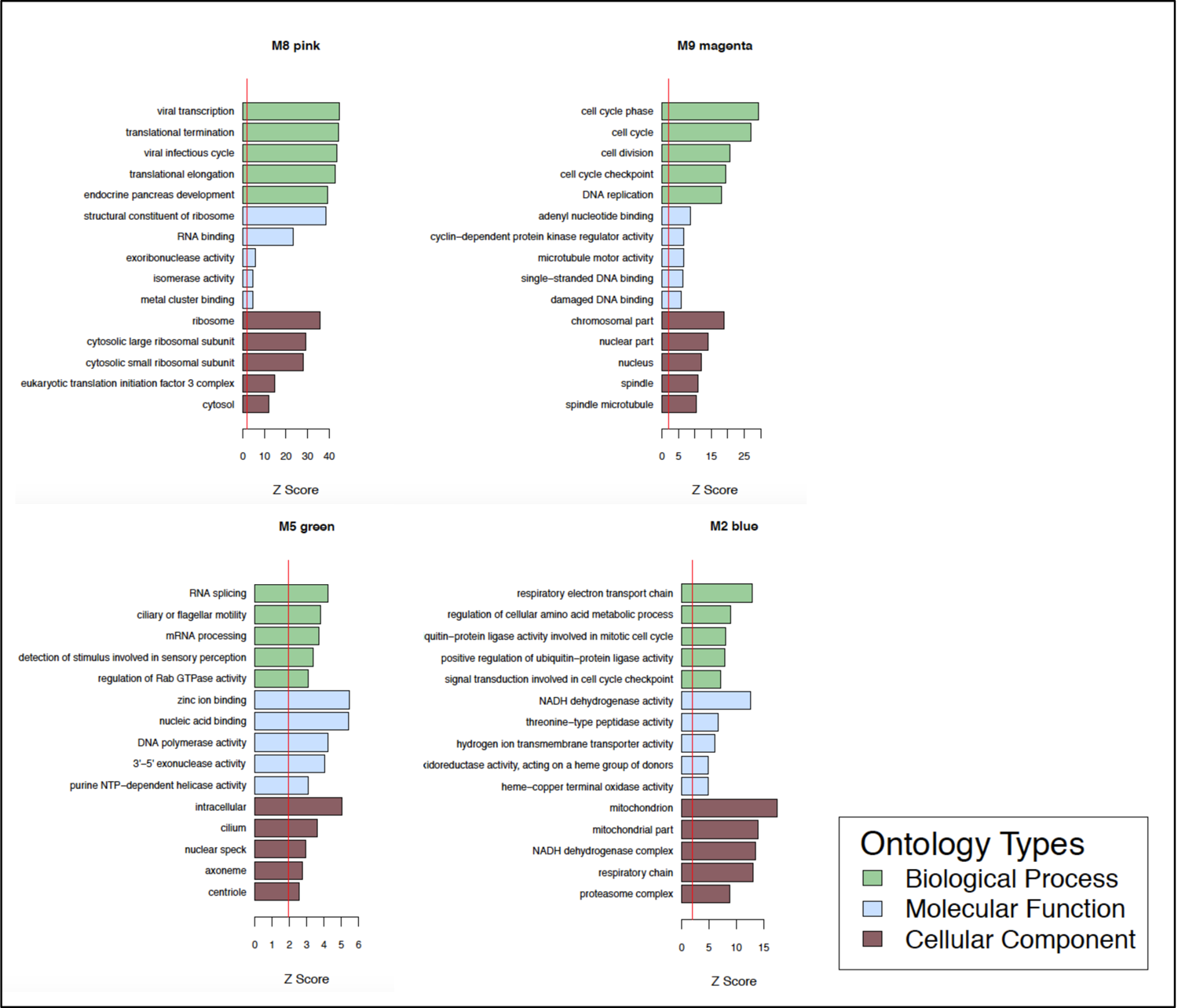
GO Elite ontologies of the four biologically significant modules positively correlated with PRAD progression. Gene ontology of the magenta, pink, green, and blue modules was determined based on their biological processes, molecular functions, and cellular components. The x-axis represents the calculated Z-scores for the gene ontologies. The y axis has the biological processes (green bars), the molecular functions (blue bars), and the cellular components for each module (brown bars). The red vertical lines shown across the bars represent the direction of the standard deviations.

**Figure 6:**
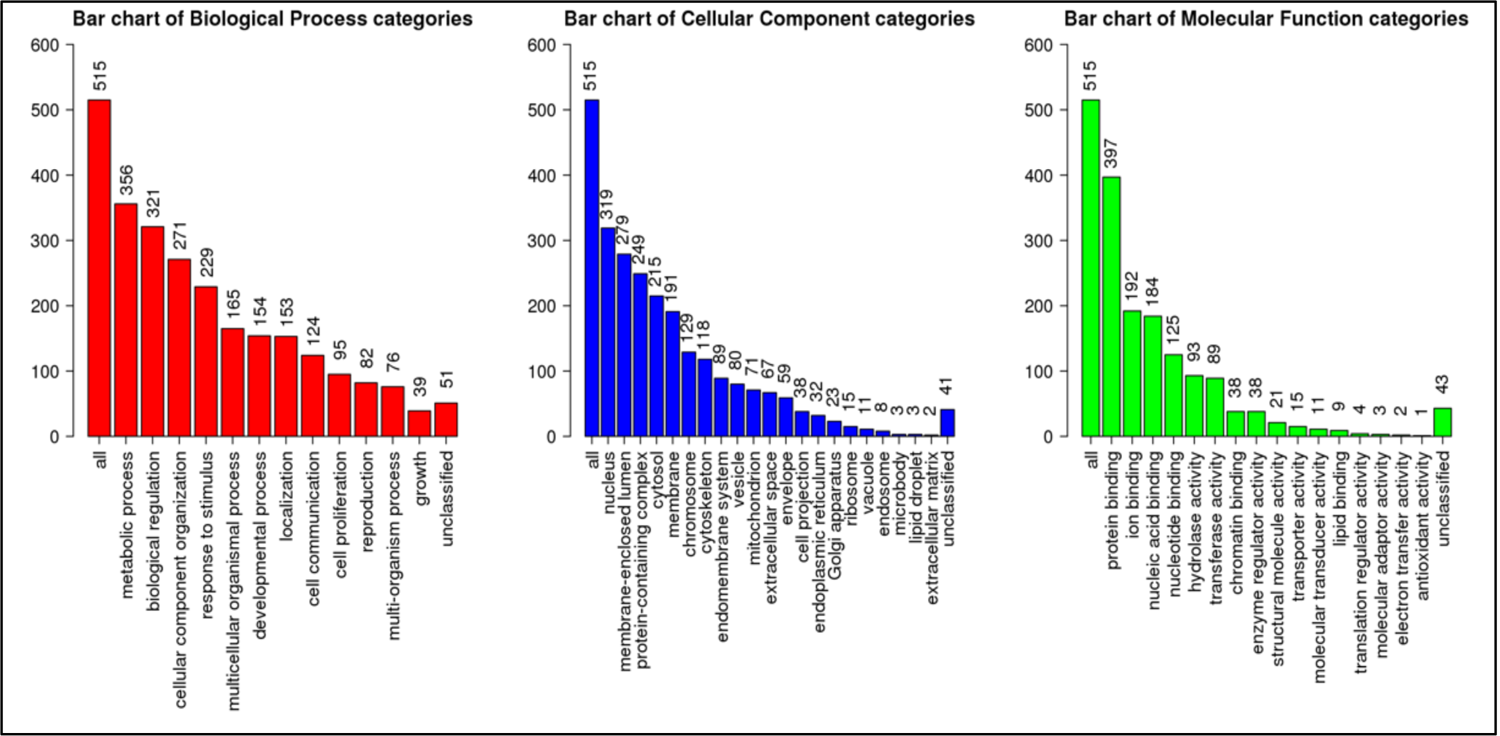
WebGestalt gene-enrichment analysis and biological ontologies for the magenta module. The x-axis represents the gene ontologies predicted based on their biological processes (red bar chart), molecular functions (green bar chart), and cellular components (blue bars). The y-axis represents the number of genes.

### Predicting protein-protein interaction and differential gene expression analyses of inflammatory genes between progressed and non-progressed PRAD samples

The inflammatory gene sets for each module were identified and computed from KEGG, DAVID, and GeneCards. Enrichr and Gene Ontology Consortium were used to validate, curate, filter, and categorize gene sets into high, middle, and low priority groups based on their association with the inflammatory signaling cascade. IRAK1 (kME = 0.394) was identified as one of the 9 inflammatory significant genes in the magenta module. The other 9 inflammatory genes present in the magenta module are TRIM59 (kME = 0.5975), TRAIP (kME = 0.5849), NFKBIL2/TONSL (kME = 0.5749), HMGB3 (kME = 0.4571), TRAF7 (kME = 0.4568), PPIL5/LRR1 (kME = 0.4513), ILF2 (kME = 0.4256), HMGB2 (kME = 0.4184), and IL1F5/IL36RN (kME = 0.3707). Enrichr was mined to generate diverse functional ontologies for the 10 inflammatory genes, categorized based on biological processes, cellular components, molecular functions, and enriched pathways.

About 50 of the 150 biological functions associated with these genes were found to be statistically significant (p < 0.05) and related to inflammatory processes. HMGB2 and IRAK1 were the most biologically enriched among the 10 genes. However, IRAK1 was the most associated with inflammation (**Supplementary Table S1**). Furthermore, protein-protein interaction (PPI) and enrichment (PPE) tools from Reactome and String-db. (v11) were used to predict and visualize the interaction networks between the inflammatory genes/proteins of interest and other genes within the inflammatory pathway. To further determine if these genes were upregulated and differentially expressed genes between patients with PRAD progression and those that did not progress, we used ANOVA at p < 0.05 to test our hypothesis and calculated the F-value, false discovery rate (FDR), and fold change (FC) between the two groups. The ANOVA-TURKEY test output identified the IRAK1 gene to have a FC = 0.119; p-value = 0.026; F-value = 4.968; FDR = 0.106. The ANOVA output for the enriched inflammatory genes is listed in **Table 3**.

**Table 3:** WebGestalt gene-enrichment analysis and biological ontologies for the magenta module. The table is showing the calculated F-value, Benjamini and Hochberg false discovery rate (FDR), ANOVA p-value = 0.05, and the fold change (FC, the difference in mean gene expression values between progressed vs non-progressed patients).

| Gene | F-Value | FDR | P-value | FC |
| --- | --- | --- | --- | --- |
| IRAK1 | 4.96776529 | 0.10641623 | 0.02629606 | 0.11896523 |
| PPIL5/LRR1 | 8.54238016 | 0.03089667 | 0.00363703 | 0.12637696 |
| HMGB3 | 4.46690454 | 0.12773763 | 0.03508388 | 0.13009469 |
| HMGB2 | 7.87684581 | 0.03820747 | 0.00521516 | 0.19869876 |
| TRAIP | 20.6970149 | 0.00086135 | 6.86e-06 | 0.31806626 |
| IL1F5/IL36RN | 8.32987379 | 0.03303962 | 0.00408493 | 1.0060423 |
| ILF2 | 2.63225541 | 0.25575456 | 0.10538342 | 0.05404065 |
| TRIM59 | 3.76977041 | 0.1651797 | 0.05278405 | 0.13270392 |
| NFKBIL2/TONSL | 10.5579031 | 0.01629371 | 0.00123986 | 0.25775675 |
| TRAF7 | 3.73438085 | 0.16748158 | 0.05390396 | 0.08140051 |

### Correlogram and differential gene expression (DGE) analysis of inflammatory genes between PRAD and non-PRAD normal samples

After identifying the 10 inflammatory genes in the magenta module, we conducted a DGE analysis using the DESeq package in R studio interface to identify their differential expression patterns (fold change) relative to non-malignant normal samples. The RNAseq expression values of 52 matched prostate tumors and adjacent normal samples were compared for the matched/paired analysis while the RNAseq of 492 tumors and 204 normal samples from non-cancerous patients were compared for the unmatched/unpaired analysis (**Figure 7**). A non-parametric test, Wilcoxon Rank Sum Test (Mann-Whitney U test) was used to determine the fold change (FC) and significant difference (p < 0.05) in gene expression levels between the tumor and normal samples for both the paired and unpaired analyses (**Table 4**).

**Figure 7:**
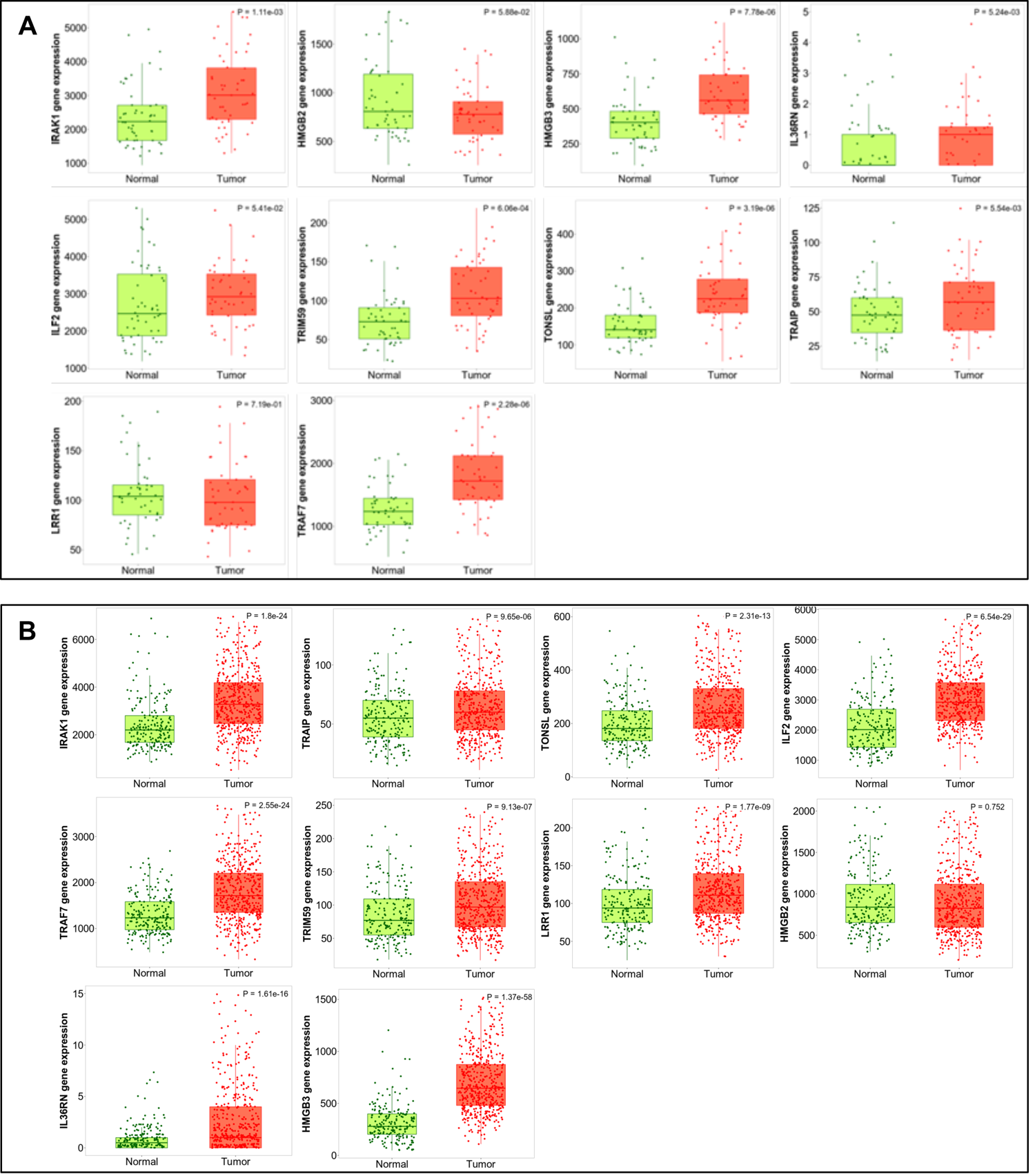
Matched/paired (7A) and unmatched/unpaired (7B) differential gene expression analysis plots. Each whisker boxplot plot shows the gene expression distribution of identified inflammatory genes in the magenta module between prostate tumor vs normal samples.

**Table 4:** Wilcoxon Rank Sum Test (Mann-Whitney U test) output for the inflammatory signature gene set in the magenta module. The table shows the p-values and fold change (FC) in gene expression levels between prostate tumor vs normal samples from the matched and unmatched analyses.

| Samples | Matched/Paired |  | Unmatched/Unpaired |  |
| --- | --- | --- | --- | --- |
| Genes | P-value | FC | P-value | FC |
| IRAK1 | 1.11e-3 | 1.27 | 1.8e-24 | 1.46 |
| PPIL5/LRR1 | 0.719 | 0.95 | 1.77e-9 | 1.25 |
| HMGB3 | 7.78e-6 | 1.58 | 1.37e-58 | 2.23 |
| HMGB2 | 5.88e-2 | 0.88 | 0.752 | 1.07 |
| TRAIP | 5.54e-3 | 1.25 | 9.65e-6 | 1.29 |
| IL1F5/IL36RN | 5.24e-03 | 8.61 | 1.61e-16 | 8.69 |
| ILF2 | 5.42e-2 | 1.21 | 6.54e-29 | 1.5 |
| TRIM59 | 6.06e-4 | 1.36 | 9.13e-7 | 1.29 |
| NFKBIL2/TONSL | 3.18e-6 | 1.54 | 2.31e-13 | 1.47 |
| TRAF7 | 2.28e-7 | 1.48 | 2.55e-24 | 1.46 |

**Table 5:** AUROC statistics for magenta inflammatory genes. Column 1 is the Gene marker and Column 2 is the AUC values. Columns 3 and 4 have the lower and upper AUC limits at 95% CI. Column 5 has the Z-scores, and Column 6 has the Wilcoxon test p-values.

| **Genes Marker** | **AUC** | **Lower Limit**<br>**(95% CI)** | **Upper Limit**<br>**(95% CI)** | **Z-score** | **P-value** |
| --- | --- | --- | --- | --- | --- |
| IRAK1 | 0.58209 | 0.5137 | 0.65047 | 2.35261 | 0.01864 |
| HMGB2 | 0.60169 | 0.53338 | 0.66999 | 2.91778 | 0.00353 |
| HMGB3 | 0.57261 | 0.50423 | 0.64098 | 2.08129 | 0.03741 |
| IL1F5/IL36RN | 0.60415 | 0.53535 | 0.67295 | 2.96707 | 0.00301 |
| ILF2 | 0.56505 | 0.49671 | 0.6334 | 1.86566 | 0.06209 |
| PPIL5/LRR1 | 0.59286 | 0.5245 | 0.66122 | 2.66254 | 0.00776 |
| TRAF7 | 0.56271 | 0.49438 | 0.63104 | 1.79882 | 0.07205 |
| TRAIP | 0.64193 | 0.57424 | 0.70962 | 4.10972 | 4.00e-05 |
| NFKBIL2/TONSL | 0.61821 | 0.55008 | 0.68634 | 3.40083 | 0.00067 |
| TRIM59 | 0.57125 | 0.50288 | 0.63962 | 2.04248 | 0.0411 |

**Table 6:** Identification of 13 DNA methylation sites in IRAK1, located on Chromosome X. 13 CpG probes were identified for our analysis based on the CpG regions in the IRAK1 transcript. The Shore indicates 0 to 2kb from the CpG island, including N_Shore (0-2 kb upstream from CGI) and S_Shore (0-2 kb downstream from CGI). While the Shelf indicates 2-4 kb from CpG island, including N_Shelf (2-4 kb upstream from CGI) and S_Shelf (2-4 kb downstream from CGI). Open Sea: Isolated CpGs in the genome.

| CpG Probe | Chromosome | Start | End | CGI position | Strand |
| --- | --- | --- | --- | --- | --- |
| cg08401365 | chrX | 154011509 | 154011510 | Open Sea | + |
| cg20494209 | chrX | 154016771 | 154016772 | N_Shelf | - |
| cg23604959 | chrX | 154018243 | 154018244 | N_Shore | + |
| cg02742918 | chrX | 154018652 | 154018653 | N_Shore | - |
| cg06334238 | chrX | 154019448 | 154019449 | Island | + |
| cg18998000 | chrX | 154019508 | 154019509 | Island | + |
| cg23121114 | chrX | 154019667 | 154019668 | Island | - |
| cg01353347 | chrX | 154019707 | 154019708 | Island | - |
| cg09520212 | chrX | 154020171 | 154020172 | Island | - |
| cg27167979 | chrX | 154020194 | 154020195 | Island | - |
| cg03050491 | chrX | 154020203 | 154020204 | Island | - |
| cg19572242 | chrX | 154020339 | 154020340 | S_Shore | + |
| cg27616996 | chrX | 154020483 | 154020484 | S_Shore | - |

All of the 10 inflammatory genes showed upregulation in the prostate tumor samples relative to the normal samples. HMGB2 has the lowest FC (0.88 and 1.07) while IL1F5/IL36RN has the highest FC (8.61 and 8.69) for both the matched and unmatched analyses, respectively. No significant difference (p > 0.05) between the PRAD and normal samples was observed for PPIL5/LRR1 and HMGB2 in the paired and unpaired analyses. Also, correlogram analysis between the 10 inflammatory genes in the magenta module reveals both positive and negative associations between the genes (**Figure 8**). IRAK1 differential expression correlates positively with TRAF7 and NFKBIL2/TONSL but negatively correlates with HMGB2 expression, while TRAIP differential expression correlates positively with TRAF7, NFKBIL2, HMGB2, and TRIM59 expression.

**Figure 8:**
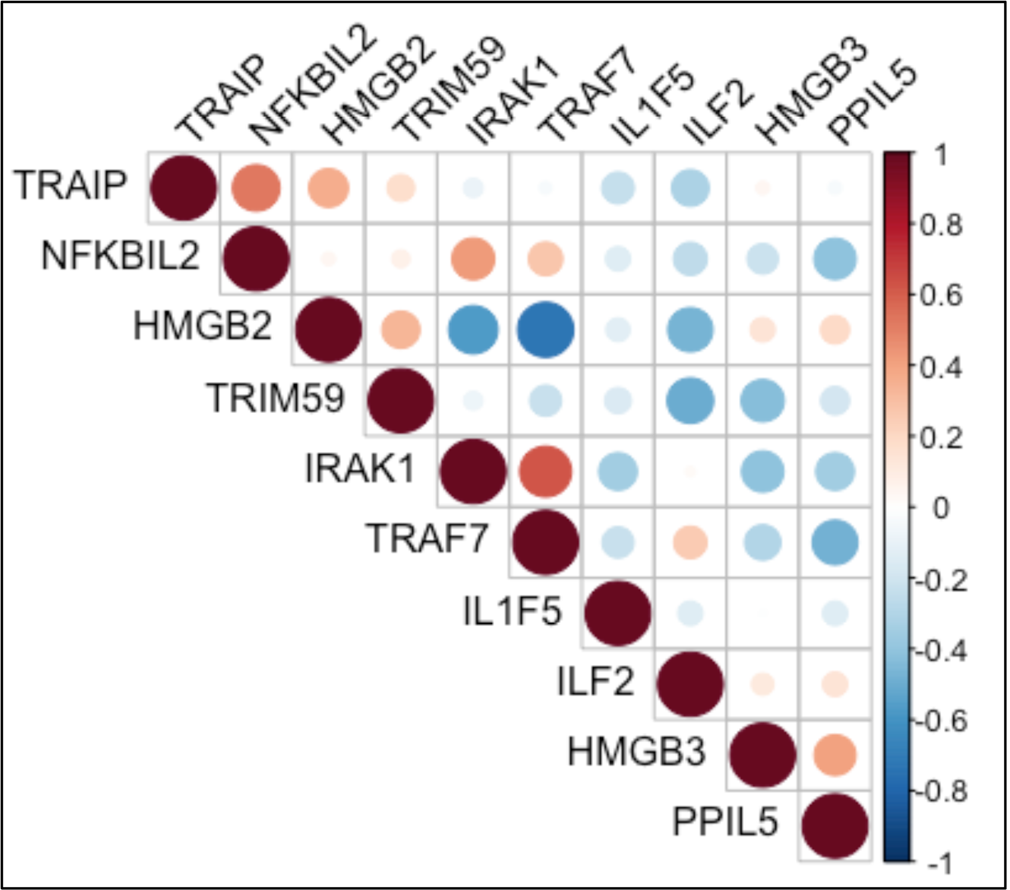
Correlogram plot showing the correlation between the 10 inflammatory genes in the magenta module. The positively and negatively correlated genes are shown in red and blue colors, respectively.

### Prognostic significance of IRAK1 overexpression in PRAD patients

To establish the prognostic and clinical significance of IRAK1, we correlated the expression RNAseq dataset and clinical data on progress-free survival for 472 PRAD patients. The expression levels of the IRAK1 gene were divided into two groups, IRAK1 high-expressing patients (n = 236) vs low-expressing patients (n = 236) using the median expression cut-off value. The two groups were compared by a Kaplan-Meier survival plot. The Kaplan-Meier survival analysis estimated and plotted the survival curves, including the overall survival (OS), disease-free survival (DFS), and progression-free survival (PFS). The Cox proportional hazards model calculated the hazard ratio with a 95% confidence interval between the two groups. The log-rank test was used to statistically test the significance (p < 0.05) of our null hypothesis. The number of patients at risk for each time interval in months (i.e., 0, 50, 100, 150 months) was also calculated. While no significance was found for OS using the median cut off, PRAD patients with high expression of IRAK1 were associated with poor outcomes i.e., lower progression-free survival time and lower disease-free survival (**Supplementary Figure S4**) as well as increased risk of progression with a significant log-rank p-value of 0.018 and a hazard ratio (HR) of 1.68 (95% CI: 1.09 – 2.59). The hazard ratio indicates that IRAK1 high-expressing PRAD patients have a 68% (HR = 1.68) higher risk of progression compared to the low-expressing PCa patients (**Figure 9**).

**Figure 9:**
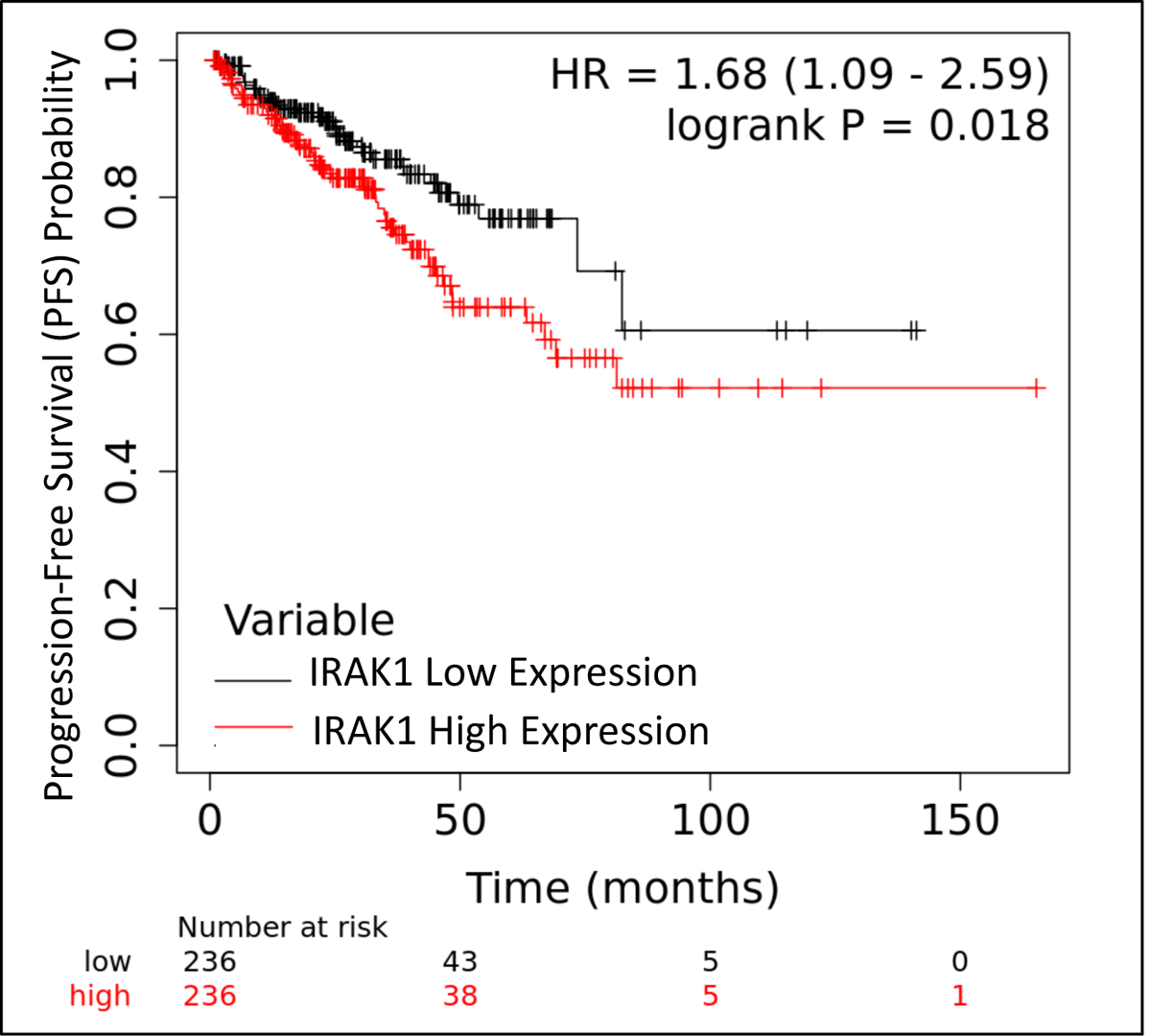
Kaplan-Meier survival analysis of PFS in IRAK1 low-expressing vs high-expressing PRAD patients. Kaplan-Meier survival analysis was performed for the PRAD dataset (n = 472). PRAD patients were stratified into two groups of IRAK1 high-expressing patients (n = 236) vs IRAK1 low-expressing patients (n = 236). Log-rank p-value and Hazard ratio were calculated for progression-free survival using the median expression value for all samples as the cut-off.

**Figure 10:**
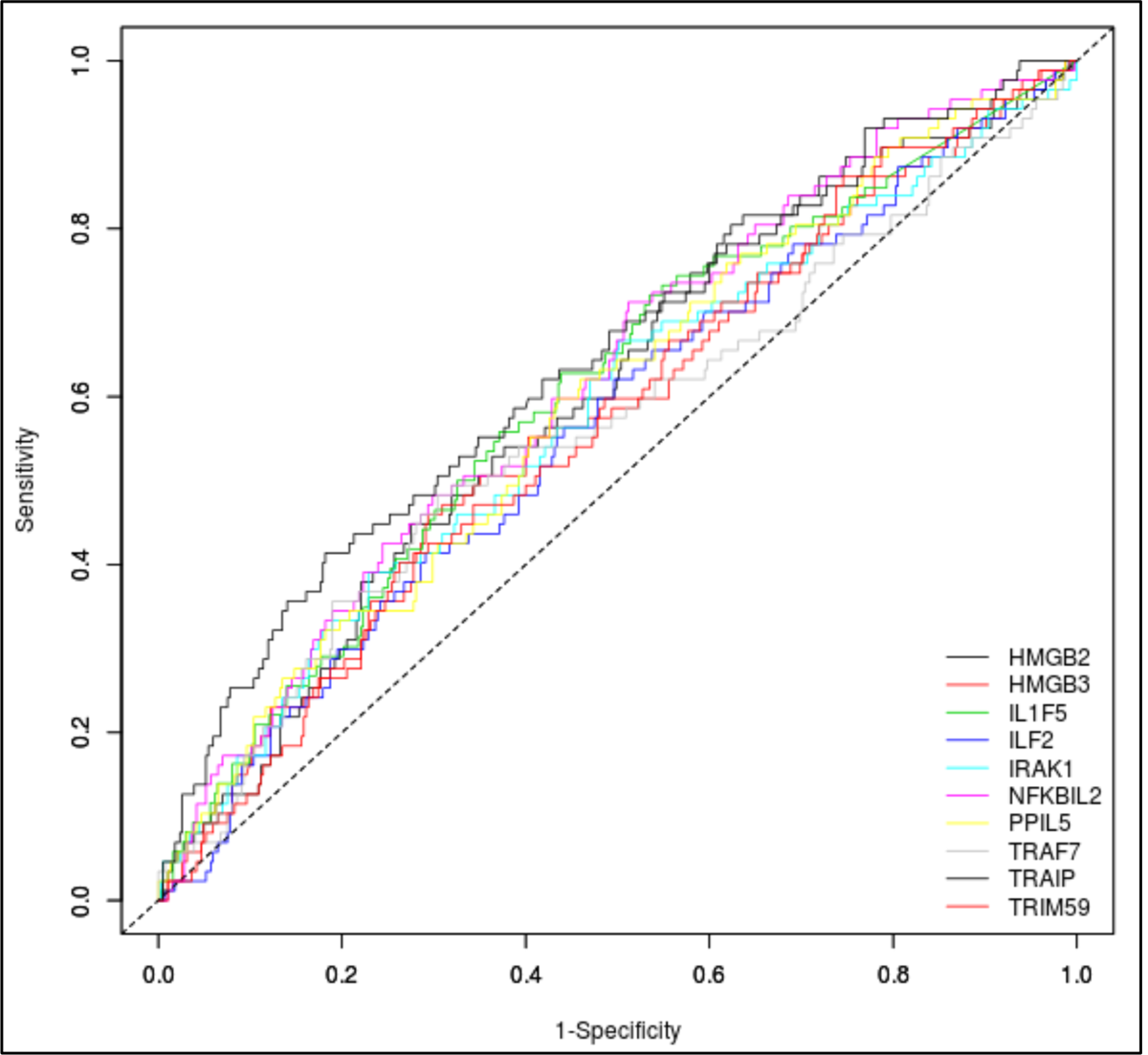
Receiver Operator Characteristic (ROC) analysis for the 10 inflammatory genes. The AUROC plot shows the prediction of 4 of the 10 genes as potential biomarkers for PRAD progression. Y-axis represents the specificity while the x-axis represents the 1-specificity. The black dotted line is the line of significance (p < 0.05). The Area under the Curve (AUC) shows that HMGB2 (AUC=0.6), IL1F5 (AUC=0.6), TRAIP (AUC=0.64), and NFKBIL2 (AUC=0.61) have the potential to predict poor outcome in ∼60% of PRAD patients with a significant p-value at p < 0.05.

**Figure 11:**
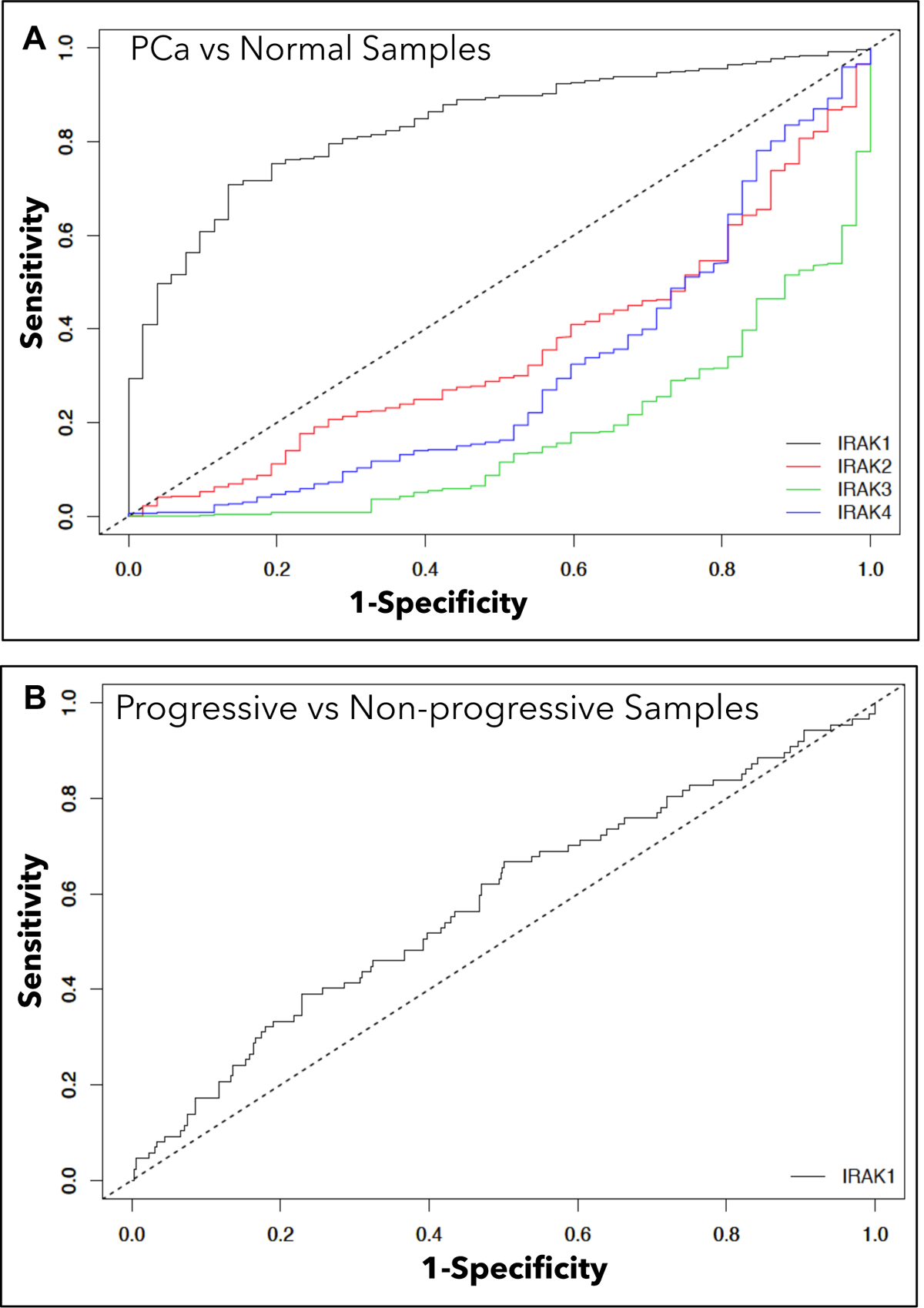
Receiver Operating Characteristic (ROC) analysis for IRAK1 in PRAD vs normal samples and progressive vs non-progressive PRAD samples. This ROC analyzes the capability of IRAK1, a representative of inflammatory genes in the magenta module to predict survival risk. The x-axis represents sensitivity while the y-axis represents 1-specificity. The black dotted lines represent the line of significance (p-value = 0.05). The ROC curves represent the gold standard of diagnostic accuracy (i.e., genes with AUC greater than 0.6 are assumed to be a potentially good diagnostic biomarker). As diagnostic test accuracy improves, the AUC approaches 1. **A.** Denotes the AUROC comparing PCa samples from normal samples. AUC of IRAK1 (0.83) shows that IRAK1 is an excellent diagnostic biomarker compared to other members of the family in PRAD patients, for differentiating between PCa and non-PCa tissue samples. **B.** Denotes the AUROC comparing progressed PRAD patients to non-progressed PRAD patients, with an AUC of 0.58 (58%).

### Protein-protein network and co-expression analyses of genes in the magenta module

Cytoscape software was used to reveal the degree of co-expression gene network interaction for all the 588 genes in the magenta module, with special emphasis on IRAK1 (**Figure 12A**). The identified genes with the closest interaction with IRAK1 include among others, malate dehydrogenase (MDH2), importin-4 (IPO4), and mannosyl (alpha-1,3-)-glycoprotein beta-1,4-N acetylglucosaminyltransferase, isozyme B (MGAT4B). Some of these genes have been studied and shown to have direct or indirect relevance in carcinogenesis. For instance, MDH2 catalyzes the reversible oxidation of malate to oxaloacetate, which is important in the malate-aspartate shuttle and ATP production in the TCA cycle. Overexpression of MDH2 has been implicated in PCa resistance to docetaxel-chemotherapy, doxorubicin-resistant uterine cancer cells, and P-glycoprotein-induced multi-drug resistance (MDR) by pumping chemotherapeutic drugs out of the cells (Liu et al., 2013). Downregulation of MDH2 has been shown to increase drug sensitivity and appears to be a potential therapeutic target to enhance the efficacy of anticancer drugs. IPO4 has been suggested to contribute to gastric cancer progression and poor prognosis (Xu et al., 2019). Inhibition of IPO4-mediated nuclear import of CEBPD has also been shown to enhance chemosensitivity by repression of PRKDC-driven DNA damage repair in cervical cancer (Zhou et al., 2020). The role of IRAK1 in IPO4 deregulation and signaling has not been studied, either in PCa or cancer-associated inflammation. MGAT4B has been shown to play a critical role in N-glycan branching on the surface of tumor cells and therefore important in tumorigenesis directly or indirectly (Ashkani and Naidoo, 2016).

To further understand the biological topography of IRAK1 to prostate carcinogenesis, protein-protein interaction (PPI) and enrichment (PPE) tools from Reactome and String-db (v11) were used to rank, analyze, and visualize the interaction networks between IRAK1 and proteins that have been associated with cancer stemness, cell survival, neuroendocrine differentiation, and chronic inflammation in PCa (**Figure 12B**). The PPI and E analyses were able to predict new biological functions for IRAK1 based on direct or indirect interactions by varying confidence levels, evidence-based criteria, and mode of action (**Supplementary Figure S10**). PPE analysis predicted biological functions such as cell death, cell communication, cell differentiation, cell migration, cell cycle, regulation of response to stress, cell proliferation, and stress response, for IRAK1. Indeed, some of these biological functions have been observed in other tumors that overexpress IRAK1, which further justifies our interest in IRAK1 and its role in PCa progression. Our PPI analysis predicted a posttranslational modification of IRAK1 by AKT1, IRAK2, and IRAK4 as well as uncharacterized effects between PI3K/AKT genes and IRAK1. Also, IRAK1 could directly bind to TLR4, TLR8, TLR9, MYD88, IRAK2, IRAK3, and IRAK4 (**Supplementary Figure S10**). The predicted interactions between IRAK1 and these genes/proteins include gene neighborhood, gene fusions, gene co-occurrence, gene co-expression, and protein homology.

### Clinical significance of the nearest co-expressed neighbors of IRAK1 in the magenta module

Cytoscape identified novel genes interacting with IRAK1 within the magenta module. Some of these genes have been reported in the literature to drive tumor progression while the oncogenic effects of others are unknown. To establish if they have any clinical significance in PCa progression, we performed a threshold analysis using AUROC with the line of significance set at 0.05 (**Figure 13**). The AUC result and plot showed that only one of the three genes directly interacting with IRAK1 could significantly (p < 0.05) predict poor outcomes in PRAD patients (**Figure 13**). MGAT4B gene is one of the nearest neighbors of IRAK1 in the magenta module and has the potential to be used as a prognostic biomarker for PRAD progression (AUC of 0.685, 95% CI: 0.6243 - 0.7458, Z-score: 5.97144). Our AUROC result indicates that MGAT4B can predict progression in 69% of PRAD patients. The other close neighbors to IRAK1 such as MDH2 and IPO4 do not have significant p-values and are therefore poor biomarkers for PRAD progression. Both IPO4 and MDH2 have a similar AUC value of 0.54 (i.e., 54% predictability of PRAD progression).

**Figure 12:**
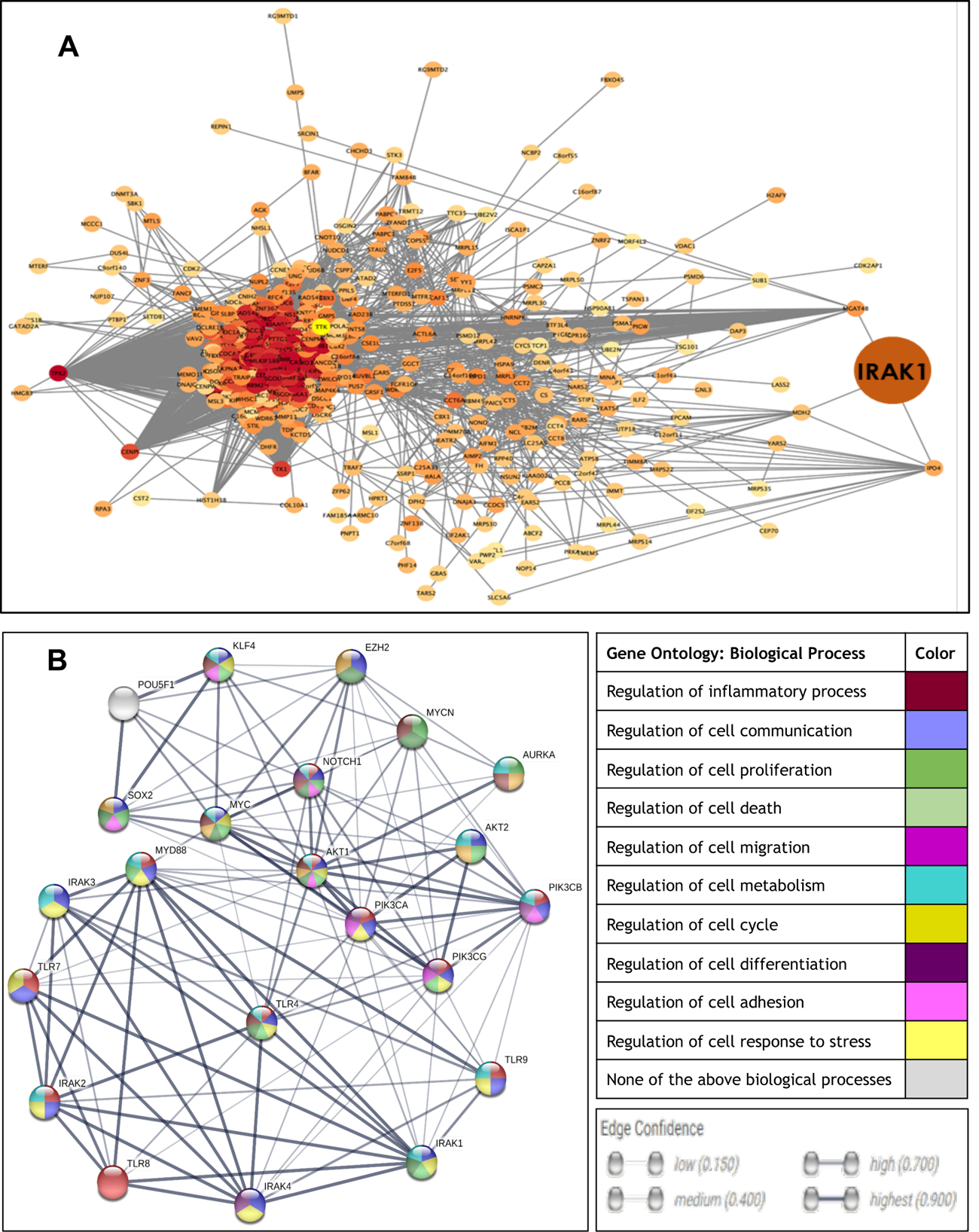
Protein-protein enrichment and network analyses of the PRAD dataset. **A.** Cytoscape network analysis showing the molecular network interaction of magenta module-hub genes. The deeper the color of nodes, the more the known significance of the gene in PCa progression. IRAK1 was found to closely interact with IPO4, MDH2, and MGAT4B. Cytoscape analysis showed that these genes have protein to protein interactions with PCa-associated genes. This is the first report to show such interaction. **B.** Protein-protein interaction (PPI) and enrichment (PPE) network analyses of IRAK1 and selected inflammation and PCa-progression proteins. showing known and predicted interaction between selected IRAK proteins and PCa progression-associated proteins. Pathway interactions were determined using evidence-based functional annotations/gene ontology and edge confidence which predicts the degree of interaction between proteins as determined using the String (v11) software.

**Figure 13:**
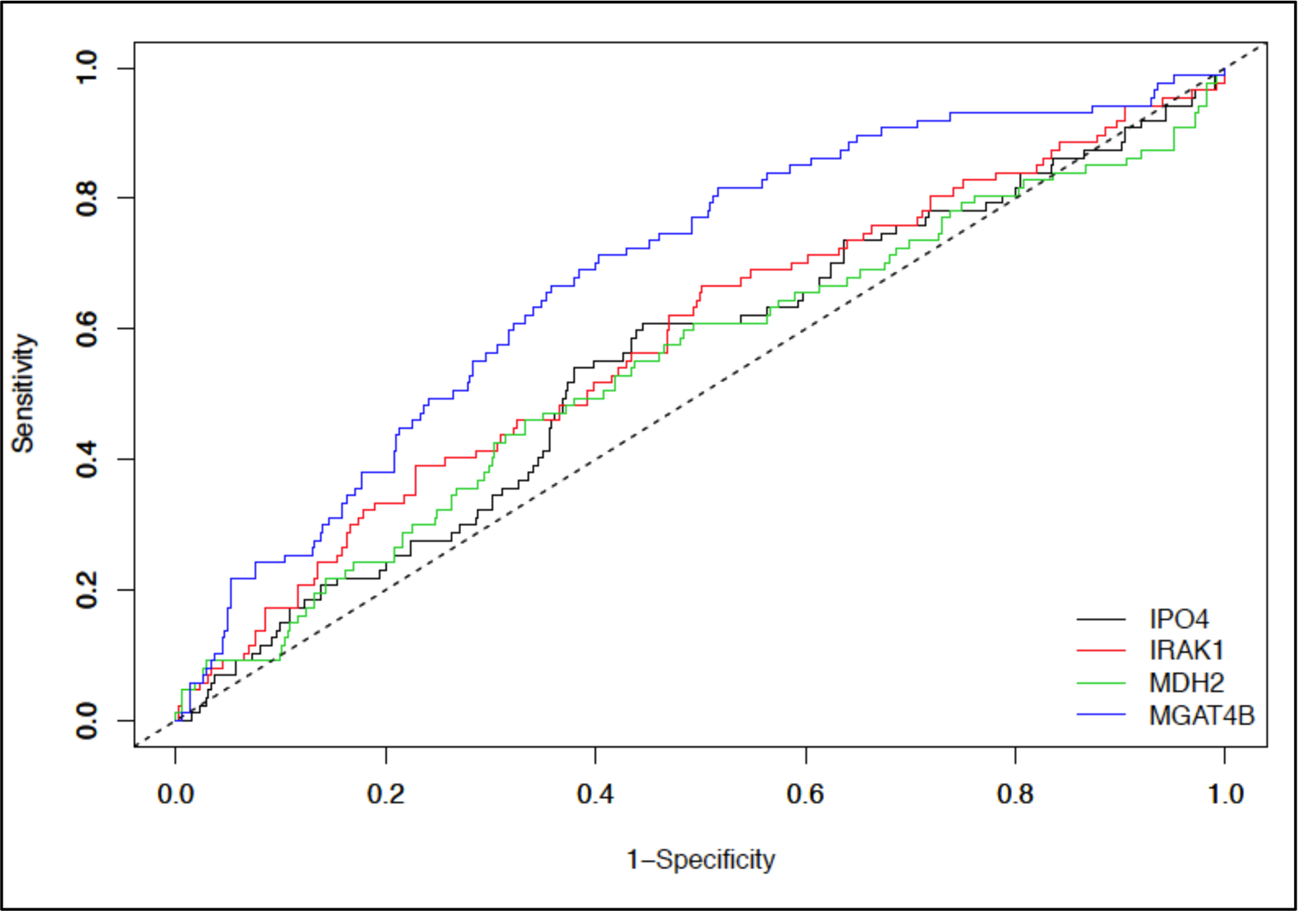
Receiver Operating Characteristic (ROC) curve analysis for IRAK1 and its closest neighbors in the magenta module. The ROC analysis reveals the capability of IRAK1 and its interacting neighbors in the magenta module to predict progression survival risk. The x-axis represents the sensitivity, and the y-axis represents the 1-specificity. The black dotted lines represent the line of significance.

### Identifying IRAK1 as a clinical biomarker for prostate cancer

Once we have established the clinical significance of increased expression of IRAK1 in all subtypes of PCa patients as well as in progression was established, we decided to determine its significance as a biomarker for PCa development and compared its clinical significance with other members of the IRAK family using AUROC analysis (**Figure 11**). IRAK1 was able to predict the presence of PCa in 83% of 472 PRAD samples (AUC: 0.83, 95% CI: 0.79412 - 0.88118, Z-score:15.204; p < 0.05) with a sensitivity of 0.708 (95% CI: 0.666 - 0.748) and specificity of 0.865 (95% CI: 0.742 - 0.944) (**Figure 11A**). Though IRAK1 overexpression was able to predict and differentiate PCa patients from normal patients with high sensitivity and specificity, IRAK1 was only able to predict PCa progression in 58% (AUC: 0.582; 95% CI: 0.51 - 0.65; Z-score: 2.3380, p-value = 0.01939) of PRAD patients (**Figure 11B**). Also, the IRAK1 differential expression profile was able to predict PCa recurrence/relapse in 60.5% of PRAD patients (**Supplementary Figure S5**). Further analysis will be needed with a larger population/sample size. This is because a larger sample size gives room for more reliable results with greater precision and power. It will also make sense to determine the predictability of IRAK1 using AUC in other PCa patient subtypes – CRPC and NEPC.

The functional impact score for each IRAK1 mutation was predicted using COSMIC (recurrent or non-recurrent), Mutation Assessor (high, medium, low, neutral), SIFT (deleterious, deleterious low_confidence, tolerated low_confidence, tolerated), and Polyphen-2 (probably damaging, possibly damaging, benign) algorithms (Flanagan et al., 2010; Gnad et al., 2013). These scores were integrated and ranked to form a new scoring scale (severe, moderate, mild) (**Table 7**). The missense mutations (R666Q, P614S, L92Q, and M34V) of IRAK1 were predicted to have mild to moderate functional impacts on PRAD progression while the frameshift mutations (L56Afs*102 (Insertion) and F290Pfs*58 (Deletion)) of IRAK1 were predicted to have severe functional impacts on PRAD progression. Of all the IRAK family genes identified, IRAK1 was found to have the highest CNVs (especially amplifications) in CRPC and NEPC compared to mPRAD and iPRAD patients. Interestingly, no mutation was identified in CRPC and NEPC patients. The strong correlation or co-expression of IRAK CNVs, mutations, and mRNA expression levels with AR, PI3K, and AURKA signaling pathway genes suggest a possible role in PCa progression (**Figure 14C**).

**Figure 14:**
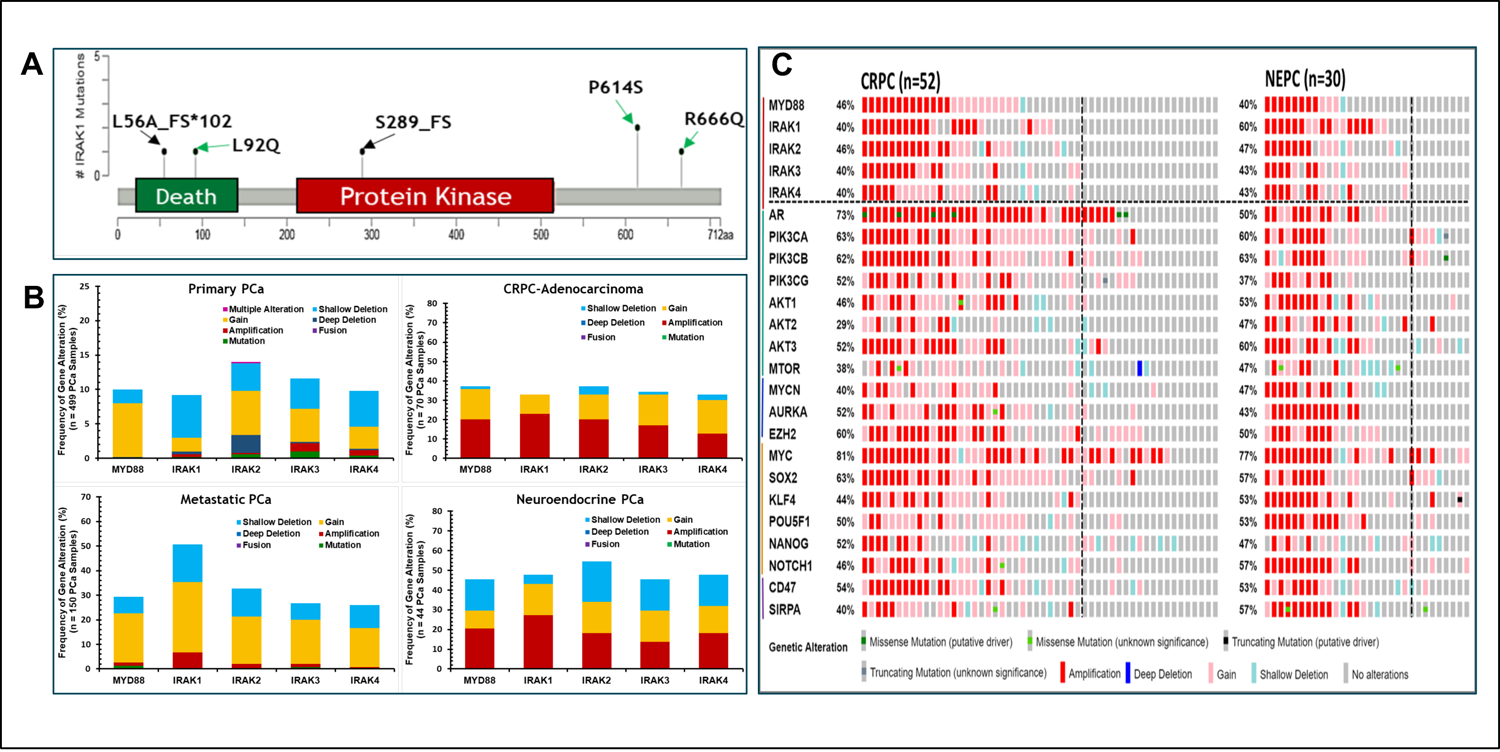
Gene alteration analysis output for IRAKs and associated genes. **A.** A Lollipop plot showing common structural mutation (missense and spliced) of IRAK1 in PCa patients from the 20 cohort studies analyzed. **B.** Frequency bar charts of alterations of MYD88 and IRAK1-4 genes in different PCa subtypes. Gene amplifications are common in CRPCs and NEPCs and relatively higher in IRAK1. **C.** Patient-based gene alteration profiles for IRAKs (MYD88 & IRAK1-4), castration resistance (AR-PI3K-AKT-mTOR), self-renewal/cancer pluripotency (MYC-SOX2-KLF4-POU5F1-NANOG-NOTCH1), neuroendocrine differentiation (MYCN-AURKA-EZH2), and “Don’t eat me” or immune evasion (CD47-SIRPA) signaling genes in CRPC (n = 52) and NEPC (n = 30) patients. Each box represents each PCa patient and the color-coding represents alteration type.

**Table 7:**
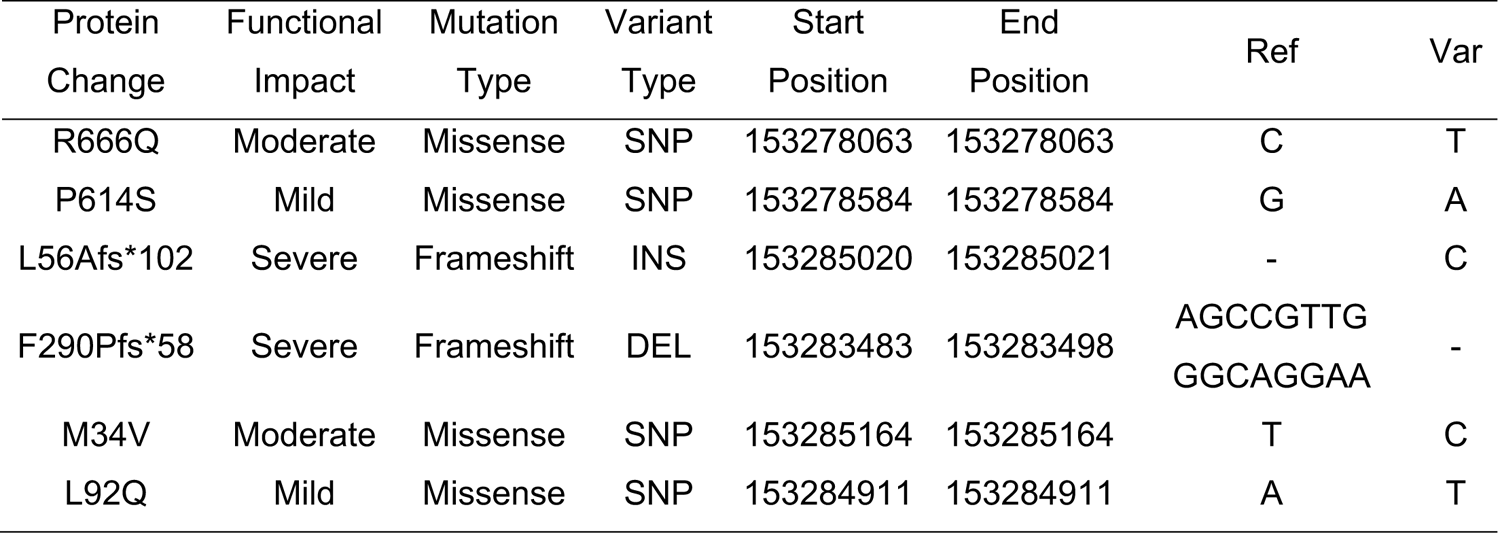
Functional impacts analysis and mutational landscape of IRAK1 in PRAD patients from 20 cohort studies. We identified 4 missense mutations (SNPs) and 2 frameshift mutations (insertion and deletion). The integrated functional impact analysis shows the frameshift mutations to be more pathogenic (severe) compared to the missense mutations (mild to moderate). The bioinformatics tools/algorithms and the ranking rubrics can be found in Supplementary Table S2.

Also, the TARGET module of the muTarget software was able to identify mutations in PRAD samples that are significantly (p < 0.05) associated with changes to IRAK1 expression. IRAK1 was found to be significantly (p < 0.05) and differentially expressed in PRAD patients with mutations in APC (Adenomatous Polyposis Coli Regulator of WNT Signaling Pathway), CDH1 (Chromodomain Helicase DNA Protein 1), TRIO (Trio Rho Guanine Nucleotide Exchange Factor), MUC17 (Mucin 17) and ADAMTSL1 (A Disintegrin and Metalloproteinase with Thrombospondin Motif) (**Figure 15**). IRAK1 was highly expressed in PRAD samples with mutations in APC, CHD1, and MUC17 genes compared to those without (Wildtype) whereas lower expression of IRAK1 was observed in PRAD samples/patients with ADAMTSL1 and TRIO mutations compared to the wildtype.

**Figure 15:**
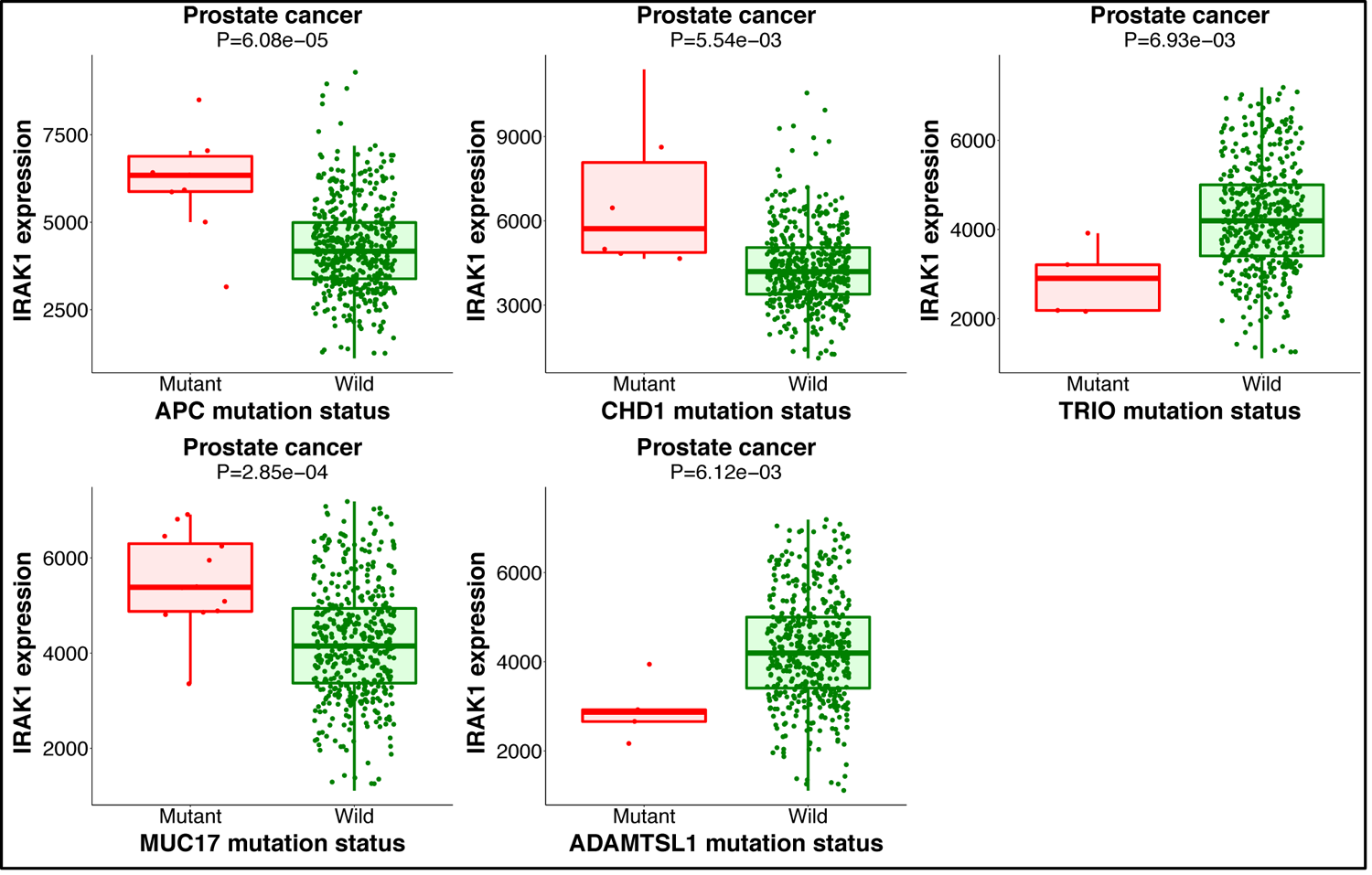
Box plots showing the association between mutation of APC, CHD1, TRIO, MUC17, and ADAMTSL1 and IRAK1 expression in PRAD samples. IRAK1 was significantly (p < 0.05) and highly expressed in PRAD samples with mutations in APC, CHD1, and MUC17 genes compared to those with no mutations (Wild-type). Whereas lower expression of IRAK1 was observed in PRAD samples/patients with ADAMTSL1 and TRIO mutations compared to the wild-type. Results were generated using the “TARGET” analysis module of MuTarget software, available at http://www.mutarget.com/.

### Hypomethylation of IRAK1 correlates with its overexpression and deregulation in PRAD samples/patients

To determine whether DNA methylation and epigenetic modifications played any role in the dysregulation and overexpression of IRAK1 in PRAD patients, we analyzed methylation data sets as highlighted in the methods section. Around 13 CpG sites were found in the IRAK1 gene transcript, with most of the methylation lesions located either close to or exactly on the promoter sites of the IRAK1 transcript known as “Island” (**Figure 16**). Concerning the CpG island (CGI), the CpG island shores were located within 2 kb regions either upstream (N-Shore) or downstream (S_Shore) of the CpG islands. The Shelves are located within a 2 kb region either upstream (N_Shelf) or downstream (S_Shelf) of the CpG island shores. Open Seas are isolated CpGs in the IRAK transcript. To identify the CpG islands (CGI) for IRAK1, we located regions with nucleotide regions with approximately 200bp in length and greater than 50% GC (Guanine-Cytosine) as well as observed/expected CpG ratio greater than 60%.

**Figure 16:**
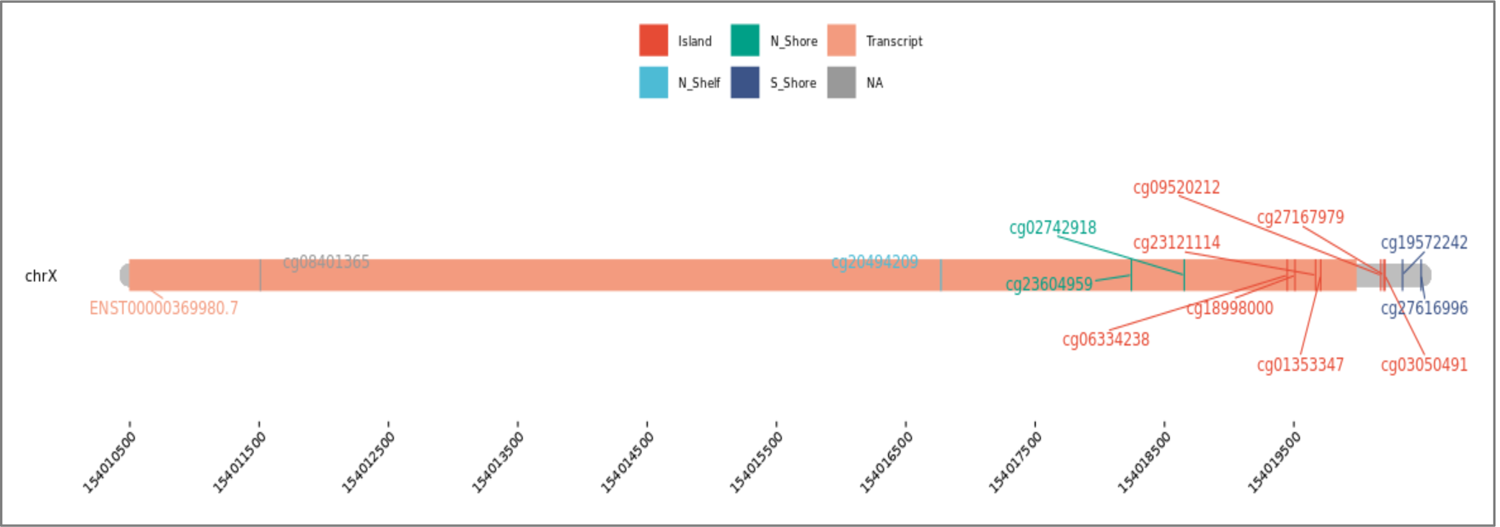
A schematic illustration of IRAK1 transcript showing the different CpG sites, including the Islands, N_Shores, S_Shores, N_Shelf, S_Shelf, and Open Seas.

About 7 of the CPG sites or probes were located on the CGI island, while 2 were located north of the island (N_Shore) and 2 on the southern side of the Island (S_Shore). The other two were located at the N_Shelf and the Open Sea (**Table 6**). The 13 CpG probes including cg08401365, cg20494209, cg23604959*, cg02742918*, cg06334238, cg18998000, cg23121114, cg01353347*, cg09520212, cg27167979*, cg03050491, cg19572242*, and cg27616996* were identified on both the promoter and transcription sites (CGI positions: Islands, N_Shores, S_Shores, N_Shelves, S_Shelves, and Open Seas). About 6 CpG probes (*) out of the 13 associated with the IRAK1 gene were found to be statistically significant (Wilcoxon test; p < 0.05) and hypomethylated based on the significance of their methylation value (or a beta-value <0.05) in PRAD patients (n = 492) compared to normal patients (n = 50) (**Figure 17**). The mean methylation values for all CpGs (Aggregation) for IRAK1 were calculated and found to have a negative correlation with IRAK1 gene expression (r = −0.41, p = 2.2e-16) as seen in **Figure 18**. This indicates that hypomethylation of IRAK1 may have some impact on the overexpression of the gene in PRAD patients.

**Figure 17:**
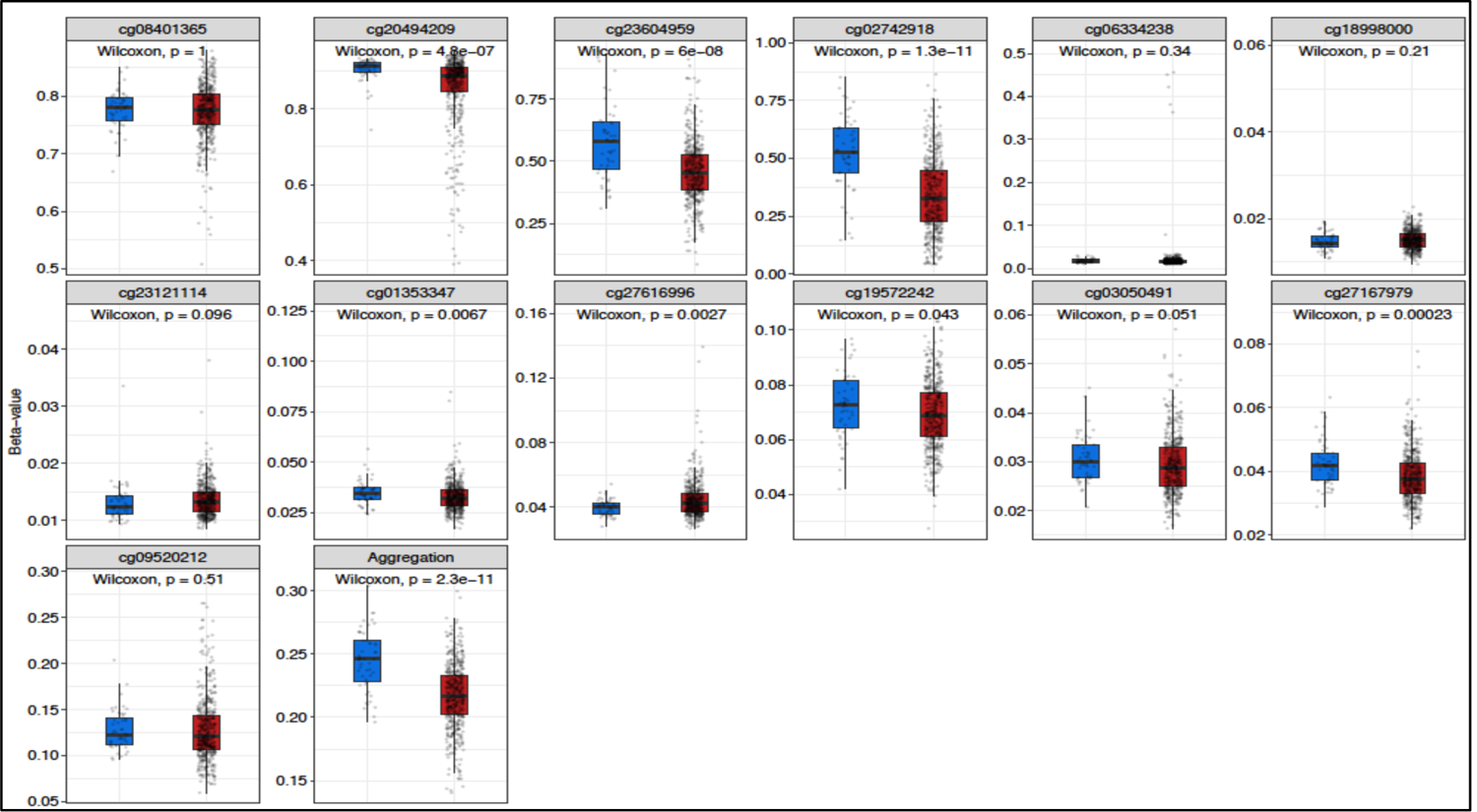
Box plots of the differential methylation between PRAD and normal samples. The red boxes represent the distribution of CpG lesions in tumor samples while the blue boxes represent the distribution of CpG lesions in normal samples. Based on the p-value of the Wilcoxon test, cg23604959, cg02742918, cg01353347, cg27167979, cg19572242, and cg27616996 were found to be hypomethylated compared to the normal samples. Further mechanistic studies will be needed to better understand the functional impacts of IRAK1 hypomethylation (epigenetics) on PCa initiation and progression.

**Figure 18:**
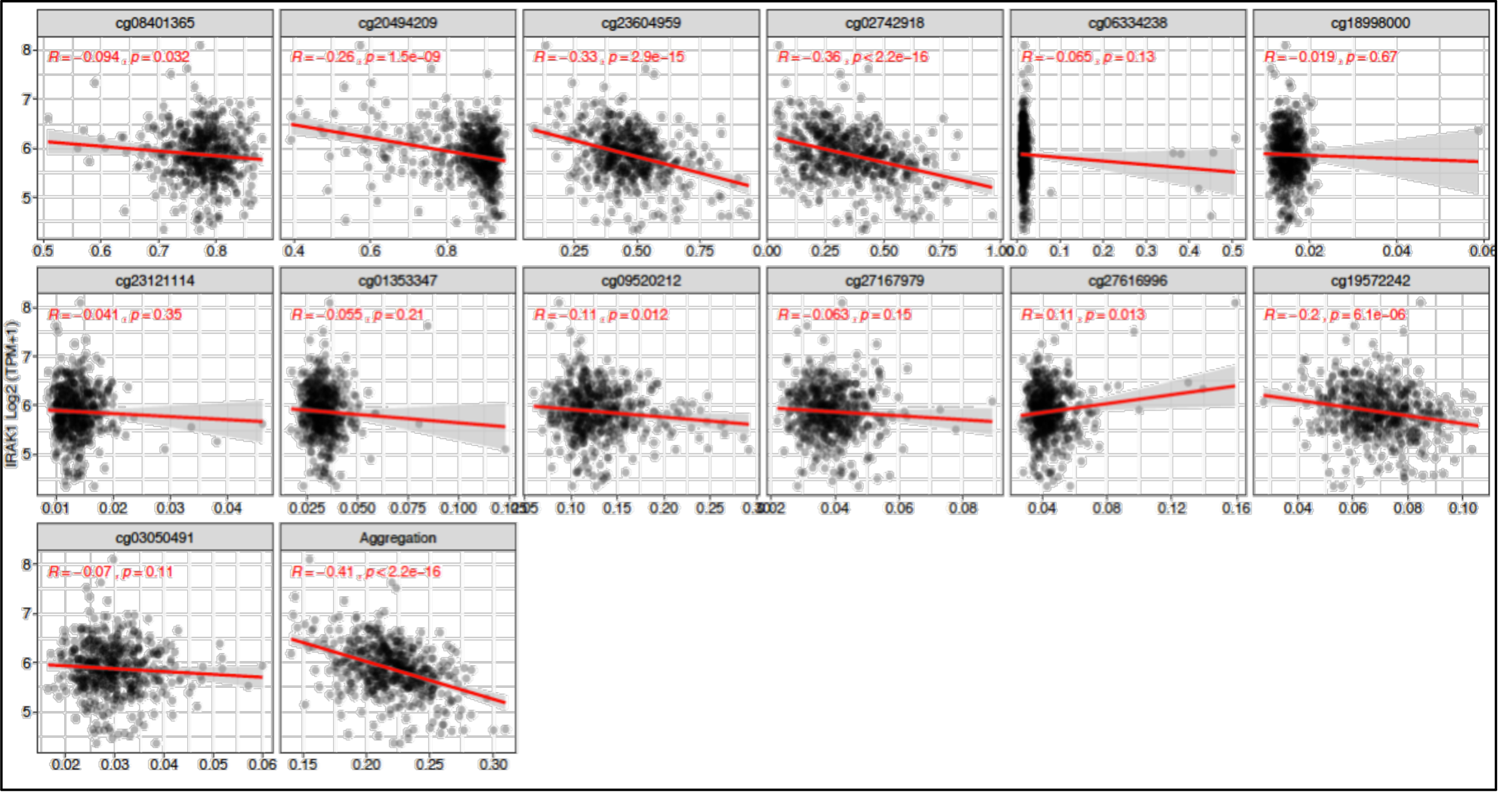
Correlation analysis plots using Pearson coefficient to identify the correlation between IRAK1 methylation and its transcript or gene expression. The aggregate mean methylation values for all13 methylation lesions shows a negative correlation to IRAK1 expression (p = 2.2e-16, r = −0.41). A total of 7 CpG probes show a negative correlation at a significant p-value of less than 0.05 while only one probe (cg27616996) shows a positive correlation at a p < 0.05.

We also determined the correlation between DNA methylation (CpGs m-values) of IRAK1 and the observed copy number variation (CNVs; − 2: homozygous deletion; − 1: single copy deletion; 0: diploid normal copy; + 1: low-level copy number amplification; + 2: high-level copy number amplification) in PRAD samples. Our analysis showed that there is a significant difference between methylation of IRAK1 and the various CNVs identified in IRAK1 (**Figure 19**). 6 of the 13 CpGs show a significant difference between IRAK1 methylation and each CNV. cg18996000 has a higher level of hypomethylation with single-copy deletion compared to others (m value, p = 0.0056). cg20494209 has a lower level of hypermethylation with homozygous deletion compared to others (m value, p = 0.0034), while cg02742918 has a lower level of hypermethylation with high-level copy number amplification compared to others (m value, p = 0.045). cg27616996 has a higher level of hypomethylation with single-copy deletion compared to others (m value, p = 0.0056), while cg09520212 has a lower level of hypomethylation in all CNVs compared to the diploid normal copy (m value, p = 0.047), while cg2312114 has a lower level of hypomethylation in low-level and high-level copy number amplification compared to others (m value, p = 0.015). The mean aggregate methylation value for all CpGs shows a higher level of hypomethylation for homozygous deletion, single copy deletion, low-level copy number amplification, high-level copy number amplification compared to the diploid normal copy (m value, p = 6e-04).

**Figure 19:**
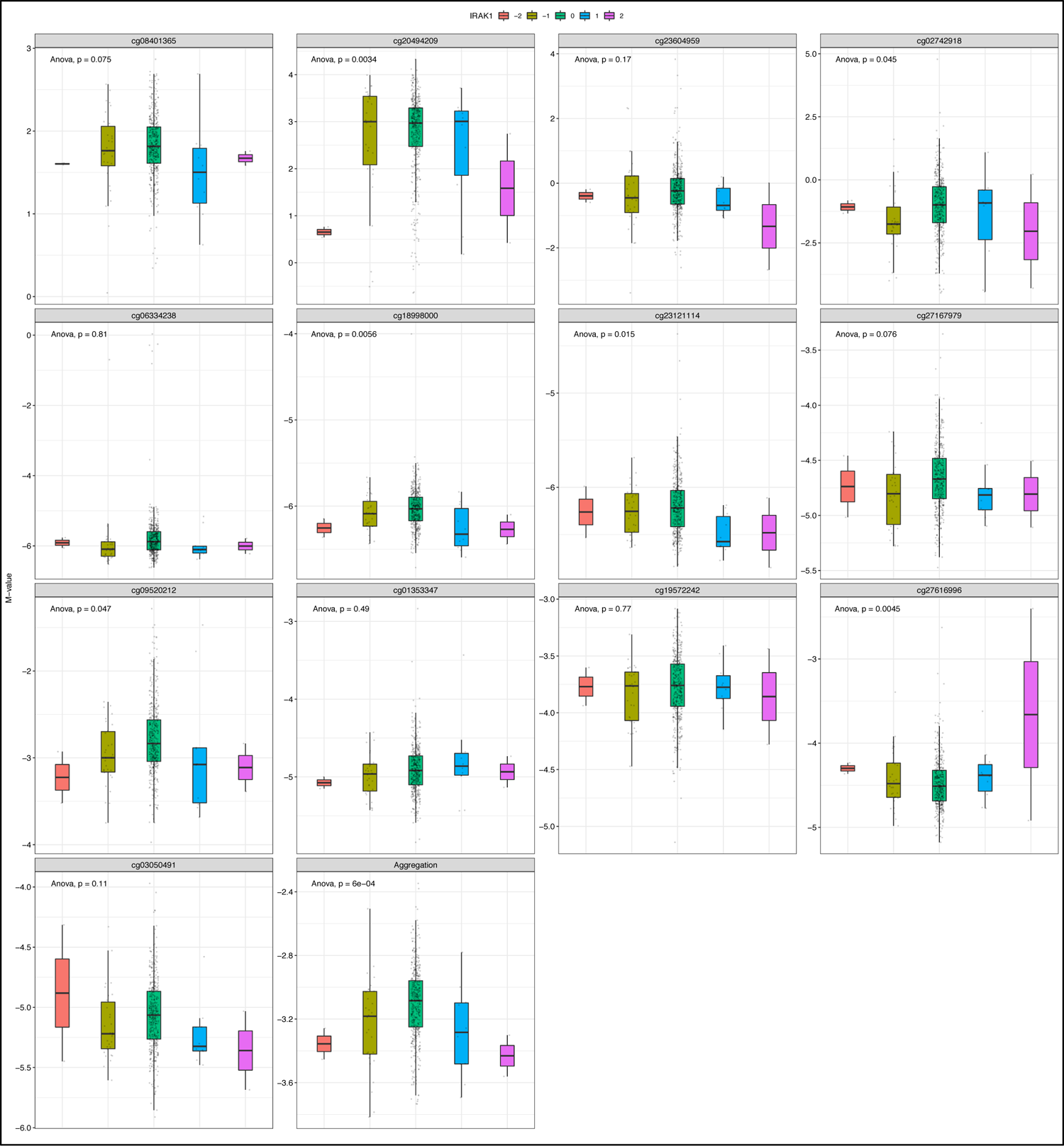
Box plots showing the relationship between methylation values (for each CpG probe) and copy number variations (CNVs) of IRAK1. Of the 13 CpGs identified, 6 show a significant difference between IRAK1 methylation and each CNV (−2: homozygous deletion; − 1: single copy deletion; 0: diploid normal copy; + 1: low-level copy number amplification; + 2: high-level copy number amplification). cg18996000 shows a higher level of hypomethylation with single-copy deletion compared to others (m value, p = 0.0056). cg20494209 shows a lower level of hypermethylation with homozygous deletion compared to others (m value, p = 0.0034). cg02742918 shows a lower level of hypermethylation with high-level copy number amplification compared to others (m value, p = 0.045). cg27616996 shows a higher level of hypomethylation with single-copy deletion compared to others (m value, p = 0.0056). cg09520212 shows a lower level of hypomethylation in all CNVs compared to the diploid normal copy (m value, p = 0.047). cg2312114 shows a lower level of hypomethylation in low-level and high-level copy number amplification compared to others (m value, p = 0.015). The mean aggregate methylation value for all CpGs shows a higher level of hypomethylation for homozygous deletion, single copy deletion, low-level copy number amplification, high-level copy number amplification compared to the diploid normal copy (m value, p = 6e-04). Each box is colored to represent: −2 (coral), −1 (olive green), 0 (green), +1 (blue), and +2 (magenta).

Additionally, we analyzed the correlation between IRAK1 methylation values and each IRAK1 transcript/isoform. Of all the 13 transcripts or isoforms identified for IRAK1, only 6 showed significant negative correlation (p < 0.05) with IRAK1 methylation. The mean aggregate methylation values of the 6 IRAK1 isoforms: ENST00000369974.6 (r = −0.338; p = 8.4e-15), ENST00000369980.7 (r = −0.19; p = 8e-06), ENST00000429936.6 (r = −0.15; p = 0.00079), ENST00000455690.5 (r = −0.14; p = 0.00085), ENST00000463031.1 (r = −0.13; p = 0.0023), ENST00000467236.1 (r = −0.21; p = 1.36e-6) were found to be significantly and negatively correlated with expression of each IRAK1 isoform (**Supplementary Figures S18A - M**).

### IRAK1 is overexpressed in PRAD patients and PCa cell lines at mRNA and protein levels

To better understand how IRAK1 interacts with its upstream and downstream regulators in PRAD patients, we performed both IPA and correlogram analysis and visualized the data outputs using heatmaps. Our analysis revealed that IRAK1 is overexpressed in most PRAD, CRPC, and NEPC patients (**Figures 19 - 21, right panel**). There was a positive correlation between IRAK1 and NF-κB-associated genes in indolent PRAD samples (**Figure 19, left panel**). A similar pattern was observed in CRPC and NEPC patients (**Figures 20 - 21, left panel**). However, more upstream and downstream inflammatory genes were found to be positively correlated with IRAK1 expression in NEPC and CRPC compared to samples with indolent PRAD. Furthermore, we confirmed that IRAK1 was highly expressed in 19 of the 30 diverse tumor types when compared to normal samples based on analysis of the TCGA PanCancer dataset (**Figure 22**). IRAK1 is the most upregulated of the 4 IRAK family genes while IRAK3 is most downregulated in PRAD, NEPC, and CRPC samples (**Supplementary Figures S6 - S8**). Also, high IRAK1 expression was found in PCa cell lines. The expression profile of IRAK1 appears to be higher in androgen receptor-negative (AR-) PCa cell lines (PC3 and DU143) compared to the androgen receptor-positive (AR+) PCa cell lines (LNCaP and C42). PC3 cells displayed the highest mRNA expression followed chronologically by DU143, C42, and LNCaP (**Figure 23**). IRAK1 protein profile of PRAD microarray tissues present in the protein atlas database shows that IRAK1 is highly expressed in high-grade compared to the low-grade PCa tissue samples (**Figures 24 and 25**). All normal prostate tissues (n = 3) included in the microarray showed low to no IRAK1 expression (Uhlén et al., 2005).

**Figure 20:**
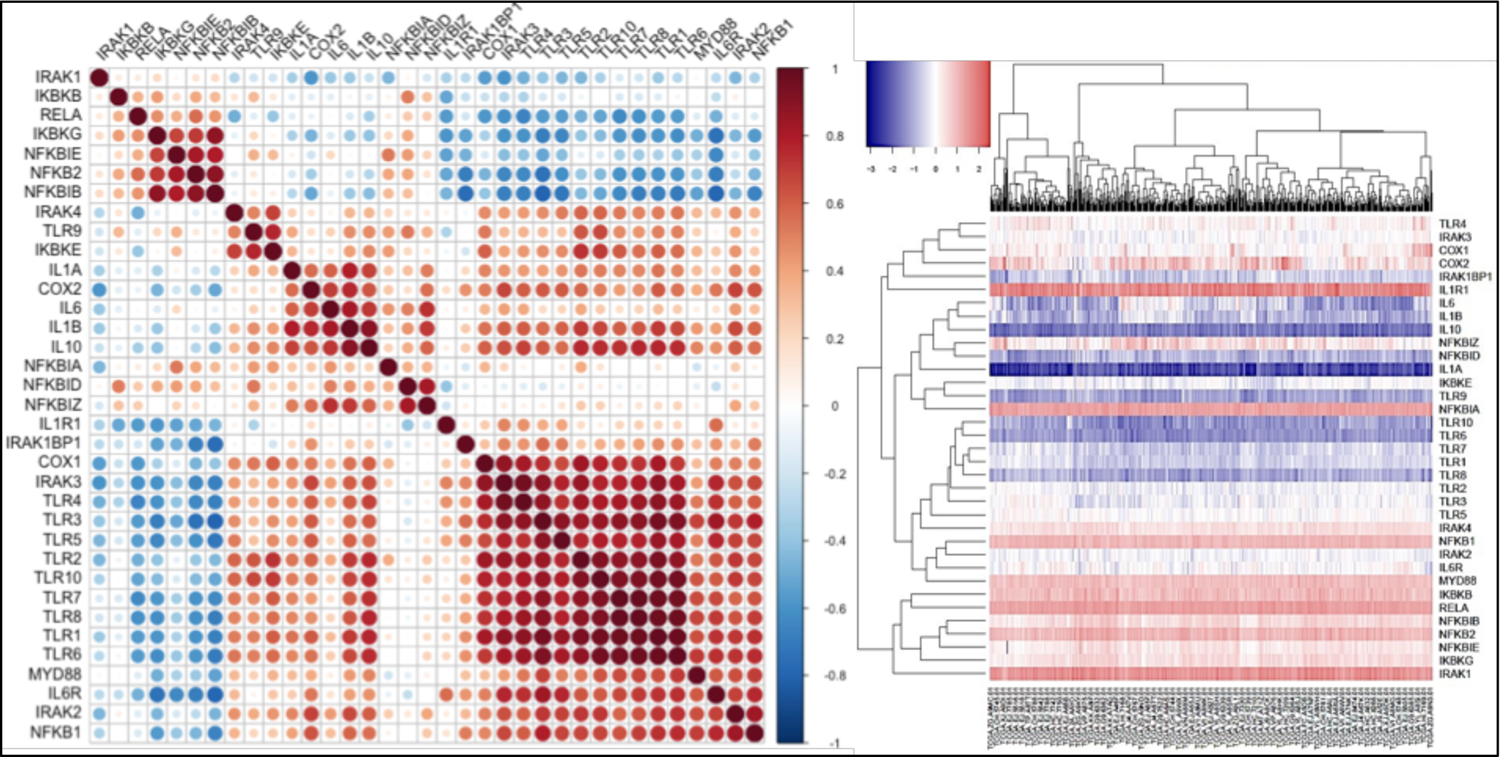
Heatmaps and correlogram showing gene expression profiles of inflammatory genes in PRAD samples or patients and the correlation between different inflammatory genes. IRAK1 was found to be highly expressed in almost all PRAD samples (n = 472). Other IRAK (2, 3, and 4) genes show low or moderate expressions in all PRAD samples. IRAK1 expression correlated positively with NF-κB transcriptional factor activators and co-expressors (Left Panel). Thus, indicating the potential role of targeting IRAK1 as a means to inhibit NF-κB signaling and pro-tumor function in PRAD patients.

**Figure 21:**
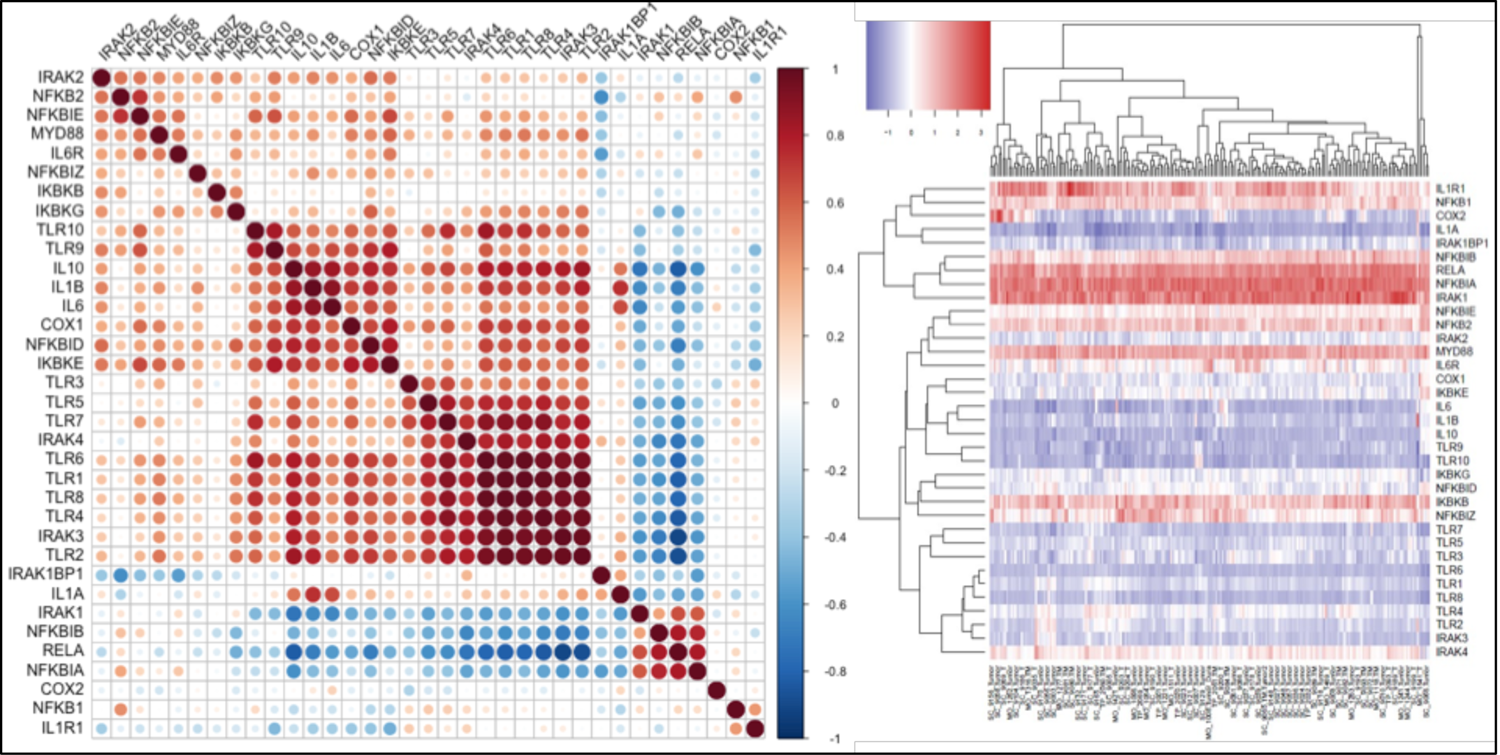
Heatmap and correlogram showing gene expression profiles of inflammation-associated genes in castration-resistant PCa (CRPC) samples or patients and the correlation between different inflammation-associated genes. IRAK1 was found to be highly expressed in almost all CRPC samples (n = 444). Other IRAK (2, 3, and 4) genes show low or moderate expressions in all CRPC samples. IRAK1 correlated positively with NF-κB transcriptional factor activators and co-expressors (**Left Panel**). Thus, indicating the potential role of targeting IRAK1 as a means to inhibit NF-κB signaling and pro-tumor function in CRPC patients.

**Figure 22:**
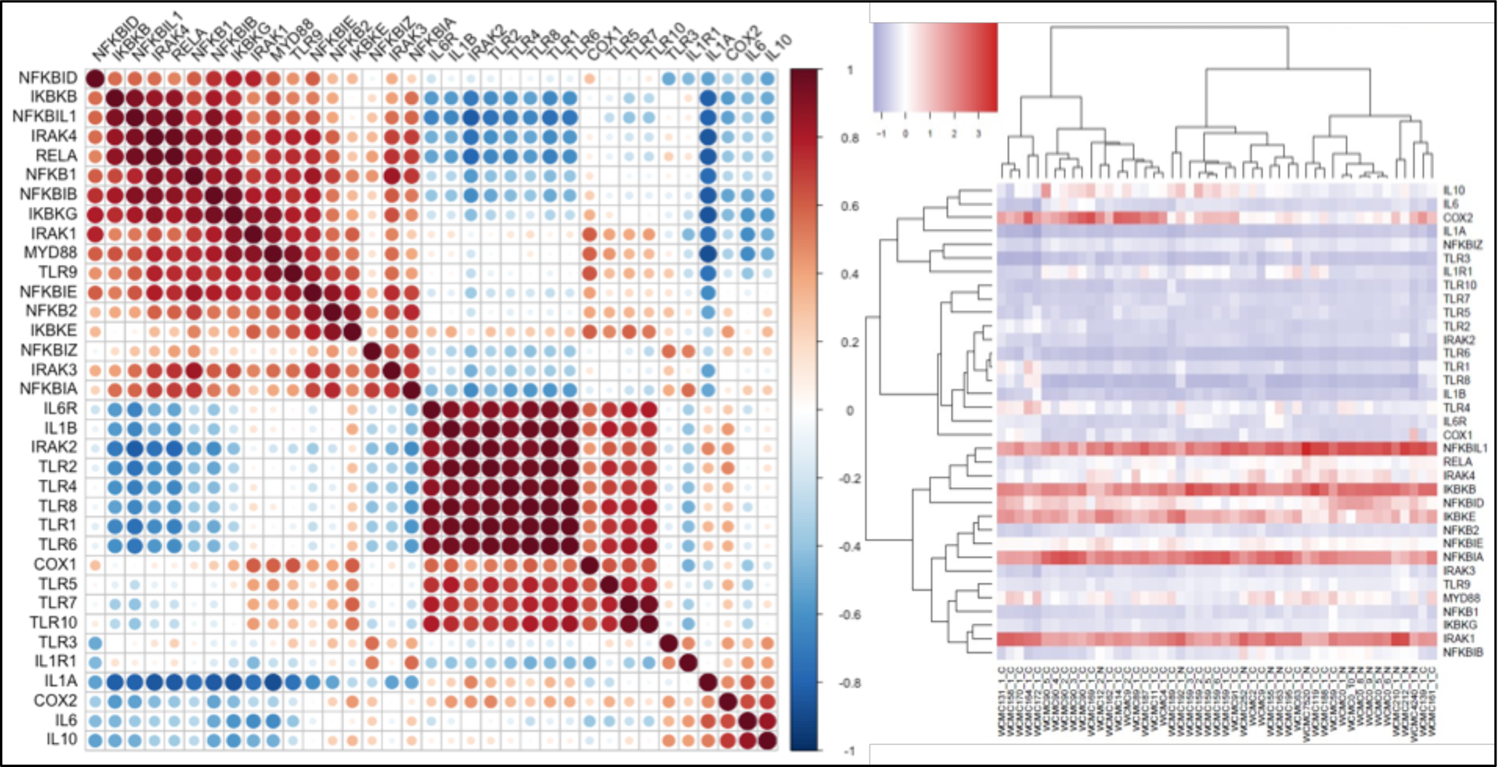
Heatmap and correlogram showing gene expression profiles of inflammation-associated genes in Neuroendocrine PCa (NEPC) samples or patients and the correlation between different inflammatory genes. IRAK1 was found to be highly expressed in almost all NEPC samples (n = 30). Other IRAK (2, 3, and 4) genes show low or moderate expressions in all NEPC samples. IRAK1 correlated massively and positively with NF-κB transcriptional factor activators and co-expressors as well as with many TLRs (**Left Panel**). Thus, indicating the potential role of targeting IRAK1 as a means to inhibit NF-κB signaling and pro-tumor function in NEPC patients.

**Figure 23:**
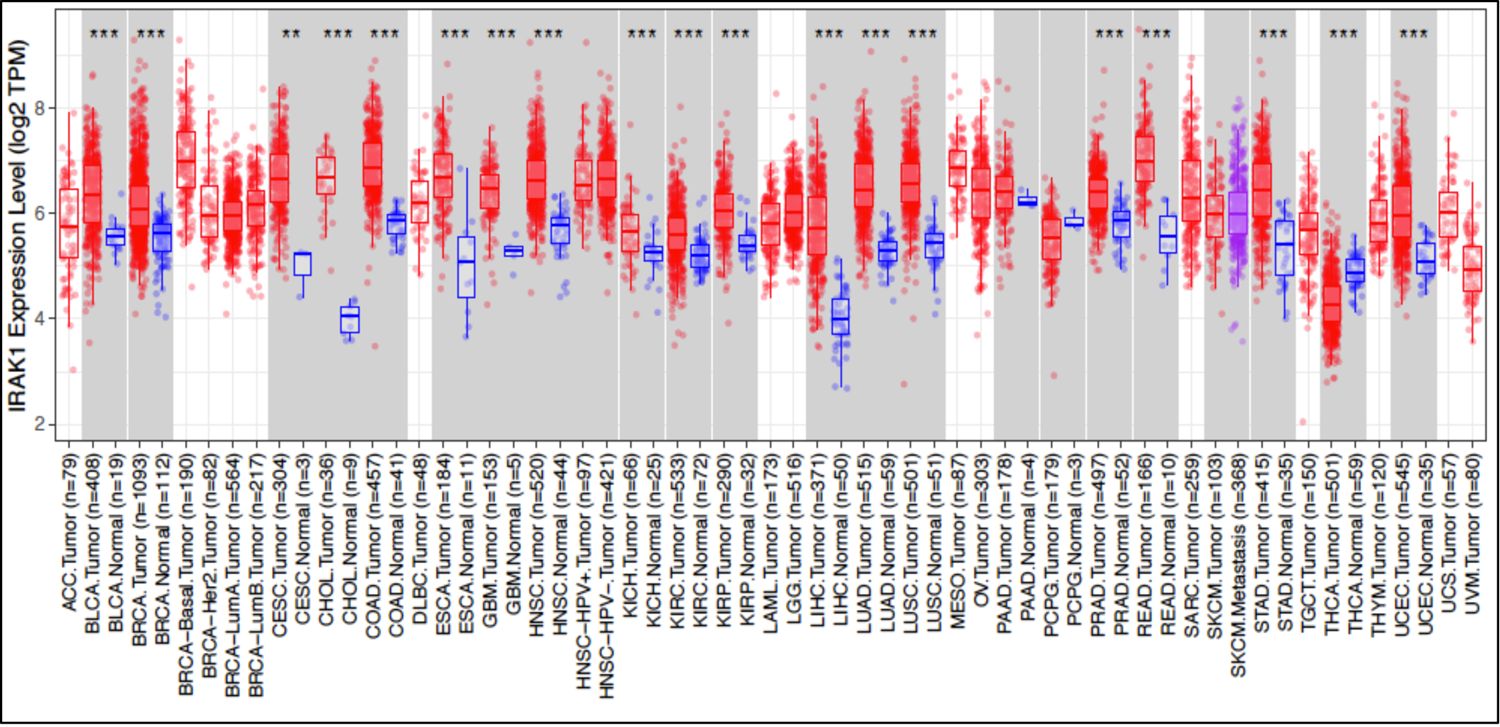
Box plot showing differential expression of IRAK1 in multiple tumor types in the TCGA Pancancer data set. Most tumors show very high expression of IRAK1 compared to normal samples. Each acronym represents the names of cancer types as described below. LAML: Acute Myeloid Leukemia, ACC: Adrenocortical carcinoma, BLCA: Bladder Urothelial Carcinoma, LGG: Brain Lower Grade Glioma, BRCA: Breast invasive carcinoma, CESC: Cervical squamous cell carcinoma and endocervical adenocarcinoma, CHOL: Cholangiocarcinoma, LCML: Chronic Myelogenous Leukemia, COAD: Colon adenocarcinoma, CNTL: Controls, ESCA: Esophageal carcinoma, GBM: Glioblastoma multiforme, HNSC: Head and Neck squamous cell carcinoma, KICH: Kidney Chromophobe, KIRC: Kidney renal clear cell carcinoma, KIRP: Kidney renal papillary cell carcinoma, LIHC: Liver hepatocellular carcinoma, LUAD: Lung adenocarcinoma, LUSC: Lung squamous cell carcinoma, DLBC: Lymphoid Neoplasm Diffuse Large B-cell Lymphoma, MESO: Mesothelioma, OV: Ovarian serous cystadenocarcinoma, PAAD: Pancreatic adenocarcinoma, PCPG: Pheochromocytoma and Paraganglioma, PRAD: Prostate adenocarcinoma, READ: Rectum adenocarcinoma, SARC: Sarcoma, SKCM: Skin Cutaneous Melanoma, STAD: Stomach adenocarcinoma, TGCT: Testicular Germ Cell Tumors, THYM: Thymoma, THCA: Thyroid carcinoma, UCS: Uterine Carcinosarcoma, UCEC: Uterine Corpus Endometrial Carcinoma, UVM: Uveal Melanoma; P-value (* < 0.05; ** < 0.01; *** < 0.001).

**Figure 24:**
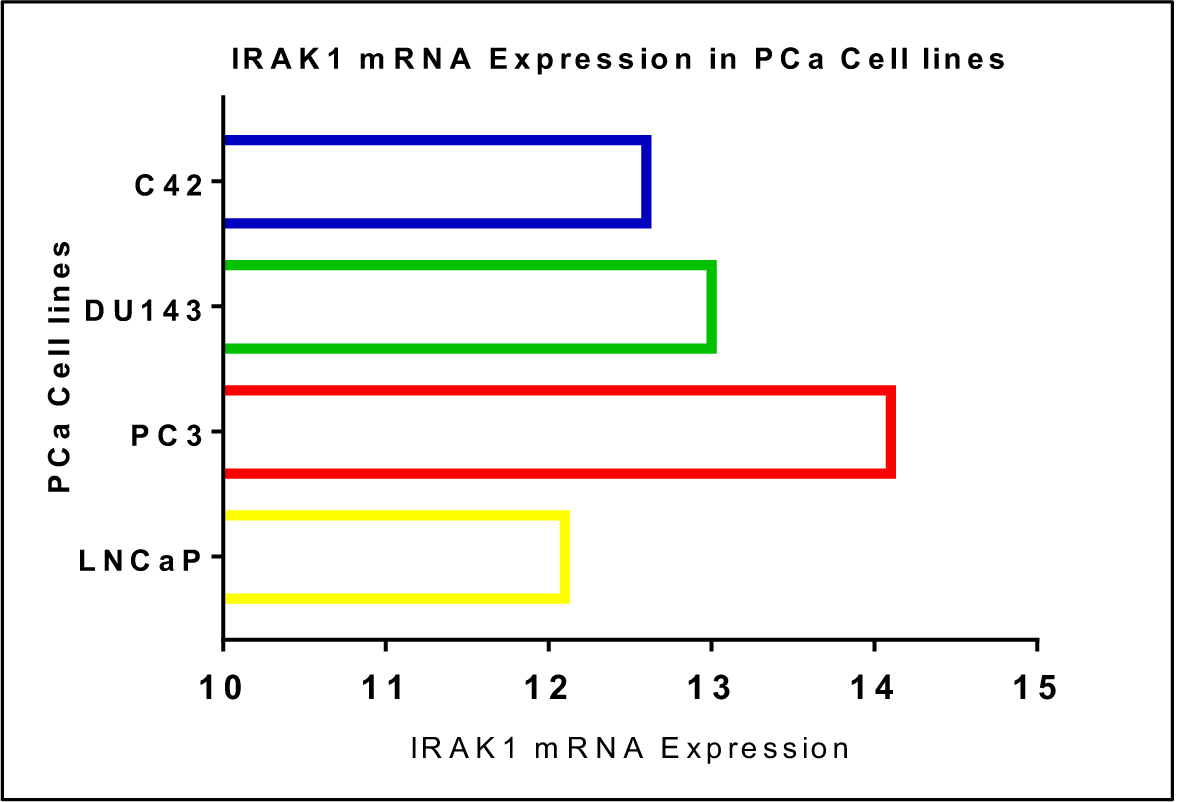
Bar chart showing the mRNA expression profile of IRAK1 in 4 prostate cancer cell lines. The PC3 cell line has the highest IRAK1 expression level while the LNCaP cell line has the lowest IRAK1 levels. https://maayanlab.cloud/archs4/gene/IRAK1#tissueexpression

**Figure 25:**
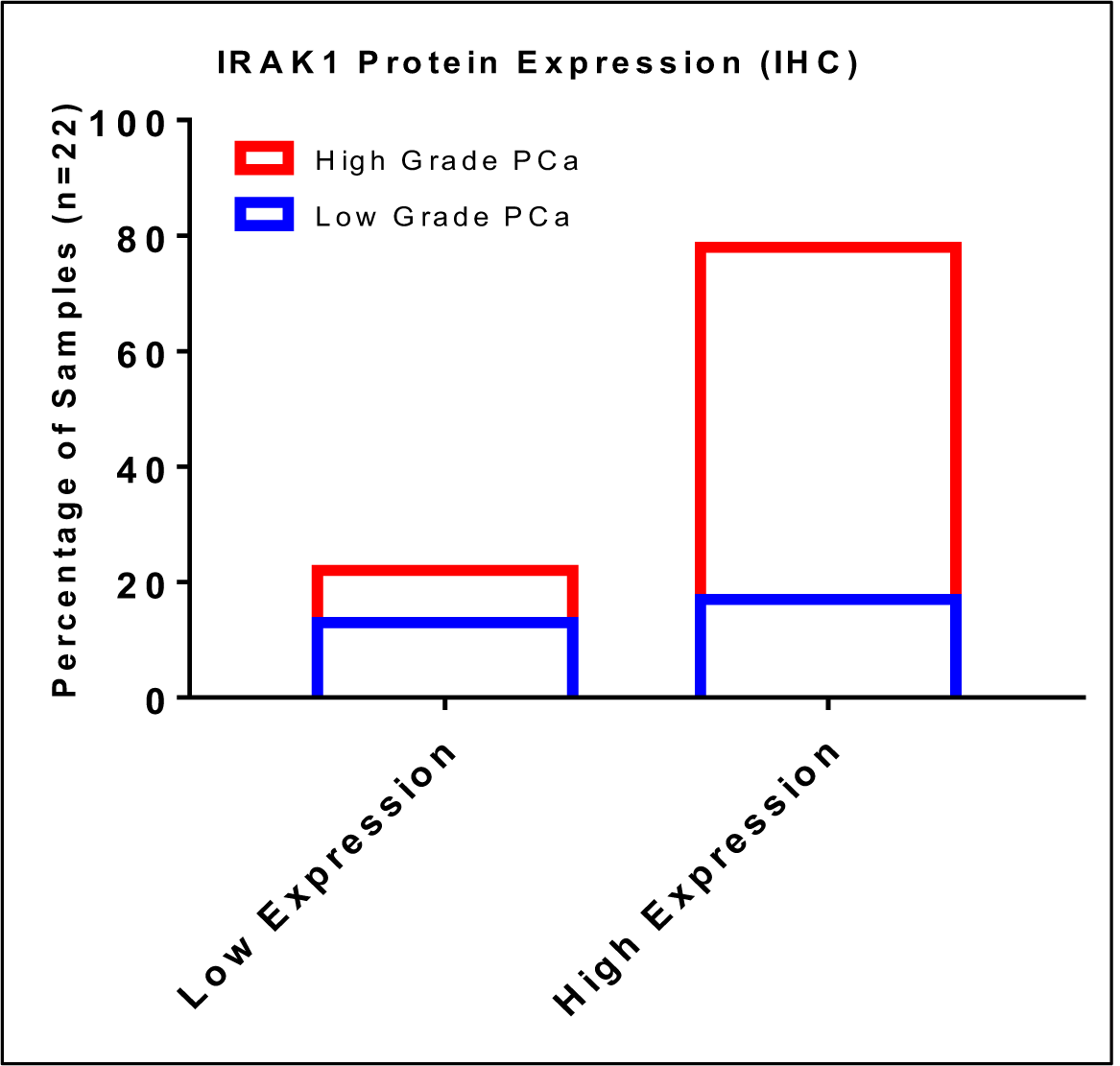
Bar chart showing the IRAK1 protein expression profile in PRAD tissue microarray (n=22) acquired from the human protein atlas database. About 61% of high-grade (Stage 8-10) and 17% of low-grade PCa samples show high expression of IRAK1.

## Discussion

In this study, we have characterized the transcriptome, genome, and epigenome features of PCa tumorigenesis with emphasis on inflammatory and PCa progression signaling mechanisms. We identified inflammatory genes undergoing deregulation, genetic alteration (mutation and CNA/CNV), and DNA methylation in our RNAseq dataset that could drive PCa progression from indolent PRAD to metastatic CRPC or NEPC. WGCNA was able to construct modules within the biological networks based on the pairwise correlation between expressed genes in each module and the clinical/phenotypic traits of interest to describe the module relatedness and identify putative hub genes of clinical significance. Importantly, WGCNA facilitated a network-based identification and characterization of inflammatory genes that are simultaneously associated with PCa progression as candidate diagnostic or prognostic biomarkers and therapeutic targets (Langfelder & Horvath, 2008). Overall, our analysis identified 10 inflammatory genes: IRAK1, PPIL5/LRR1, HMGB3, HMGB2, TRAIP, IL1F5/IL36RN, ILF2, TRIM59, NFKBIL2/TONSL, and TRAF7, found to be associated with PCa progression.

Interleukin-1 receptor-associated kinase 1 (IRAK1) is a critical regulator of the TLR/IL-1 pathway and one of the four mammalian members of the IRAKs. IRAKs are a family of regulatory proteins with an important role in the inflammatory signaling cascades of two receptor families, toll-like receptors (TLRs) and interleukin-1 receptors (Jain et al., 2014). To this point, only four mammalian members of the IRAK family have been identified: IRAK1, IRAK2, IRAK3/IRAKM, and IRAK4. Interestingly, only IRAK1 and IRAK4 exhibit kinase-like activity, which makes them druggable and targetable candidates compared to IRAK2 and IRAK3, which are pseudokinases (Flannery & Bowie, 2010; Jain et al., 2014; Rothschild et al., 2018).

Functionally, IRAKs signaling is important in the activation of downstream inflammatory molecules, some of which are known to promote tumor growth, metastasis, immune suppression, and chemoresistance in tumor cells (Singer et al., 2018). Importantly, IRAK1 is involved in the activation of the NF-κB transcription activities (Zhang et al., 2016). The inhibition of IRAK1 and IRAK4 has been reported to be therapeutically beneficial in the treatment of melanoma, leukemia, pancreatic, Kaposi sarcoma, hepatocellular, cervical, and breast cancer, among others (Hu et al., 2018; Li et al., 2015; Sun et al., 2006; Sun et al., 2014; Wee et al., 2015; Zhang et al., 2016). Despite the available reports and data on the roles of IRAKs in several chronic diseases, including tumors, the oncogenic significance of IRAKs at the molecular level in PCa remains unclear.

Peptidylprolyl Isomerase (Cyclophilin)-Like 5 (PPIL5)/Leucine-Rich Repeat Protein 1 (LRR1) is a negative regulator of TNFRSF9/4-1BB, a member of the tumor necrosis factor receptor (TNFR) superfamily. Overexpression of LRR1 is believed to suppress the activation of NF-κB induced by TNFRSF9 or TNF receptor-associated factor 2 (TRAF2) (Jang et al., 2001). Both High Mobility Group Boxes 2 and 3 (HMGB2 and HMGB3) are known to act as chemokines that promote the proliferation and migration of epithelial and endothelial cells. They sometimes act as damage-associated molecular patterns (DAMPs) by inducing proinflammatory responses. HMGB1, another isomer, had previously been reported to induce sterile inflammation in prostate tumors (Pandey et al., 2015).

TRAF Interacting Protein (TRAIP) is known to interact with TRAF1 and TRAF2. It is a negative regulator of innate immune signaling and inhibits the activation of NF-κB1 mediated by TNF in immune cells. TRAIP has also been shown to promote malignant behaviors in liver cancer, melanoma, and breast cancer (Guo, 2020). However, its importance in PCa progression is unknown and needs to be defined. Tumor necrosis factor 7 (TRAF7) is another inflammatory gene identified in the magenta module. It is highly involved in inflammation by activation of the NEMO-RelA-NF-κB signaling pathways, as well as induction of tumor progression (Zhu et al 2018). The clinical significance of TRAF7 in PCa has not been studied and needs to be outlined. Interleukin Enhancer Binding Factor 2 (ILF2) is a transcriptional factor known to be involved in the expression of IL-2. The knockdown of ILF2 has been shown to impede cell growth while its upregulation has been linked with poor prognosis and clinical outcome in gastric cancer and hepatocellular carcinoma (Cheng et al., 2016, Yin et al., 2017). Whether dysregulation of ILF2 is clinically significant in PCa progression has not been studied.

IL-1 Family member 5 (IL1F5)/Interleukin 36 Receptor Antagonist (IL36RN) encodes an anti-inflammatory cytokine known as IL36Ra. This gene has been associated with tumor progression in breast, colorectal, bladder, ovarian, and lung cancers (Walsh & Fallon, 2018). Whether a similar effect can be replicated in PCa is unknown. However, IL1F is mainly produced by epithelial cells and is shown to have multiple roles through its interaction with several inflammatory genes in cancers (Queen et al., 2019). Tripartite Motif Containing 59 (TRIM59) is a multifunctional regulator of the innate immune signaling pathway. The overexpression of TRIM59 has been correlated with poor prognosis in multiple tumors, including pancreatic cancer, breast cancer, and ovarian cancer (Mandell et al., 2020). Also, its upregulation has been shown to promote cancer cell proliferation (Liu et al., 2018). The clinical importance of TRIM59 in PCa is currently unknown and needs to be clarified. Though NF-κB Inhibitor-like Protein 2 (NFKBIL2)/Tonsoku-like DNA Repair Protein (TONSL) interacts with NF-κB to prevent its activation, the overexpression of TONSL has been identified as an independent unfavorable prognostic indicator of hepatocellular carcinoma (Yu et al., 2019).

Although many of the identified inflammatory genes have both inflammatory and non-inflammatory properties, what makes them significant in tumor progression depends not only on their inflammatory functions but also on their non-inflammatory oncogenic or tumor-suppressing abilities such as DNA binding and repair abilities, apoptosis regulation, regulation of cell differentiation, division, metabolism and proliferation (**Supplementary Table S1**). Detailed mechanistic studies are needed to specifically elucidate the individual role of these inflammatory genes in PCa progression will be beneficial in developing novel therapeutic strategies targeting the dysregulation of these genes in PRAD/PCa cells.

There is no doubt that the 10 inflammatory genes identified have the potential to play a significant impact in PRAD progression based on results from our WGCNA and post hoc or multivariate analyses (**Figures 2 - 26 and S1 - S18, Tables 2 - 7 and S1 - S2**). AUROC and KM survival analyses revealed the diagnostic potentials of some of these inflammatory genes, especially with regards to PRAD development and progression. Nevertheless, the expression of most genes in the magenta module was predicted to be positively correlated with PRAD progression and relapse. In this study, we specifically focused on IRAK1 due to its critical role in the inflammatory signaling cascades as well as its association with carcinogenesis from previous studies (Kutikhin & Yuzhalin, 2012; Li et al., 2015; Sun et al., 2014). Further downstream analyses were performed on IRAK1 as a representative of the 10 identified inflammatory genes.

**Figure 26:**
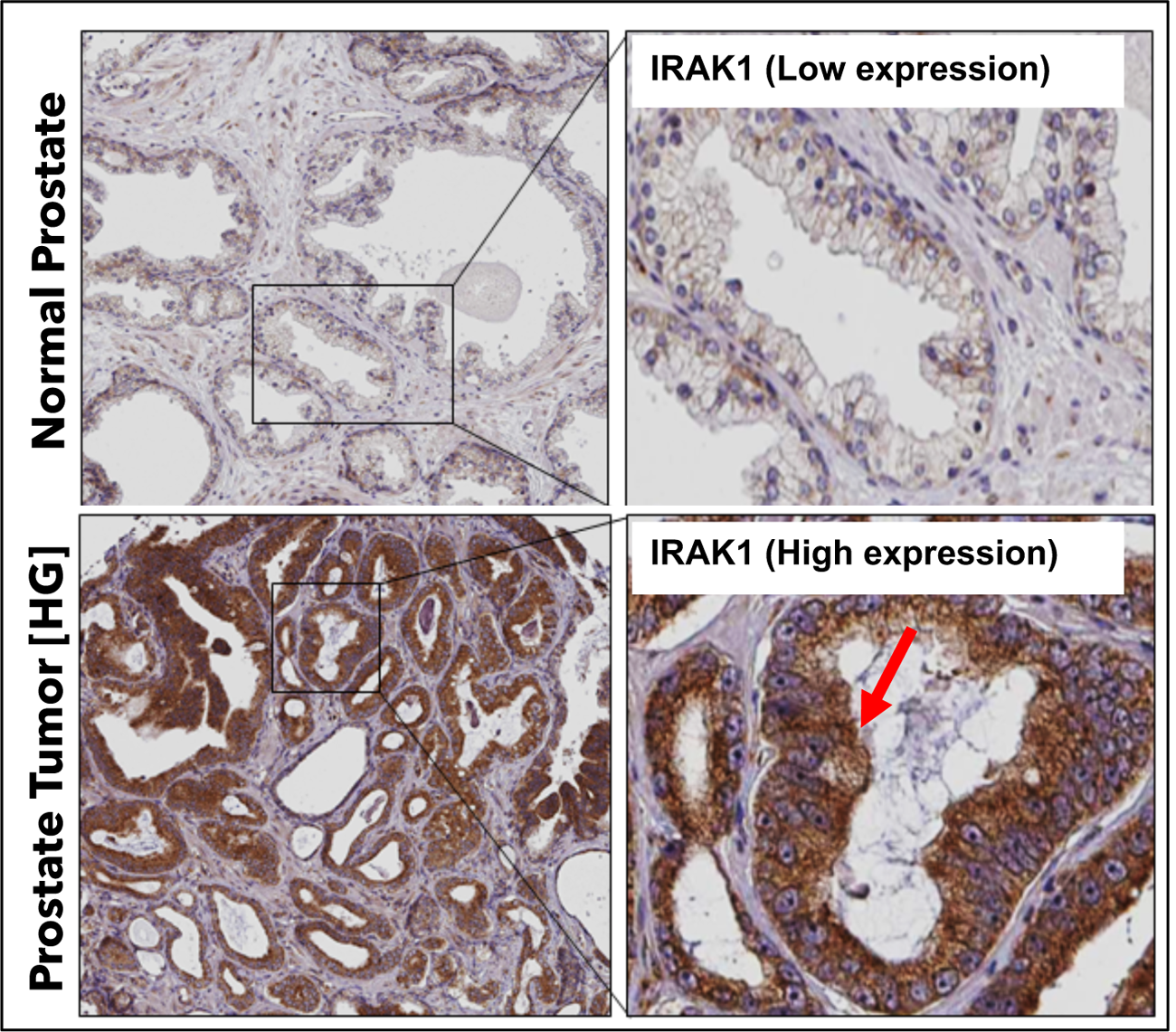
Representative Immunohistochemistry images of tissue samples from normal prostate & PCa (high-grade) stained with anti-IRAK1 antibody (brown). The nuclei were counter-stained with Hematoxylin (blue). All images were obtained from The Human Protein Atlas; http://www.proteinatlas.org. Scale bar = 100μm.

Functional enrichment analysis is an important approach for the interpretation of gene lists derived from the large-scale genome, transcriptome, and proteome studies (Zambon et al., 2012). WebGestalt and Enrichr, among others, were able to provide a platform for the identification and characterization (gene ontology, pathway, and network module analysis, gene-phenotype association, gene-disease association, gene-drug association, and chromosomal location) of the 10 inflammatory genes in the magenta module, especially the most studied inflammatory gene candidate in this study, IRAK1 (Kuleshov et al., 2016; Wang et al., 2017; Zhang et al., 2005). The differential gene analysis revealed the upregulation of IRAK1 in PCa samples compared to non-PRAD normal samples.

Variable degrees of CNAs and mutations have been identified and described in PCa (Lalonde et al., 2014; Taylor et al., 2010). Consistent with previous literature, we identified several missense and frameshift mutations of IRAK1 in PCa samples that may be of clinical significance. The functional impact of each identified mutation was predicted using COSMIC, SIFT, Mutation Assessor, and Polyphen algorithms, and found to be mutation-dependent (Flanagan et al., 2010; Gnad et al., 2013). For instance, frameshift (insertion and deletion) mutations of IRAK1 were found to be more functionally impactful or severe when compared to the missense mutations (**Table 7**).

Also, the genetic alteration frequencies of most inflammatory genes were found to be positively correlated with the aggressive status of PCa. For instance, the CRPC and NEPC samples analyzed in this study show extensive burdens of CNA and lesser somatic mutations of IRAK1 compared to indolent and low-Gleason PRAD samples (**Figure 14B** **& C**). This is in accordance with previous studies that found a link between cancer relapse and the pattern of CNA in 168 primary tumors, raising the possibility of CNA as a prognostic biomarker (Taylor et al., 2010). Another study also found CNA burden across the genome, the percentage of the tumor genome affected by CNA to be associated with PCa biochemical recurrence and metastasis after surgery in two cohort studies (Hieronymus et al., 2014). Further mechanistic studies will be needed to understand the biological and clinical significance of these mutations and CNVs in prostate tumorigenesis. Next, we identified hypomethylation lesions in IRAK1 by identifying CpG sites of IRAK1 with significant methylation values in correlation with CNVs and transcript or gene expression profiles of IRAK1 in PRAD samples.

Collectively, our integrative genomic analyses show a co-occurrence or interdependence between alterations of IRAKs and diverse genes associated with castration resistance, stemness, immunosuppression, and NED in PCa patients. IRAK1 was shown to be the most significant and frequently altered gene among the 10 inflammatory genes and IRAKs in the aggressive PCa samples. Previously, inhibition of IRAK1 and/or IRAK4 by either gene silencing or small molecule inhibitors have been shown to induce apoptosis and decrease in cell viability of lymphoblastic leukemia, hepatocellular carcinoma, melanoma, gastric cancer, pancreatic cancer, and cervical cancer (Hu et al., 2018; Li et al., 2015; Sun et al., 2006; Sun et al., 2014; Wee et al., 2015; Zhang et al., 2016). The upregulation of IRAK1 has been reported to enhance angiogenesis and metastasis of breast cancer (Singer et al., 2018). Other studies suggest that dysregulation of IRAK1 may drive tumorigenesis by activating oncogenic pathways necessary for cancer cell survival, proliferation, and metastasis (Hu et al., 2018; Li et al., 2015; Sun et al., 2014; Wee et al., 2015; Zhang et al., 2016). However, how dysregulation of IRAK1 drives prostate tumorigenesis, especially PCa progression, has not been explored.

The PI3K and its downstream kinases (i.e., AKT/PKB and mTOR) are frequently upregulated and genetically altered in CRPCs. A previous study has found PI3K, AKT, and mTOR oncogenes to be genetically altered in 42% of primary prostate tumor samples and 100% of metastatic prostate tumor samples (Bitting & Armstrong, 2013). Since PI3K signal transduction drives castration resistance and CRPC survival, the link between IRAK1-mediated inflammation signaling and PI3K/AKT/mTOR signaling needs to be clearly defined in PCa. Though our analysis revealed an association between the PI3K-AKT-mTOR and IRAKs signaling pathways, whether the dysregulation of IRAK1 in PRAD and CRPCs promotes castration resistance via the AR and PI3K-AKT-mTOR pathway has not been studied. In liver cancer, IRAK1 overexpression was found to be important for the maintenance of aggressive tumor-initiating/stem cells (Cheng et al., 2018).

Our analysis revealed a significant correlation between the co-expression/co-modification of IRAK1 and genes associated with PCa stemness, neuroendocrine, and castration resistance signaling. Since PCSCs are maintained by upregulation of the PI3K-AKT-mTOR-NF-κB, SOX2-OCT4-KLF4-MYC, Wnt-β-catenin, Notch, TGF-β, and Hedgehog signaling pathways, more mechanistic studies will be needed to elucidate and define the role of IRAK1 signaling and dysregulation in PCa stemness, neuroendocrine differentiation, immunosuppression, metabolic reprogramming, and castration resistance (Rybak et al., 2011; Rybak et al., 2015).

## Conclusion

We have identified 10 inflammatory genes associated with PCa progression and characterized IRAK1 as a potential therapeutic target in PCa. The dysregulation of IRAK1 in PRAD patients was shown to possibly have genetic and epigenetic causal effects, which justifies further studies on its impact in chronic inflammation-driven PCa progression as well as its use as a diagnostic or prognostic biomarker. DNA hypomethylation of IRAK1 was found to correlate with its overexpression in PRAD patients/samples, which suggests that the contribution of post-translational modifications such as DNA methylation plays an important role in the dysregulation of IRAK1 in PCa cells. Collectively, we have provided evidence that suggests the potential therapeutic benefits of targeting chronic inflammation in PCa as a means to prevent progression and aggressiveness of the disease in survivors.

## Data Availability

Data can be accessed from the corresponding author upon request. The codes/WGCNA script and the multivariate analysis result outputs can be found on GitHub: https://github.com/soseni2013/WGCNA-for-Cancer

## Acknowledgments

We want to thank all members of Kumi-Diaka’s lab for their insight, suggestions, discussions, and technical support. This project was partially supported by funding to Saheed Oseni through the Graduate Student Thesis and Dissertation Scholarship. We are also grateful to the Dean’s Office of FAU Charles E. Schmidt College of Science for the supplemental funding through the Science Graduate Research Scholarship and Vincent Saurino Fellowship awarded to Saheed Oseni.

## Author Contributions

SOO, MP, WA, GBF, JH, and JKD were involved in the conceptualization and study design. SOO was involved in data collection and curation. SOO, OA, AA, and AK were involved in data preprocessing and analyses. SOO wrote and drafted the original manuscript. JK, GBF, MP, and WA supervised the project. All co-authors contributed to the reviewing, editing, and final approval of the manuscript.

### List of Supplementary Materials

Supplementary Method S1: Preparing genomic and clinical data for biological network analysis

Supplementary Table S1: Enrichr gene enrichment analysis of the 10 inflammatory genes in the magenta module based on a significant p-value.

Supplementary Table S2: Functional impact ranking rubrics for IRAK1 gene mutations in PRAD samples using algorithms from different tools.

Supplementary Figure S1: Batch effect correction plots for the PRAD dataset.

Supplementary Figure S2: Hierarchical sample cluster heatmap of PRAD eigengene network.

Supplementary Figure S3: Hierarchical gene expression cluster heatmap of PRAD eigengene network.

Supplementary Figure S4: Kaplan-Meier DFS survival analysis for high-expressing vs low-expressing PRAD samples.

Supplementary Figure S5: Receiver Operating Characteristic (ROC) analysis for IRAK in predicting cancer relapse.

Supplementary Figure S6: Heatmap of the expression profile of IRAK family genes in indolent PRAD patients (n = 472) showing the overexpression of IRAK1 compared to other IRAKs.

Supplementary Figure S7: Heatmap of the expression profile of IRAK family genes in neuroendocrine PCa patients (n = 30) showing the overexpression of IRAK1 compared to other IRAKs.

Supplementary Figure S8: Heatmap of the expression profile of IRAK family genes in castration-resistant PCa patients (n = 444) showing the overexpression of IRAK1 compared to other IRAKs.

Supplementary Figure S9: Lollipop plots showing the identified structural mutations of IRAK1, IRAK2, IRAK3, and IRAK4 in PCa patients from the 20 cohort studies.

Supplementary Figure S10: Protein-protein interaction and enrichment network analyses in PRAD.

Supplementary Figure S11: Genes Word-Cloud Generator showing all the 588 genes identified in the magenta module.

Supplementary Figure S12: Diseases Word-Cloud Generator showing all the diseases associated with aberrant IRAK1 signaling.

Supplementary Figure S13: Boxplots to compare Eigengene values between progressed vs non-progressed patients in the magenta and brown modules.

Supplementary Figure S14: A flow chart summarizing the post hoc multivariate analyses undertaken for screening and identification of significant inflammatory genes in the magenta module.

Supplementary Figure S15: A principal component analysis (PCA) dimensionality reduction of IRAK1 expression in PRAD and non-PRAD normal samples.

Supplementary Figure S16. GO Elite ontologies of the 9 biologically significant modules.

Supplementary Figure S17: A detailed genomic sketch of IRAK1 showing information on the 13 transcripts/isoforms identified as well as exons, gene coding sequence regions (CDS), and untranslated regions (UTR).

Supplementary Figure S18 (A-M): Correlation analysis plots using Pearson coefficient to identify the correlation between various IRAK1 CpG methylation values and the expression levels of IRAK1 isoforms.

## References

1. Adzhubei, I., Jordan, D. M., & Sunyaev, S. R. (2013). Predicting functional effect of human missense mutations using PolyPhen-2. Current protocols in human genetics. Chapter 7, Unit 7. 20. https://doi.org/10.1002/0471142905.hg0720s76

2. Ashkani, J., Naidoo, K. (2016). Glycosyltransferase Gene Expression Profiles Classify Cancer Types and Propose Prognostic Subtypes. Sci Rep, 6, 26451, 1–8. https://doi.org/10.1038/srep26451

3. Bitting, R. L., & Armstrong, A. J. (2013). Targeting the PI3K/Akt/mTOR pathway in castration-resistant prostate cancer. Endocrine-Related Cancer, 20(3), 83–99. https://doi.org/10.1530/ERC-12-0394

4. Cerami, E., Gao, J., Dogrusoz, U., Gross, B. E., Sumer, S. O., Aksoy, B. A., Jacobsen, A., Byrne, C. J., Heuer, M. L., Larsson, E., Antipin, Y., Reva, B., Goldberg, A. P., Sander, C., & Schultz, N. (2012). The cBio cancer genomics portal: an open platform for exploring multidimensional cancer genomics data. Cancer Discovery, 2(5), 401–404. https://doi.org/10.1158/2159-8290.CD-12-0095

5. Cheng, B. Y., Lau, E. Y., Leung, H. W., Oi-Ning Leung, C., Ho, N. P., Gurung, S., Cheng, L. K., Ho Lin, C., Cheuk-Lam Lo, R., Ma, S., Oi-Lin Ng, I., Lee, T. K., & Ka Shing, L. (2018). IRAK1 augments cancer stemness and drug resistance via the AP-1/AKR1B10 signaling cascade in hepatocellular carcinoma. Cancer Res, 78(9), 2332–2342. https://doi.org/10.1158/0008-5472.CAN-17-2445

6. Cheng, S., Jiang, X., Ding, C., Du, C., Owusu-Ansah, K.G., Weng, X., Hu, W., Peng, C., Lv, Z., Tong, R., Xiao, H., Xie, H., Zhou, L., Wu, J., Zheng, S. (2016). Expression and the critical role of interleukin enhancer-binding factor 2 in hepatocellular carcinoma. Int. J. Mol. Sci., 17, 1373. https://doi.org/10.3390/ijms17081373

7. Flannery, S., & Bowie, A. G. (2010). The interleukin-1 receptor-associated kinases: Critical regulators of innate immune signaling. Biochemical Pharmacology, 80(12), 1981– 1991. https://doi.org/10.1016/J.BCP.2010.06.020

8. Flanagan, S. E., Patch, A. M., & Ellard, S. (2010). Using SIFT and PolyPhen to predict loss-of-function and gain-of-function mutations. Genetic testing and molecular biomarkers, 14(4), 533–537. https://doi.org/10.1089/gtmb.2010.0036

9. Gnad, F., Baucom, A., Mukhyala, K., Manning, G., & Zhang, Z. (2013). Assessment of computational methods for predicting the effects of missense mutations in human cancers. BMC genomics, 14 Suppl 3(Suppl 3), S7. https://doi.org/10.1186/1471-2164-14-S3-S7

10. Guo, Z., Zeng, Y., Chen, Y., Liu, M., Chen, S., Yao, M., Zhang, P., Zhong, F., Jiang, K., He, S., & Yuan, G. (2020). TRAIP promotes malignant behaviors and correlates with poor prognosis in liver cancer. Biomedicine & Pharmacotherapy, 124, 109857. https://doi.org/10.1016/j.biopha.2020.109857

11. Hasin, Y., Seldin, M. & Lusis, A. (2017). Multi-omics approaches to disease. Genome Biol, 18, 83. https://doi.org/10.1186/s13059-017-1215-1

12. Heindl, A., Nawaz, S., & Yuan, Y. (2015). Mapping spatial heterogeneity in the tumor microenvironment: A new era for digital pathology. Lab Invest., 95, 377–384. https://doi.org/10.1038/labinvest.2014.155

13. Héninger, E., Krueger, T. E. G., & Lang, J. M. (2015). Augmenting antitumor immune responses with epigenetic modifying agents. Frontiers in Immunology. 6(29), 1–14. DOI=10.3389/fimmu.2015.00029. https://doi.org/10.3389/fimmu.2015.00029

14. Hieronymus, H., Schultz, N., Gopalan, A., Carver, B. S., Chang, M. T., Xiao, Y., Heguy, A., Huberman, K., Bernstein, M., Assel, M., Murali, R., Vickers, A., Scardino, P. T., Sander, C., Reuter, V., Taylor, B. S., & Sawyers, C. L. (2014). Copy number alteration burden predicts prostate cancer relapse. Proceedings of the National Academy of Sciences of the United States of America, 111(30), 11139–11144. https://doi.org/10.1073/pnas.1411446111

15. Hu, Q., Song, J., Ding, B., Cui, Y., Liang, J., & Han, S. (2018). miR-146a promotes cervical cancer cell viability via targeting IRAK1 and TRAF6. Oncology Reports, 39(6), 3015–3024. https://doi.org/10.3892/or.2018.6391

16. Jain, A., Kaczanowska, S., Davila, E., & Research, B. (2014). IL-1 receptor-associated kinase signaling and its role in inflammation, cancer progression, and therapy resistance. Frontiers in Immunology, 5(553), 1–8. https://doi.org/10.3389/fimmu.2014.00553

17. Jang, L. K., Lee, Z. H., Kim, H. H., Hill, J. M., Kim, J. D., & Kwon, B. S. (2001). A novel leucine-rich repeat protein (LRR-1): potential involvement in 4-1BB-mediated signal transduction. Molecules and Cells, 12(3), 304–312.

18. Jianjiong Gao, Bülent Arman Aksoy, Ugur Dogrusoz, Gideon Dresdner, Benjamin Gross, S. Onur Sumer, Yichao Sun, Anders Jacobsen, Rileen Sinha, Erik Larsson, Ethan Cerami, Chris Sander, and N. S. (2012). Integrative analysis of complex cancer genomics and clinical profiles using the cBioPortal. Sci Signal, 269(6), 1–34. https://doi.org/10.1126/scisignal.2004088.Integrative

19. Koch, A., De Meyer, T., Jeschke, J., & Van Criekinge, W. (2015). MEXPRESS: Visualizing expression, DNA methylation, and clinical TCGA data. BMC Genomics, 16(636), 1–6. https://doi.org/10.1186/s12864-015-1847-z

20. Kutikhin, A. G., & Yuzhalin, A. E. (2012). Are Toll-like receptor gene polymorphisms associated with prostate cancer? Cancer Management and Research, 2012, 23–29. https://doi.org/10.2147/CMAR.S28683

21. Lalonde, E., Ishkanian, A.S., Sykes, J., Fraser, M., Ross-Adams, H., Erho, N., Dunning, M.J., Halim, S., Lamb, A.D., Moon, N.C., et al. (2014). Tumor genomic and microenvironmental heterogeneity for integrated prediction of 5-year biochemical recurrence of prostate cancer: a retrospective cohort study. Lancet Oncol. 15, 1521– 1532.

22. Langfelder, P., & Horvath, S. (2008). WGCNA: An R package for weighted correlation network analysis. BMC Bioinformatics, 9, 559. https://doi.org/10.1186/1471-2105-9-559

23. Li, Z., Younger, K., Gartenhaus, R., Joseph, A. M., Hu, F., Baer, M. R., Brown, P., & Davila, E. (2015). Inhibition of IRAK1/4 sensitizes T cell acute lymphoblastic leukemia to chemotherapies. Journal of Clinical Investigation, 125(3), 1081–1097. https://doi.org/10.1172/JCI75821

24. Liu, Q., Harvey, C. T., Geng, H., Xue, C., Chen, V., Beer, T. M., & Qian, D. Z. (2013). Malate dehydrogenase 2 confers docetaxel resistance via regulations of JNK signaling and oxidative metabolism. The Prostate, 73(10), 1028–1037. https://doi.org/10.1002/pros.22650

25. Liu, Y., Dong, Y., Zhao, L., Su, L., Diao, K., & Mi, X. (2018). TRIM59 overexpression correlates with poor prognosis and contributes to breast cancer progression through the AKT signaling pathway. Molecular carcinogenesis, 57(12), 1792–1802. https://doi.org/10.1002/mc.22897

26. Mandell, M. A., Saha, B., & Thompson, T. A. (2020). The tripartite nexus: Autophagy, cancer, and tripartite motif-containing protein family members. Frontiers in pharmacology, 11, 308. https://doi.org/10.3389/fphar.2020.00308

27. Nagy Á. & Győrffy B. (2021). muTarget: A platform linking gene expression changes and mutation status in solid tumors, International Journal of Cancer,148:502–511

28. Ng, P. C. & Henikoff, S. (2003). SIFT: Predicting amino acid changes that affect protein function. Nucleic acids research, 31(13), 3812–3814. https://doi.org/10.1093/nar/gkg509

29. Ohandjo, A. Q., Liu, Z., Dammer, E. B., Dill, C. D., Griffen, T. L., Carey, K. M., Hinton, D. E., Meller, R., & Lillard, J. W., Jr (2019). Transcriptome network analysis identifies CXCL13-CXCR5 signaling modules in the prostate tumor immune microenvironment. Scientific reports, 9(1), 14963. https://doi.org/10.1038/s41598-019-46491-3

30. Pandey, S., Singh, S., Anang, V., Bhatt, A. N., Natarajan, K., & Dwarakanath, B. S. (2015). Pattern recognition receptors in cancer progression and metastasis. Cancer growth and metastasis, 8, 1–10. https://doi.org/10.4137/CGM.s24314

31. Pezaro, C. J., Omlin, A. G., Altavilla, A., Lorente, D., Ferraldeschi, R., Bianchini, D., Dearnaley, D., Parker, C., de Bono, J. S., & Attard, G. (2014). Activity of Cabazitaxel in castration-resistant prostate cancer progressing after docetaxel and next-generation endocrine agents. European Urology, 66(3), 459–465. https://doi.org/10.1016/J.EURURO.2013.11.044

32. Queen, D., Ediriweera, C., & Liu, L. (2019). Function and Regulation of IL-36 Signaling in Inflammatory Diseases and Cancer Development. Frontiers in cell and developmental biology, 7, 317. https://doi.org/10.3389/fcell.2019.00317

33. R Core Team (2020). R: A language and environment for statistical computing. R Foundation for Statistical Computing, Vienna, Austria. https://www.R-project.org/

34. Robinson, D., Van Allen, E. M., Wu, Y.-M., Schultz, N., Lonigro, R. J., Mosquera, J. M., Montgomery, B., Taplin, M.-E., Pritchard, C. C., Attard, G., Beltran, H., Abida, W., Bradley, R. K., Vinson, J., Cao, X., Vats, P., Kunju, L. P., Hussain, M., Feng, F. Y., Chinnaiyan, A. M. (2015). Integrative clinical genomics of advanced prostate cancer. Cell, 161(5), 1215– 1228. https://doi.org/10.1016/j.cell.2015.05.001

35. Rothschild, D. E., McDaniel, D. K., Ringel-Scaia, V. M., & Allen, I. C. (2018). Modulating inflammation through the negative regulation of NF-κB signaling. Journal of Leukocyte Biology, 103(6), 1131–1150. https://doi.org/10.1002/JLB.3MIR0817-346RRR

36. Rybak, A. P., Bristow, R. G., & Kapoor, A. (2015). Prostate cancer stem cells: deciphering the origins and pathways involved in prostate tumorigenesis and aggression. Oncotarget, 6(4), 1900–1919. https://doi.org/10.18632/oncotarget.2953

37. Rybak, A. P., He, L., Kapoor, A., Cutz, J. C., & Tang, D. (2011). Characterization of sphere-propagating cells with stem-like properties from DU145 prostate cancer cells. Biochimica et Biophysica Acta (BBA) - Molecular Cell Research, 1813(5), 683–694. https://doi.org/10.1016/j.bbamcr.2011.01.018

38. Schiller, D. S., & Parikh, A. (2011). Identification, pharmacologic considerations, and management of prostatitis. American Journal Geriatric Pharmacotherapy, 9(1), 37–48. https://doi.org/10.1016/j.amjopharm.2011.02.005

39. Sciarra, A., Mariotti, G., Salciccia, S., Gomez, A. A., Monti, S., Toscano, V., & Di Silverio, F. (2008). Prostate growth and inflammation. The Journal of Steroid Biochemistry and Molecular Biology, 108(3–5), 254–260. https://doi.org/10.1016/j.jsbmb.2007.09.013

40. Siegel, RL, Miller, KD, Fuchs, H, Jemal, A. (2021). Cancer Statistics, 2021. CA: A Cancer Journal for Clinicians, 7, 7–33. http://onlinelibrary.wiley.com/journal/10.3322/ (ISSN)1542-4863/issues

41. Singer, J. W., Fleischman, A., Al-Fayoumi, S., Mascarenhas, J. O., Yu, Q., & Agarwal, A. (2018). Inhibition of interleukin-1 receptor-associated kinase 1 (IRAK1) as a therapeutic strategy. Oncotarget, 9(70), 33416–33439. https://doi.org/10.18632/oncotarget.26058

42. Sim, N. L., Kumar, P., Hu, J., Henikoff, S., Schneider, G., & Ng, P. C. (2012). SIFT web server: predicting effects of amino acid substitutions on proteins. Nucleic acids research, 40(Web Server issue), W452–W457. https://doi.org/10.1093/nar/gks539

43. Sondka, Z., Bamford, S., Cole, C. G., Ward, S. A., Dunham, I., Forbes, S. A. (2018). The COSMIC Cancer Gene Census: describing genetic dysfunction across all human cancers. Nat Rev Cancer, 18, 696–705. https://doi.org/10.1038/s41568-018-0060-1

44. Sun, J., Wiklund, F., Hsu, F. C., Bälter, K., Zheng, S. L., Johansson, J. E., Chang, B., Liu, W., Li, T., Turner, A. R., Li, L., Li, G., Adami, H. O., Isaacs, W. B., Xu, J., & Grönberg, H. (2006). Interactions of sequence variants in interleukin-1 receptor-associated kinase4 and the Toll-like receptor 6-1-10 gene cluster increase prostate cancer risk. Cancer Epidemiology Biomarkers and Prevention, 15(3), 480–485. https://doi.org/10.1158/1055-9965.EPI-05-0645

45. Sun, M., Yang, P., Du, L., Yang, Y., & Ye, J. (2014). The role of Interleukin-1 receptor-associated kinases in Vogt-Koyanagi-Harada disease. PLoS ONE, 9(4), e93214. https://doi.org/10.1371/journal.pone.0093214

46. Tan, J., Jin, X., & Wang, K. (2019). Integrated bioinformatics analysis of potential biomarkers for prostate cancer. Pathology oncology research: POR, 25(2), 455–460. https://doi.org/10.1007/s12253-017-0346-8

47. Tang, Z., Li, C., Kang, B., Gao, G., Li, C., & Zhang, Z. (2017). GEPIA: a web server for cancer and normal gene expression profiling and interactive analyses. Nucleic acids research, 45(W1), W98–W102. https://doi.org/10.1093/nar/gkx247

48. Taylor, B.S., Schultz, N., Hieronymus, H., Gopalan, A., Xiao, Y., Carver, B.S., Arora, V.K., Kaushik, P., Cerami, E., Reva, B., et al. (2010). Integrative genomic profiling of human prostate cancer. Cancer cell, 18, 11–22.

49. Uhlén, M., Björling, E., Agaton, C., Szigyarto, C. A., Amini, B., Andersen, E., Andersson, A. C., Angelidou, P., Asplund, A., Asplund, C., Berglund, L., Bergström, K., Brumer, H., Cerjan, D., Ekström, M., Elobeid, A., Eriksson, C., Fagerberg, L., Falk, R., Fall, J., Forsberg, M., Björklund MG, Gumbel K, Halimi, A., Hallin, I., Hamsten, C., Hansson, M., Hedhammar, M., Hercules, G., Kampf, C., Larsson, K., Lindskog, M., Lodewyckx, W., Lund J, Lundeberg, J., Magnusson, K., Malm, E., Nilsson, P., Odling, J., Oksvold, P., Olsson, I., Oster, E., Ottosson, J., Paavilainen, L., Persson, A., Rimini, R., Rockberg, J., Runeson, M., Sivertsson, A., Sköllermo, A., Steen, J., Stenvall, M., Sterky, F., Strömberg, S., Sundberg, M., Tegel, H., Tourle, S., Wahlund, E., Waldén, A., Wan, J., Wernérus, H., Westberg, J., Wester, K., Wrethagen, U., Xu, L. L., Hober, S., Pontén, F. (2005). A human protein atlas for normal and cancer tissues based on antibody proteomics. Mol Cell Proteomics, 4(12):1920–32. PubMed: 16127175 DOI: 10.1074/mcp.M500279-MCP200

50. Wagenlehner, F. M. E., Elkahwaji, J. E., Algaba, F., Bjerklund-Johansen, T., Naber, K. G., Hartung, R., & Weidner, W. (2007). The role of inflammation and infection in the pathogenesis of prostate carcinoma. BJU International, 100(4), 733–737. https://doi.org/10.1111/j.1464-410X.2007.07091.x

51. Walsh, P. T., & Fallon, P. G. (2018). The emergence of the IL-36 cytokine family as novel targets for inflammatory diseases. Annals of the New York Academy of Sciences, 1417(1), 23–34. https://doi.org/10.1111/nyas.13280

52. Wee, Z. N., Yatim, S. M. J. M., Kohlbauer, V. K., Feng, M., Goh, J. Y., Yi, B., Lee, P. L., Zhang, S., Wang, P. P., Lim, E., Tam, W. L., Cai, Y., Ditzel, H. J., Hoon, D. S. B., Tan, E. Y., & Yu, Q. (2015). IRAK1 is a therapeutic target that drives breast cancer metastasis and resistance to paclitaxel. Nature Communications, 6, 8746. https://doi.org/10.1038/ncomms9746

53. Weinstein, J. N., Collisson, E. A., Mills, G. B., Mills Shaw, K. R., Ozenberger, B. A., Ellrott, K., Shmulevich, I., Sander, C., & Stuart, J. M. (2013). The Cancer Genome Atlas Pan-Cancer analysis project. Nature Genetics, 45, 1113–1120 https://doi.org/10.1038/ng.2764

54. Wodarz, D., Sorace, R., & Komarova, N. L. (2014). Dynamics of Cellular Responses to Radiation. PLoS Computational Biology, 10(4), e1003513. https://doi.org/10.1371/journal.pcbi.1003513

55. Xu W. X., Zhang, J., Hua, Y. T., Yang, S. J., Wang, D. D., Tang, J. H. (2020). An integrative pan-cancer analysis revealing LCN2 as an oncogenic immune protein in the tumor microenvironment. Frontiers in Oncology, 10, 2798. DOI=10.3389/fonc.2020.605097. https://www.frontiersin.org/article/10.3389/fonc.2020.605097

56. Xu, X., Zhang, X., Xing, H., Liu, Z., Jia, J., Jin, C., Zhang, Y. (2019). Importin-4 functions as a driving force in human primary gastric cancer. J Cell Biochem, 120, 12638–12646. https://doi.org/10.1002/jcb.28530

57. Yang, Y., Dong, X., Xie, B., Ding, N., Chen, J., Li, Y., Zhang, Q., Qu, H., & Fang, X. (2015). Databases and web tools for cancer genomics study. Genomics, Proteomics & Bioinformatics, 13(1), 46–50. https://doi.org/10.1016/j.gpb.2015.01.005

58. Yeh, D. W., Huang, L. R., Chen, Y. W., Huang, C. Y. F., & Chuang, T. H. (2016). Interplay between inflammation and stemness in cancer cells: the role of toll-like receptor signaling. Journal of Immunology Research, 2016, 1–14. https://doi.org/10.1155/2016/4368101

59. Yin, Z. H., Jiang, X. W., Shi, W. B., Gui, Q. L., & Yu, D. F. (2017). Expression and Clinical Significance of ILF2 in Gastric Cancer. Disease markers, 2017, 4387081. https://doi.org/10.1155/2017/4387081

60. Yu, B., Ding, Y., Liao, X., Wang, C., Wang, B., & Chen, X. (2019). Overexpression of TONSL might be an independent unfavorable prognostic indicator in hepatocellular carcinoma. Pathology, research, and practice, 215(5), 939–945. https://doi.org/10.1016/j.prp.2019.01.044

61. Zhang, D., Li, L., Jiang, H., Knolhoff, B. L., Lockhart, A. C., Wang-Gillam, A., Denardo, D. G., Ruzinova, M. B., & Lim, K.-H. (2016). Constitutive IRAK4 activation underlies poor prognosis and chemoresistance in pancreatic ductal adenocarcinoma. Clinical Cancer Research, 23(7), 1748–59. https://doi.org/10.1158/1078-0432.CCR-16-1121

62. Zhou, Y., Liu, F., Xu, Q., Yang, B., Li, X., Jiang, S., Hu, L., Zhang, X., Zhu, L., Zhu, X., Shao, H., Dai, M., Shen, Y., Ni, B., Wang, S., Zhang, Z., Teng, Y. (2020). Inhibiting Importin 4-mediated nuclear import of CEBPD enhances chemosensitivity by repression of PRKDC-driven DNA damage repair in cervical cancer. Oncogene, 39, 5633–5648 https://doi.org/10.1038/s41388-020-1384-3

63. Zhu S., Jin J., Gokhale S., Lu A. M., Shan H., Feng J., Xie P. (2018). Genetic alterations of TRAF proteins in human cancers. Frontiers in Immunology, 9, 2111. 10.3389/fimmu.2018.02111. https://www.frontiersin.org/article/10.3389/fimmu.2018.02111

