## Supplementary Materials for "Integrative genomic and epigenomic analyses identified IRAK1 as a novel target for chronic inflammation-driven prostate tumorigenesis"

**List of Supplementary Materials**

Supplementary Method S1: Preparing genomic and clinical data for biological network analysis

Supplementary Method S2: WGCNA R packages for biological network analysis

Supplementary Table S1: Enrichr gene enrichment analysis of the 10 inflammatory genes in the magenta module based on a significant p-value.

Supplementary Table S2: Functional impact ranking rubrics for IRAK1 gene mutations in PRAD samples using algorithms from different tools.

Supplementary Figure S1: Batch effect correction plots for the PRAD dataset.

Supplementary Figure S2: Hierarchical sample cluster heatmap of PRAD eigengene network.

Supplementary Figure S3: Hierarchical gene expression cluster heatmap of PRAD eigengene network.

Supplementary Figure S4: Kaplan-Meier DFS survival analysis for high-expressing vs low-expressing PRAD samples.

Supplementary Figure S5: Receiver Operating Characteristic (ROC) analysis for IRAK in predicting cancer relapse.

Supplementary Figure S6: Heatmap of the expression profile of IRAK family genes in indolent PRAD patients (n = 472) showing the overexpression of IRAK1 compared to other IRAKs.

Supplementary Figure S7: Heatmap of the expression profile of IRAK family genes in neuroendocrine PCa patients (n = 30) showing the overexpression of IRAK1 compared to other IRAKs.

Supplementary Figure S8: Heatmap of the expression profile of IRAK family genes in castration-resistant PCa patients (n = 444) showing the overexpression of IRAK1 compared to other IRAKs.

Supplementary Figure S9: Lollipop plots showing the identified structural mutations of IRAK1, IRAK2, IRAK3, and IRAK4 in PCa patients from the 20 cohort studies.

Supplementary Figure S10: Protein-protein interaction and enrichment network analyses in PRAD.

Supplementary Figure S11: Genes Word-Cloud Generator showing all the 588 genes identified in the magenta module.

Supplementary Figure S12: Diseases Word-Cloud Generator showing all the diseases associated with aberrant IRAK1 signaling.

Supplementary Figure S13: Boxplots to compare Eigengene values between progressed vs non-progressed patients in the magenta and brown modules.

Supplementary Figure S14: A flow chart summarizing the post hoc multivariate analyses undertaken for screening and identification of significant inflammatory genes in the magenta module.

Supplementary Figure S15: A principal component analysis (PCA) dimensionality reduction of IRAK1 expression in PRAD and non-PRAD normal samples.

Supplementary Figure S16. GO Elite ontologies of the 9 biologically significant modules.

Supplementary Figure S17: A detailed genomic sketch of IRAK1 showing information on the 13 transcripts/isoforms identified as well as exons, gene coding sequence regions (CDS), and untranslated regions (UTR).

Supplementary Figure S18 (A-M): Correlation analysis plots using Pearson coefficient to identify the correlation between various IRAK1 CpG methylation values and the expression levels of IRAK1 isoforms.

**Supplementary Method S1:** Preparing clinical and genomic datasets for biological network analysis. The non-numeric clinical data were converted to numeric and binary units as follows:

Tumor Type: “0” -> “PRAD Acinar-type” or “1” -> “Other PRAD Subtype”.

PFS_Status: “0” -> “Remission (Non-progressed)” and “1” -> “Progressed/Recurred”.

OS_Status/Vital Status: “0” -> “Dead” or “1” -> “Living”.

DS_Status: “0” -> “Disease-free”; and “1” -> “Recurred”.

Race: “1” -> “White American”; “2” -> “African American”; or “3” -> “Asian American”.

Prior Radiation Therapy Status: “0”-> “No” or “1” -> “Yes”.

Biochemical Recurrence: “0”-> “No” or “1”-> “Yes”.

Gleason Score: “0” -> “G0” or “T0”; “1” -> “G6” or “T1/T2a/T2b/T2c”; “2” -> “G7 or “T3a/T3b”; and “3” -> “G8 or “T4”.

Tumor (T) Staging Pathology: “0” -> “T0”; “1” -> “T1”; “2” -> “T2a”; “3” -> “T2b”; “4” -> “T2c”; “5” -> “T3a”; “6” -> “T3b” or “7” -> “T4”.

Nearby Lymph Node (N) Staging: “0” -> “N0” or “1” -> “N1”.

Distant Metastasis (M) Staging: “0” -> “M0”; “1” -> “M1”; “2” -> “M1a”; “3” -> “M1b”; or “4” -> “M1c”.

DFS_Status: “0” -> “Alive or Dead with tumor-free” or “1” -> “Dead or Alive with tumor”.

Cancer Status: “0” -> “Tumor-free” or “1” -> “With tumor”.

Biochemical Recurrence: “0” -> No or “1” -> “Yes”.

PFS_Months: “1” -> “Above cut-off: If > 25.8 (median) months” or “0 -> “Below cut-off, if < 25.8 (median) months”.

DFS_Months: “1” -> “Above cut-off, If > 30.35 (median) months” or “0” -> “Below cut-off, if < 30.4 (median) months”.

OS_Months: “0” -> “Below cut-off, if < 30.4 (median) months”; or “1” -> “Above cut-off: If > 30.4 (median) months”.

DS_Months: “1” -> “Above cut-off: If > 30.4 (median) months”; or “0” -> “Below median cut-off, if < 30.4 (median) months”.

Acronyms: OS: Overall survival; PFS: Progression-free survival; DFS: Disease-free survival; DS: Disease status

**Supplementary Method S2:** WGCNA R packages for biological network analysis.

The WGCNA package was installed from the Comprehensive R Archive Network (CRAN), the standard repository for R add-on packages. To run the WGCNA package, the installation tools/algorithms for the following packages were obtained from Bioconductor and installed in R: “GO.db”, “dynamicTreeCut”, “doParallel”, “parallel”, “stats”, “foreach”, “AnnotationDbi”, “impute”, “utils”, “splines”, “matrixStats”, “Hmisc”, “fastcluster”, “survival”, “grDevices”, and “preprocessCore”. Other supportive packages used while running the WGCNA pipeline include “NMF”, “igraph”, “ggplot2”, “RColorBrewer”, “Cairo”, "doParallel", "biomaRt", "plotly", "stringr", “boot”, and "callr".

**Supplementary Table S1:** Enrichr gene enrichment analysis of the 10 inflammatory genes in the magenta module based on a significant p-value. The first column is showing their enriched gene ontology (biological processes), the second column is the p-value (< 0.05) while the last column indicates the genes of the 10 inflammatory genes involved in the biological processes. Of the 10 genes, IRAK1 and HMGB2 have the most biologically significant functions ($p < 0.05$).

| Gene Ontologies | P-values | Genes |
|---|---|---|
| Positive regulation of megakaryocyte differentiation (GO:0045654) | 0.0038941 | HMGB2 |
| Regulation of nuclease activity (GO:0032069) | 0.0038941 | HMGB2 |
| V(D)J recombination (GO:0033151) | 0.00454175 | HMGB2 |
| DNA topological change (GO:0006265) | 0.0058359 | HMGB2 |
| Positive regulation of nuclease activity (GO:0032075) | 0.0064824 | HMGB2 |
| DNA ligation involved in DNA repair (GO:0051103) | 0.00777422 | HMGB2 |
| Apoptotic DNA fragmentation (GO:0006309) | 0.00970904 | HMGB2 |
| Regulation of stem cell proliferation (GO:0072091) | 0.01035321 | HMGB2 |
| DNA ligation (GO:0006266) | 0.01164039 | HMGB2 |
| Regulation of cell development (GO:0060284) | 0.01292602 | HMGB2 |
| Positive regulation of erythrocyte differentiation (GO:0045648) | 0.01421011 | HMGB2 |
| DNA catabolic process, endonucleolytic (GO:0000737) | 0.01421011 | HMGB2 |
| Apoptotic nuclear changes (GO:0030262) | 0.01485157 | HMGB2 |

| | | |
|---|---|---|
| Regulation of nervous system development (GO:0051960) | 0.01613335 | HMGB2 |
| Positive regulation of myeloid cell differentiation (GO:0045639) | 0.01869229 | HMGB2 |
| Regulation of erythrocyte differentiation (GO:0045646) | 0.02060746 | HMGB2 |
| Positive regulation of DNA binding (GO:0043388) | 0.02188232 | HMGB2 |
| Regulation of megakaryocyte differentiation (GO:0045652) | 0.03139508 | HMGB2 |
| Regulation of DNA binding (GO:0051101) | 0.03328735 | HMGB2 |
| Regulation of neurogenesis (GO:0050767) | 0.03643356 | HMGB2 |
| Cell chemotaxis (GO:0060326) | 0.03831674 | HMGB2 |
| Nucleosome assembly (GO:0006334) | 0.03831674 | HMGB2 |
| Defense response to Gram-positive bacterium (GO:0050830) | 0.04144784 | HMGB2 |
| Chromatin assembly (GO:0031497) | 0.04144784 | HMGB2 |
| Positive regulation of endothelial cell proliferation (GO:0001938) | 0.04269764 | HMGB2 |
| Defense response to Gram-negative bacterium (GO:0050829) | 0.04519273 | HMGB2 |
| DNA conformation change (GO:0071103) | 5.93e-05 | HMGB2 HMGB3 |
| DNA geometric change (GO:0032392) | 1.07e-04 | HMGB2 HMGB3 |
| DNA metabolic process (GO:0006259) | 0.01709624 | HMGB2 HMGB3 |
| Chromatin remodeling (GO:0006338) | 0.00249491 | HMGB2, HMGB3 |
| DNA recombination (GO:0006310) | 0.03517621 | HMGB3 |
| Negative regulation of interleukin-17 production (GO:0032700) | 0.0058359 | IL36RN |
| Regulation of interferon-gamma secretion (GO:1902713) | 0.0058359 | IL36RN |
| Negative regulation of response to cytokine stimulus (GO:0060761) | 0.0064824 | IL36RN |
| Defense response to fungus (GO:0050832) | 0.01099699 | IL36RN |
| Negative regulation of interferon-gamma production (GO:0032689) | 0.01292602 | IL36RN |
| Regulation of interleukin-17 production (GO:0032660) | 0.01356826 | IL36RN |
| Negative regulation of cytokine secretion (GO:0050710) | 0.02569772 | IL36RN |
| Negative regulation of cytokine-mediated signaling pathway (GO:0001960) | 3.00e-04 | IL36RN, TRAIP |
| Nucleic acid-templated transcription (GO:0097659) | 0.04830315 | ILF2 |
| Toll-like receptor 9 signaling pathway (GO:0034162) | 0.00841955 | IRAK1 |
| Toll-like receptor 4 signaling pathway (GO:0034142) | 0.00970904 | IRAK1 |

| | | |
|---|---|---|
| Lipopolysaccharide-mediated signaling pathway (GO:0031663) | 0.01035321 | IRAK1 |
| Activation of NF-kappaB-inducing kinase activity (GO:0007250) | 0.01035321 | IRAK1 |
| Regulation of response to cytokine stimulus (GO:0060759) | 0.01292602 | IRAK1 |
| Cytoplasmic pattern recognition receptor signaling pathway (GO:0002753) | 0.01549266 | IRAK1 |
| Nucleotide-binding domain, leucine-rich repeat-containing receptor signaling pathway (GO:0035872) | 0.01613335 | IRAK1 |
| Nucleotide-binding oligomerization domain-containing signaling pathway (GO:0070423) | 0.01805313 | IRAK1 |
| Myd88-dependent toll-like receptor signaling pathway (GO:0002755) | 0.01869229 | IRAK1 |
| Positive regulation of NIK/NF-kappaB signaling (GO:1901224) | 0.02251918 | IRAK1 |
| Pattern recognition receptor signaling pathway (GO:0002221) | 0.03076356 | IRAK1 |
| Positive regulation of type I interferon production (GO:0032481) | 0.03957031 | IRAK1 |
| Stress-activated MAPK cascade (GO:0051403) | 0.03957031 | IRAK1 |
| Cellular response to type I interferon (GO:0071357) | 0.04144784 | IRAK1 |
| Type I interferon signaling pathway (GO:0060337) | 0.04144784 | IRAK1 |
| JNK cascade (GO:0007254) | 0.04207293 | IRAK1 |
| Response to interleukin-1 (GO:0070555) | 0.04830315 | IRAK1 |
| Cellular response to lipopolysaccharide (GO:0071222) | 0.00154582 | IRAK1, HMGB2 |
| Response to a molecule of bacterial origin (GO:0002237) | 0.00178962 | IRAK1, HMGB2 |
| Response to lipopolysaccharide (GO:0032496) | 0.00440096 | IRAK1, HMGB2 |
| Positive regulation of gene expression (GO:0010628) | 0.01221852 | IRAK1, HMGB2, ILF2 |
| Positive regulation of transcription, DNA-templated (GO:0045893) | 0.03284618 | IRAK1, HMGB2, ILF2 |
| Regulation of cytokine-mediated signaling pathway (GO:0001959) | 0.0015124 | IRAK1, IL36RN |
| Cellular response to cytokine stimulus (GO:0071345) | 0.03426669 | IRAK1, IL36RN |
| Regulation of I-kappaB kinase/NF-kappaB signaling (GO:0043122) | 0.00749946 | IRAK1, TRIM59 |
| Response to lipid (GO:0033993) | 0.00360765 | IRAK1, HMGB2 |

| | | |
|---|---|---|
| Regulation of transcription, DNA-templated (GO:0006355) | 0.01616214 | IRAK1, HMGB2, HMGB3, ILF2 |
| Positive regulation of nucleic acid-templated transcription (GO:1903508) | 0.00372615 | IRAK1, HMGB2, ILF2 |
| Cytoplasmic sequestering of protein (GO:0051220) | 0.00970904 | TONSL |
| Cytoplasmic sequestering of transcription factor (GO:0042994) | 0.01035321 | TONSL |
| Negative regulation of transcription factor imports into the nucleus (GO:0042992) | 0.01485157 | TONSL |
| Replication fork processing (GO:0031297) | 0.01933106 | TONSL |
| DNA-dependent DNA replication maintenance of fidelity (GO:0045005) | 0.02251918 | TONSL |
| Recombinational repair (GO:0000725) | 0.04519273 | TONSL |
| Double-strand break repair via homologous recombination (GO:0000724) | 0.04830315 | TONSL |
| Activation of MAPKKK activity (GO:0000185) | 0.00454175 | TRAF7 |
| Regulation of apoptotic signaling pathway (GO:2001233) | 0.01613335 | TRAF7 |
| Positive regulation of apoptotic signaling pathway (GO:2001235) | 0.04830315 | TRAF7 |
| Positive regulation of MAP kinase activity (GO:0043406) | 0.00569748 | TRAF7, IRAK1 |
| Activation of protein kinase activity (GO:0032147) | 0.00968586 | TRAF7, IRAK1 |
| Positive regulation of MAPK cascade (GO:0043410) | 0.01461125 | TRAF7, IRAK1 |
| Positive regulation of intracellular signal transduction (GO:1902533) | 0.03749848 | TRAF7, IRAK1 |
| Apoptotic process (GO:0006915) | 0.00952691 | TRAF7, TRAIP |
| Negative regulation of tumor necrosis factor-mediated signaling pathway (GO:0010804) | 0.01099699 | TRAIP |
| Regulation of tumor necrosis factor-mediated signaling pathway (GO:0010803) | 0.03580507 | TRAIP |
| Negative regulation of I-kappaB kinase/NF-kappaB signaling (GO:0043124) | 0.03013166 | TRIM59 |

**Supplementary Table S2:** Functional impact ranking rubrics for IRAK1 gene mutations in PRAD samples using algorithms from different tools.

| Analysis Tools | Functional Impact Score | | | |
|---|---|---|---|---|
| Mutation Assessor | Neutral | Low | Medium | High |
| SIFT | Tolerated | Tolerated low_confidence | Deleterious low_confidence | Deleterious |
| Polyphen | Benign | | Possibly damaging | Probably damaging |
| COSMIC | Non-recurrent | | Recurrent | |
| Integrated Method | Mild | | Moderate | Severe |

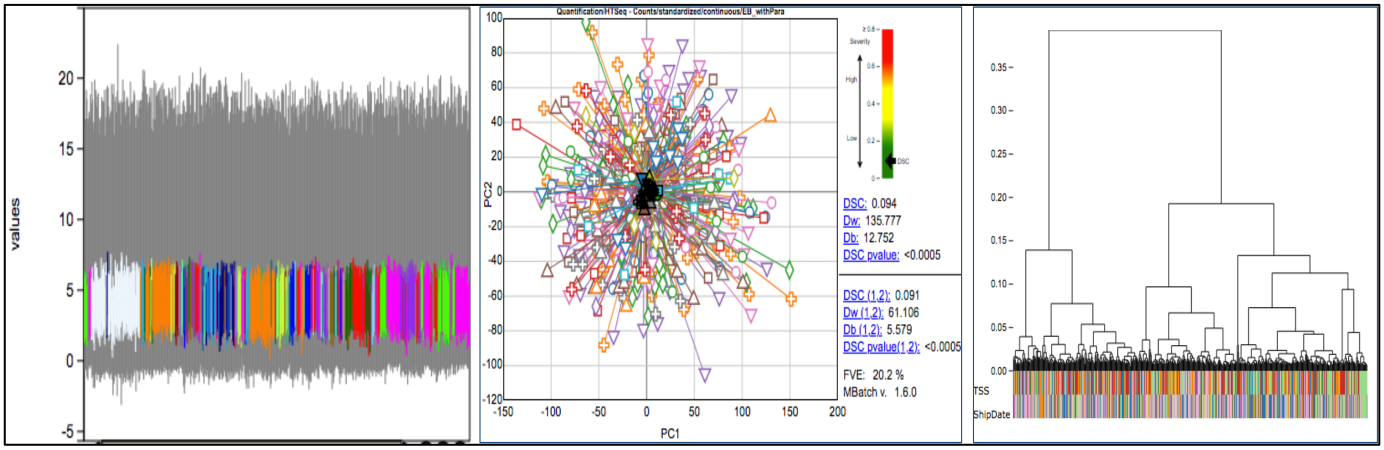


**Supplementary Figure S1:** Batch effect correction plots for the PRAD dataset. The boxplot (Left Panel), PCA (Middle Panel), and hierarchical clustering (Right Panel) plots showed uniform and less dispersion or variance between samples after preprocessing analysis using MBatch, ComBat, and TAMPOR to remove outliers. The hierarchical clustering, principal component analysis (PCA), and boxplot outputs from the batch effect viewer following the preprocessing indicate homogeneity between samples and a lower dispersion and variance  (severity) between the samples based on a DSC value of less than 0.5 (p < 0.0005) (middle panel).

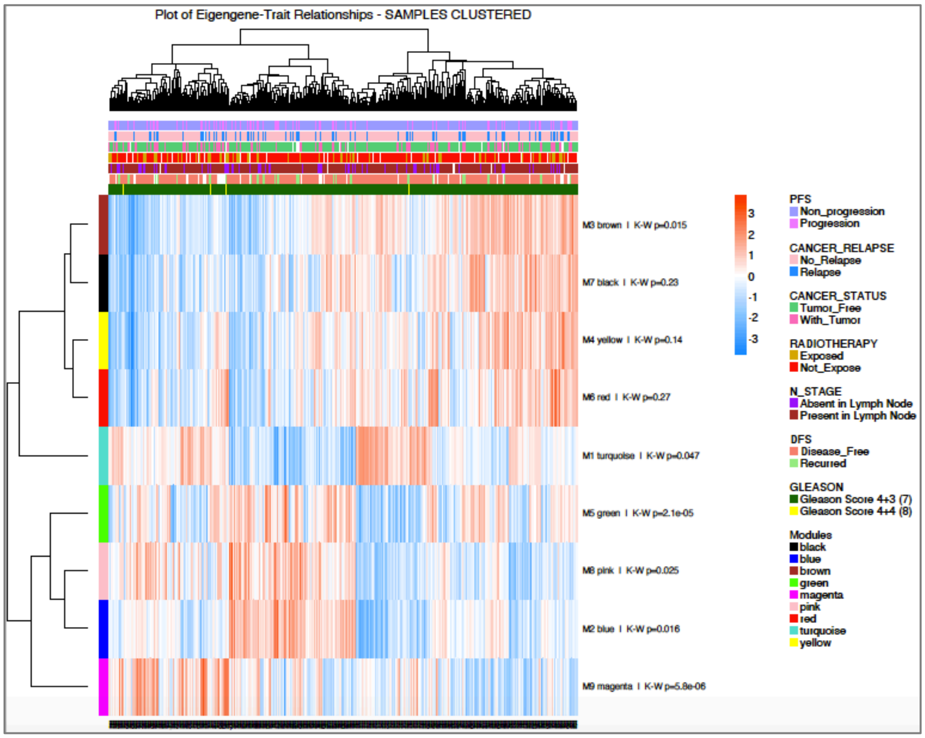


**Supplementary Figure S2:** Hierarchical sample cluster heatmap of PRAD eigengene network. The heatmap displays the hierarchical clustering of Z-scaled module eigengenes. Clusters are solely based on the relationship of eigengene to each sample.

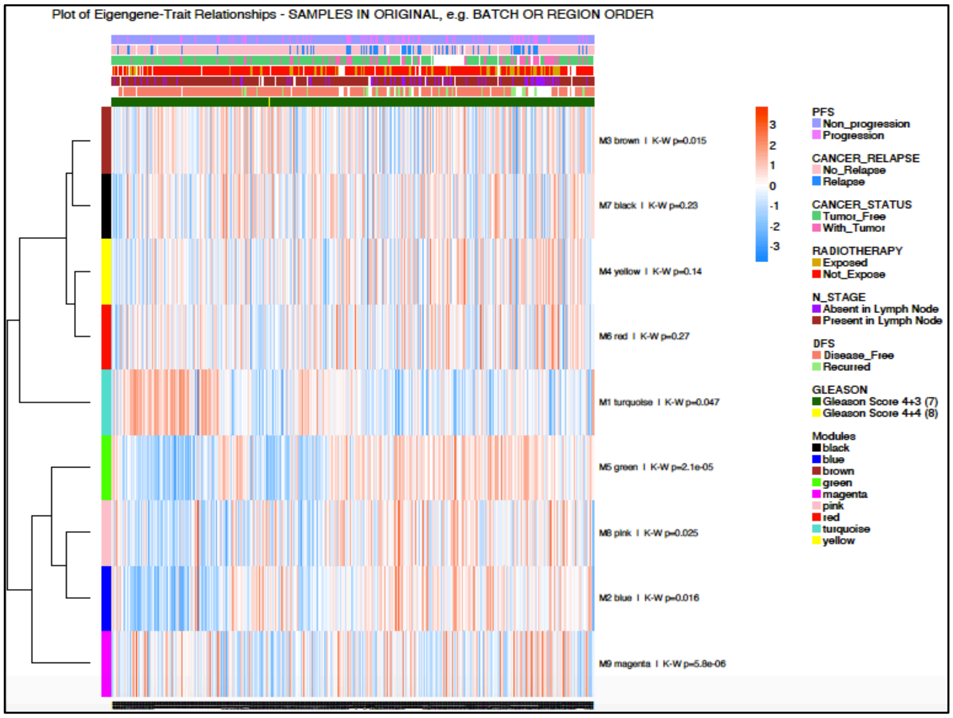

**Supplementary Figure S3:** Hierarchical gene expression cluster heatmap of PRAD eigengene network. The heatmap displays the hierarchical clustering of Z-scaled module eigengenes. Clusters are solely based on the distinct heterogeneous expression profile of each patient.

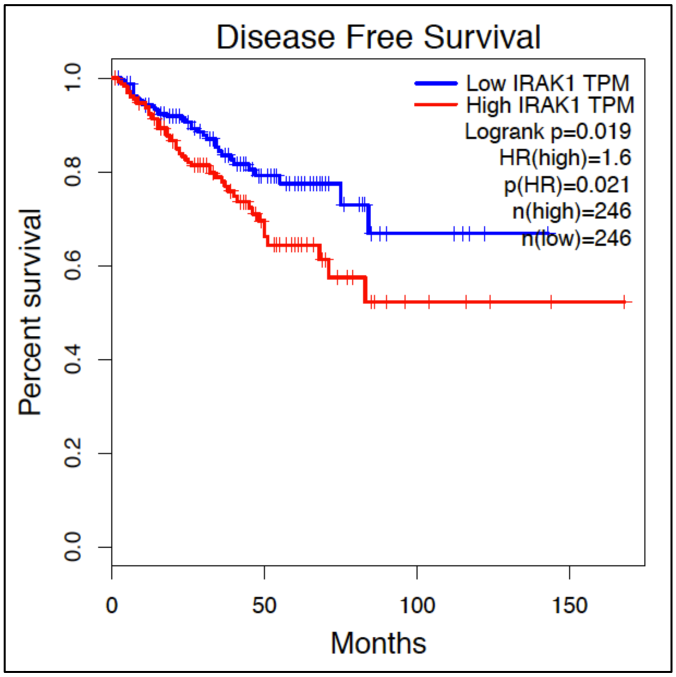


**Supplementary Figure S4:** Kaplan-Meier DFS survival analysis for high-expressing vs low-expressing PRAD samples. Kaplan-Meier survival analysis was performed for the PRAD dataset (n=472). PRAD patients were stratified into two groups of IRAK1 high-expressing patients (n=246) vs IRAK1 low-expressing patients (n=246). Log-rank p-value and Hazard ratio were calculated for disease-free survival status (DFS) using the median expression value for all samples as the cut-off.

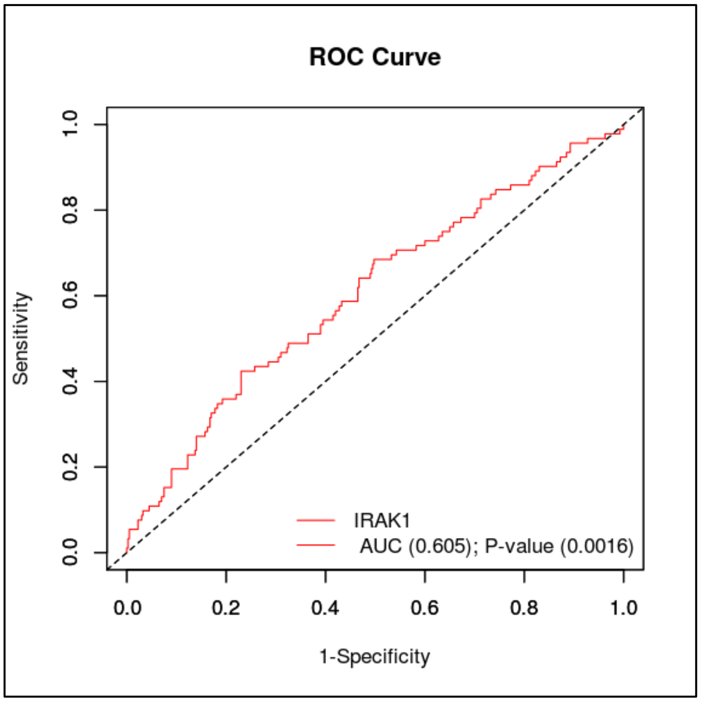


**Supplementary Figure S5:** Receiver Operating Characteristic (ROC) analysis for IRAK in predicting cancer relapse. This ROC analyzes the capability of IRAK1, a representative of inflammation-associated genes in the magenta module to predict survival risk. IRAK1 was able to predict 60% of PRAD relapse with an AUC of 0.6 for relapse vs non-relapse patients. The sensitivity is represented on the x-axis while 1-specificity is represented on the y-axis. The black dotted lines represent the line of significance (p = 0.05). The ROC curves represent the gold standard of diagnostic accuracy (any gene with AUC >0.6 is assumed to be of good clinical significance and potential diagnostic biomarker) on the upper and left axes in the unit square.

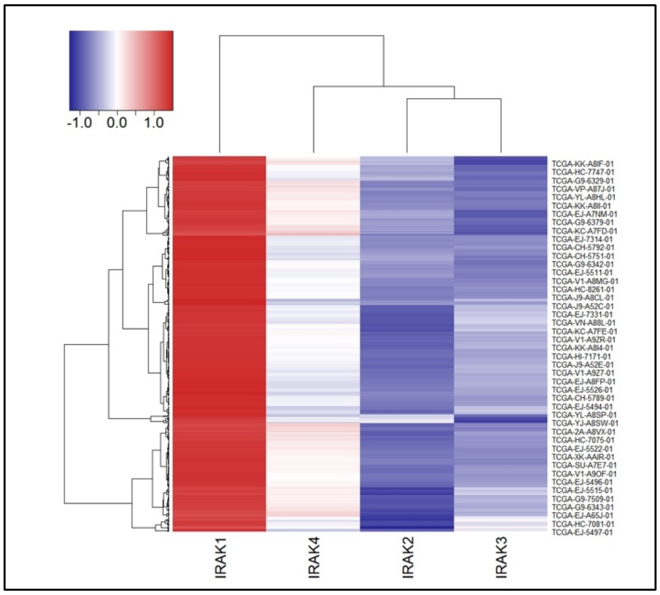


**Supplementary Figure S6:** Heatmap of the expression profile of IRAK family genes in indolent PRAD patients (n = 472) showing the overexpression of IRAK1 compared to other IRAKs.

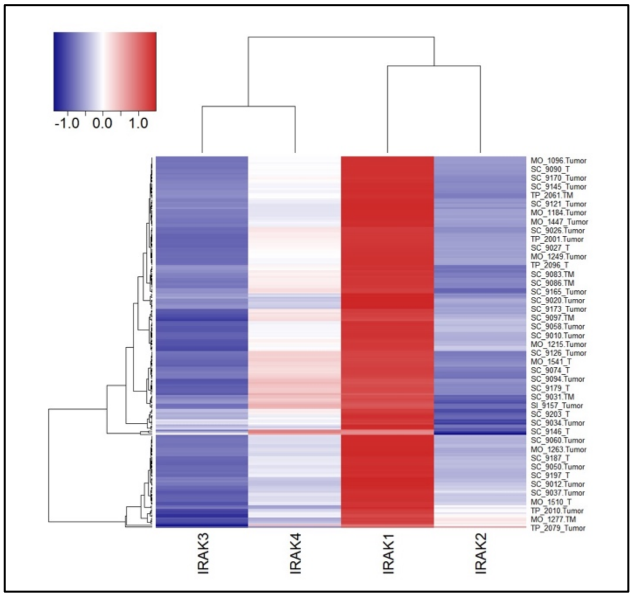


**Supplementary Figure S7:** Heatmap of the expression profile of IRAK family genes in neuroendocrine PCa patients (n = 30) showing the overexpression of IRAK1 compared to other IRAKs**.**

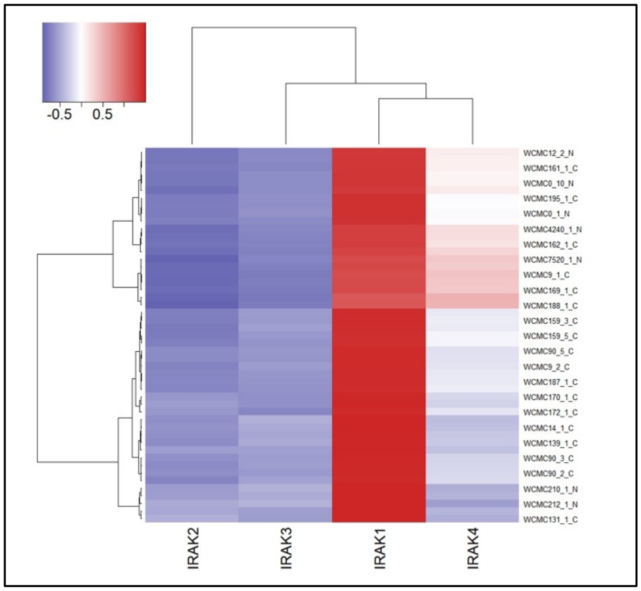


**Supplementary Figure S8:** Heatmap of the expression profile of IRAK family genes in castration-resistant PCa patients (n = 444) showing the overexpression of IRAK1 compared to other IRAKs.

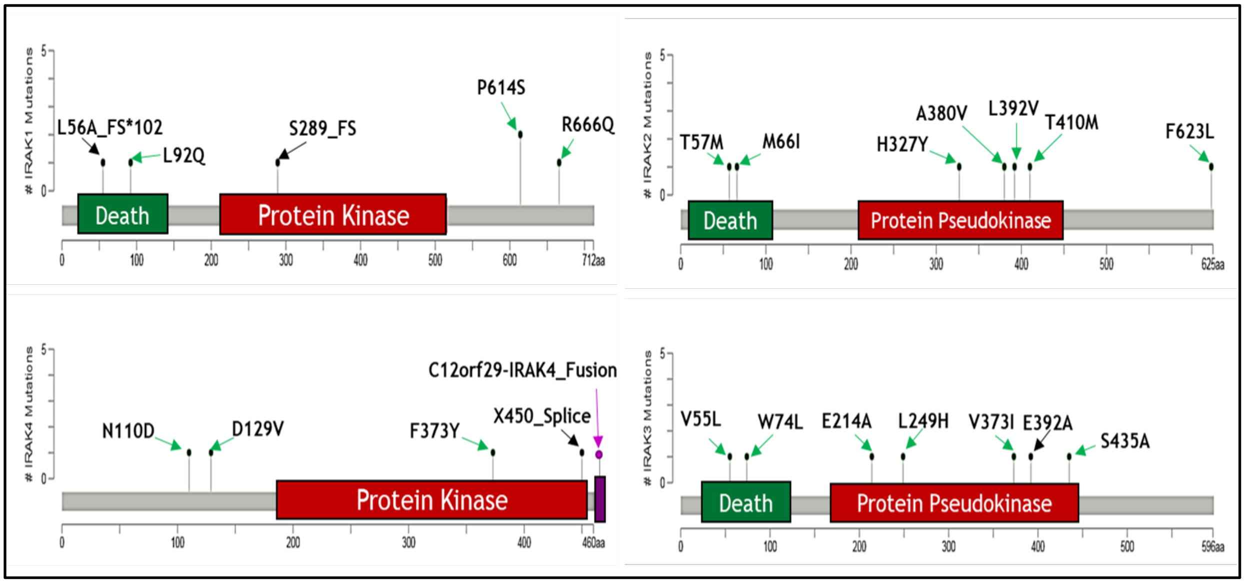


**Supplementary Figure S9:** Lollipop plots showing the identified structural mutations of IRAK1, IRAK2, IRAK3, and IRAK4 in PCa patients from the 20 cohort studies. We observed missense (green arrow), frameshift mutations (black arrow), and fusion mutations. The missense and frameshift mutations were mostly found in the death domains and protein kinase/pseudokinase domains of IRAKs. No missense mutation was found in all the CRPC or NEPC patients analyzed using tools such as MutSigCV.

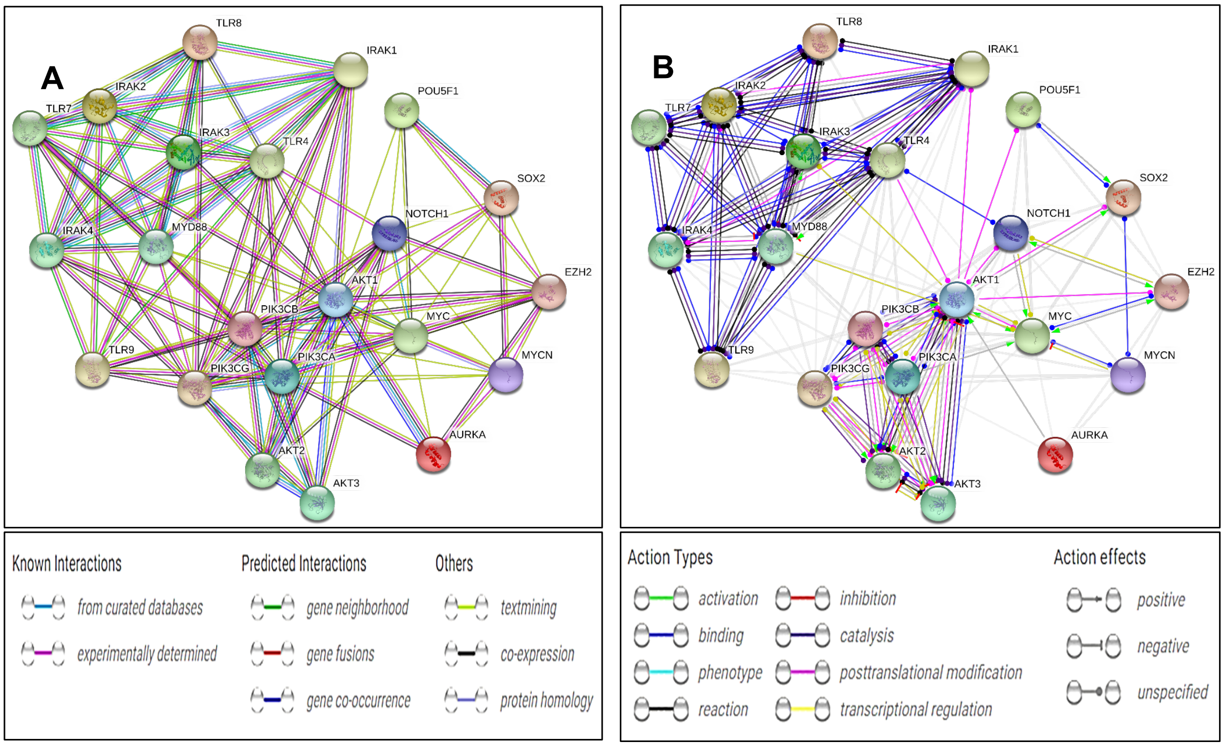


**Supplementary Figure S10:** Protein-protein interaction and enrichment network analyses in PRAD. **A.** Evidence-based protein-protein interaction (PPI) network analysis between IRAK1 and selected inflammation and PCa-progression proteins. **B.** mode of action protein-protein interaction (PPI) network analyses between IRAK1 and selected inflammation and PCa-progression proteins. Pathway interactions were determined using evidence and mode of action associated functional annotations/gene ontology which predicts the degree of interaction between proteins as determined using the String_db (v11) software. Our PPI analysis as shown on the right panel predicted a posttranslational modification of IRAK1 by AKT1, IRAK2, and IRAK4 as well as uncharacterized effects on PI3K and AKT genes. IRAK1 could also directly bind to TLR4, TLR8, TLR9, MYD88, IRAK2, IRAK3, and IRAK4. Predicted interactions between IRAK1 and these genes include gene neighborhood, gene fusions, gene co-occurrence, gene co-expression, and protein homology.

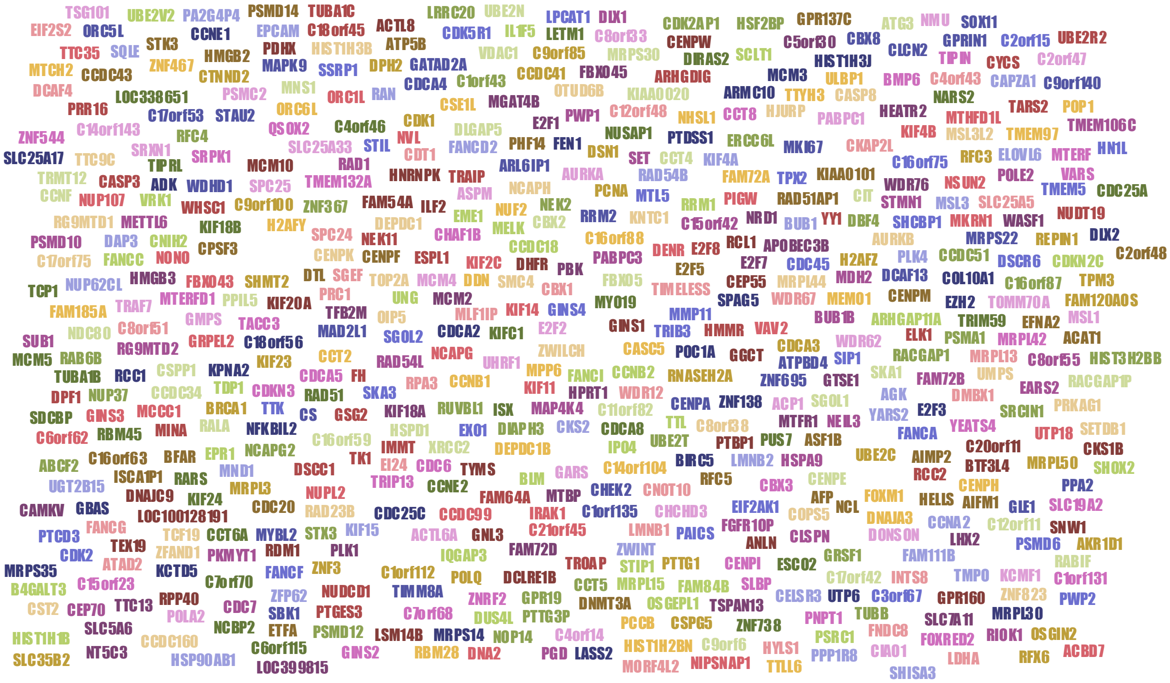

**Supplementary Figure S11:** Genes Word-Cloud Generator showing all the 588 genes identified in the magenta module.

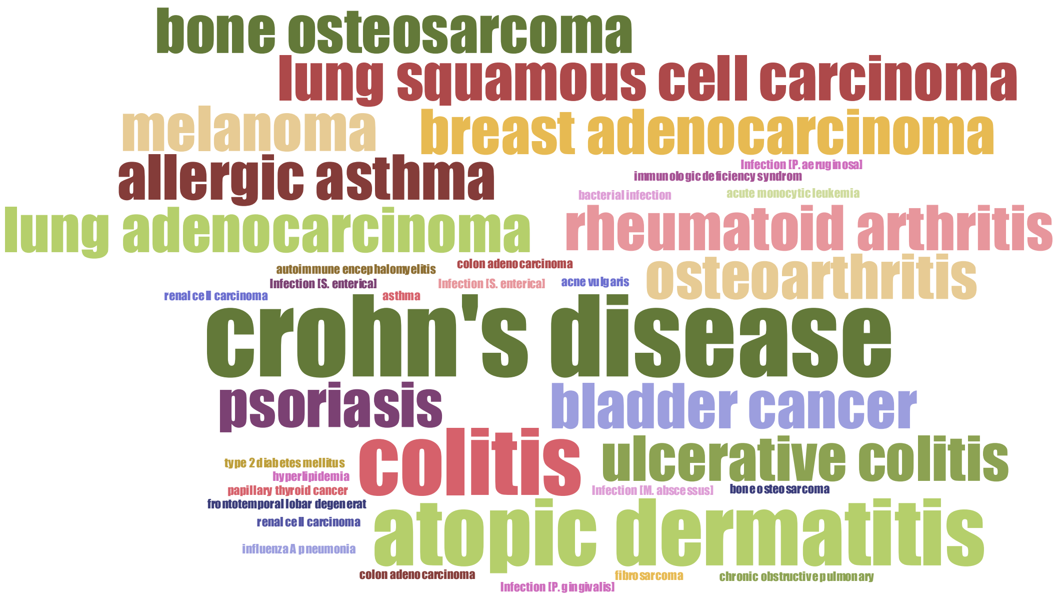


**Supplementary Figure S12:** Diseases Word-Cloud Generator showing all the diseases associated with aberrant IRAK1 signaling. This is based on results of studies text-mined from scientific literature (PubMed and Google Scholar) and data-mined using the activity plot ($p < 0.05$) in Ingenuity Pathway Analysis (IPA) software by Qiagen.

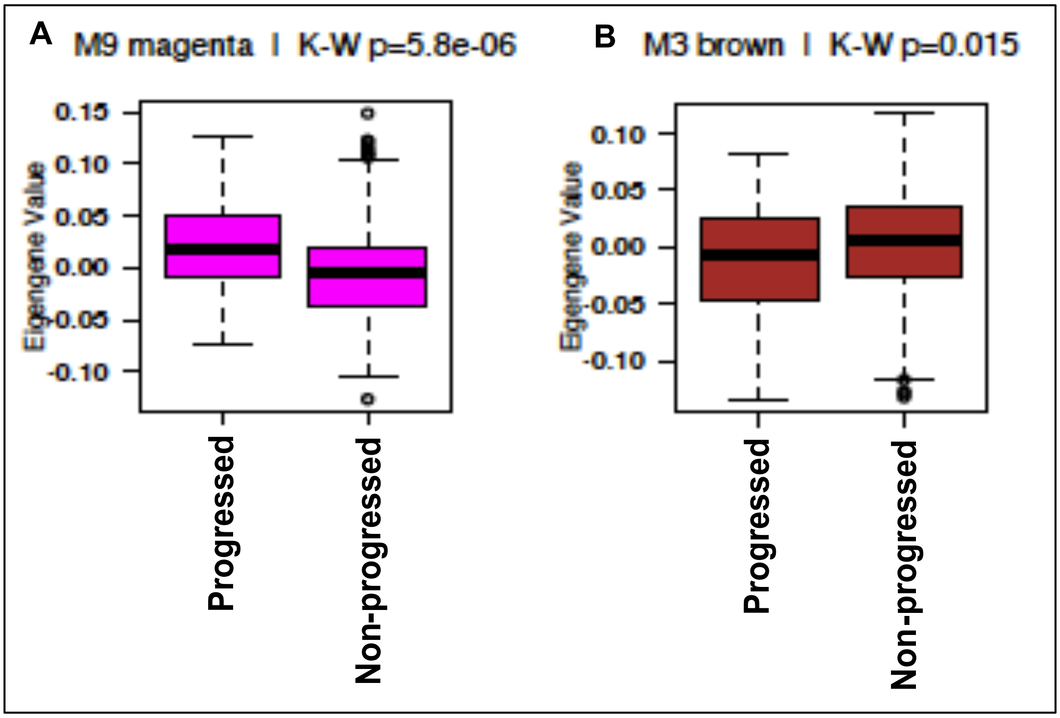


**Supplementary Figure S13:** Boxplots to compare Eigengene values between progressed vs non-progressed patients in the magenta and brown modules. The degree of significance between progressed and non-progressed PRAD patients was determined using Kruskal-Wallis (K-W) test or one-way ANOVA. **A.** Boxplot showing the difference between the median eigengene values (of genes in the magenta module) between progressed PRAD patients and the non-progressed patients. This indicates a significant upregulation ($p = 5.8e-06$) and connectedness of genes of the magenta (M9) module in progressed patients compared to the non-progressed PRAD patients. **B.** Boxplot showing the difference between the median eigengene values (of genes in the brown module) between progressed and non-progressed PRAD patients. This indicates a significant downregulation ($p = 0.015$) of genes of the brown (M3) module in progressed patients compared to the non-progressed PRAD patients. IRAK1 is one of the 10 inflammatory hub genes identified in the magenta module, highly expressed in

PRAD patients as well as progressed patients. Whereas IRAK3 is one of the inflammatory genes in the brown module. IRAK3 was observed to be differentially downregulated in PRAD samples/patients as well as progressed compared to non-progressed patients.

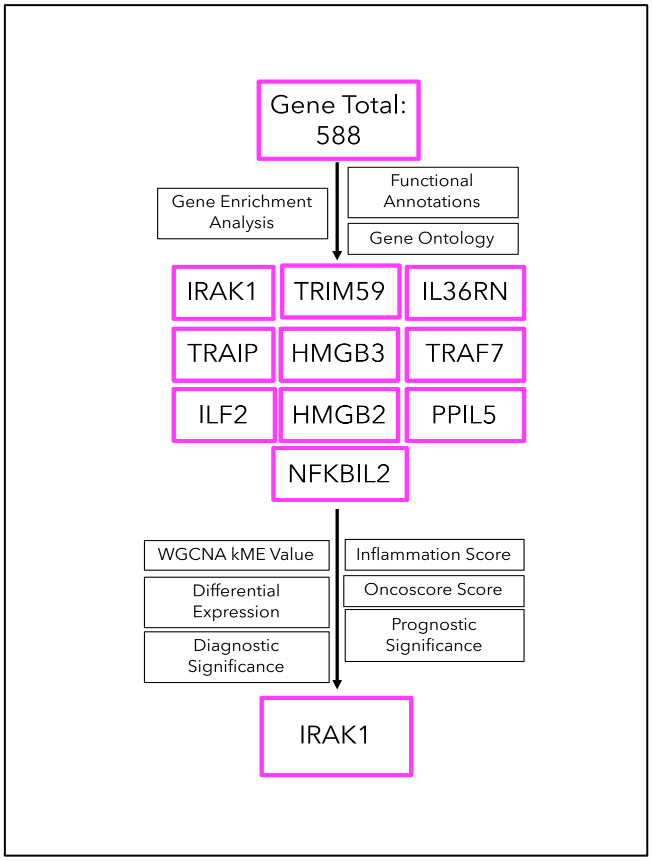


**Supplementary Figure S14:** A flow chart summarizing the post hoc multivariate analyses undertaken for screening and identification of significant inflammatory genes in the magenta module.

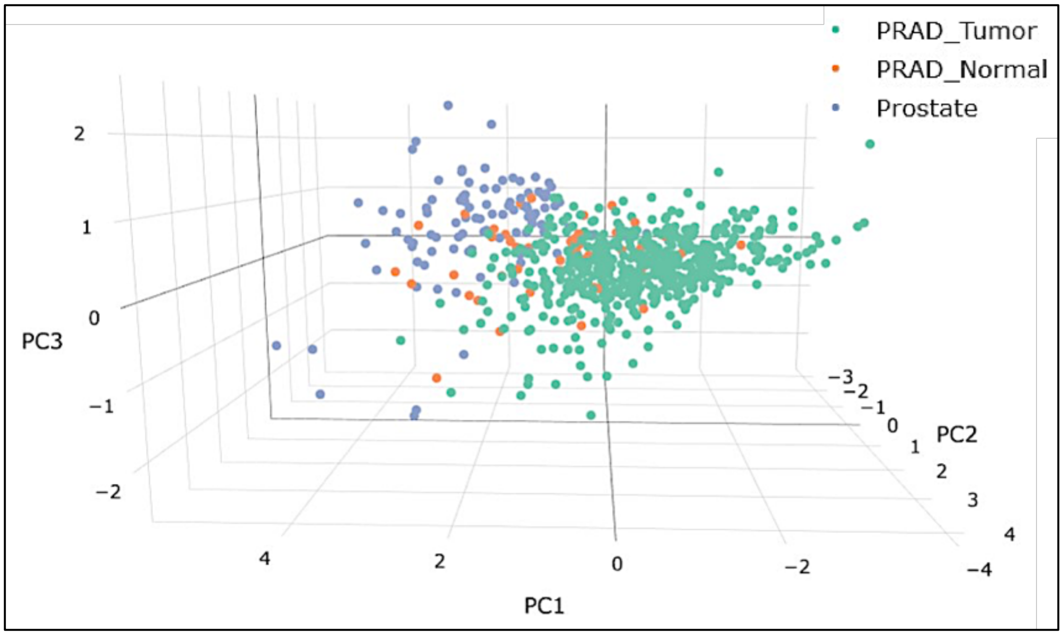


**Supplementary Figure S15:** A principal component analysis (PCA) dimensionality reduction of IRAK1 expression in PRAD and non-PRAD normal samples. The 3D PCA plot demonstrates a capability to separate expression of IRAK1 in PRAD samples (n=494; green dots) from matched normal prostate samples from PRAD patients (n = 52; orange dots) and unmatched normal prostate samples (downloaded from GTEx database, n=245; blue dots) from PRAD patients and non-PRAD patients using PCA analysis tool of GEPIA2 (http://gepia2.cancer-pku.cn/#dimension).

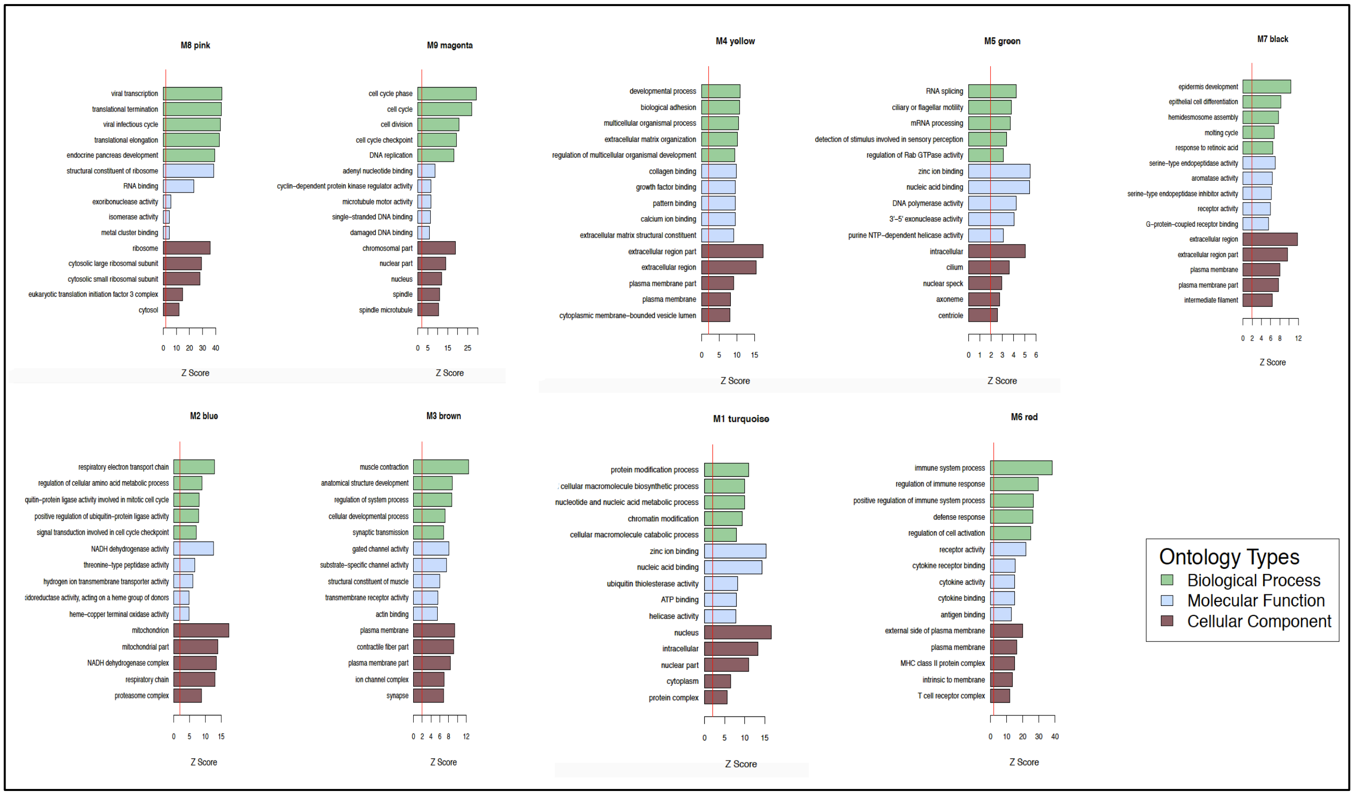


**Supplementary Figure S16:** GO Elite ontologies of the 9 biologically significant modules. The x-axis represents the calculated Z-scores for the gene ontologies. The y axis has the biological processes (green bars), the molecular functions (blue bars), and the cellular components for each module (brown bars). The red vertical lines shown across the bars represent the direction of the standard deviations.

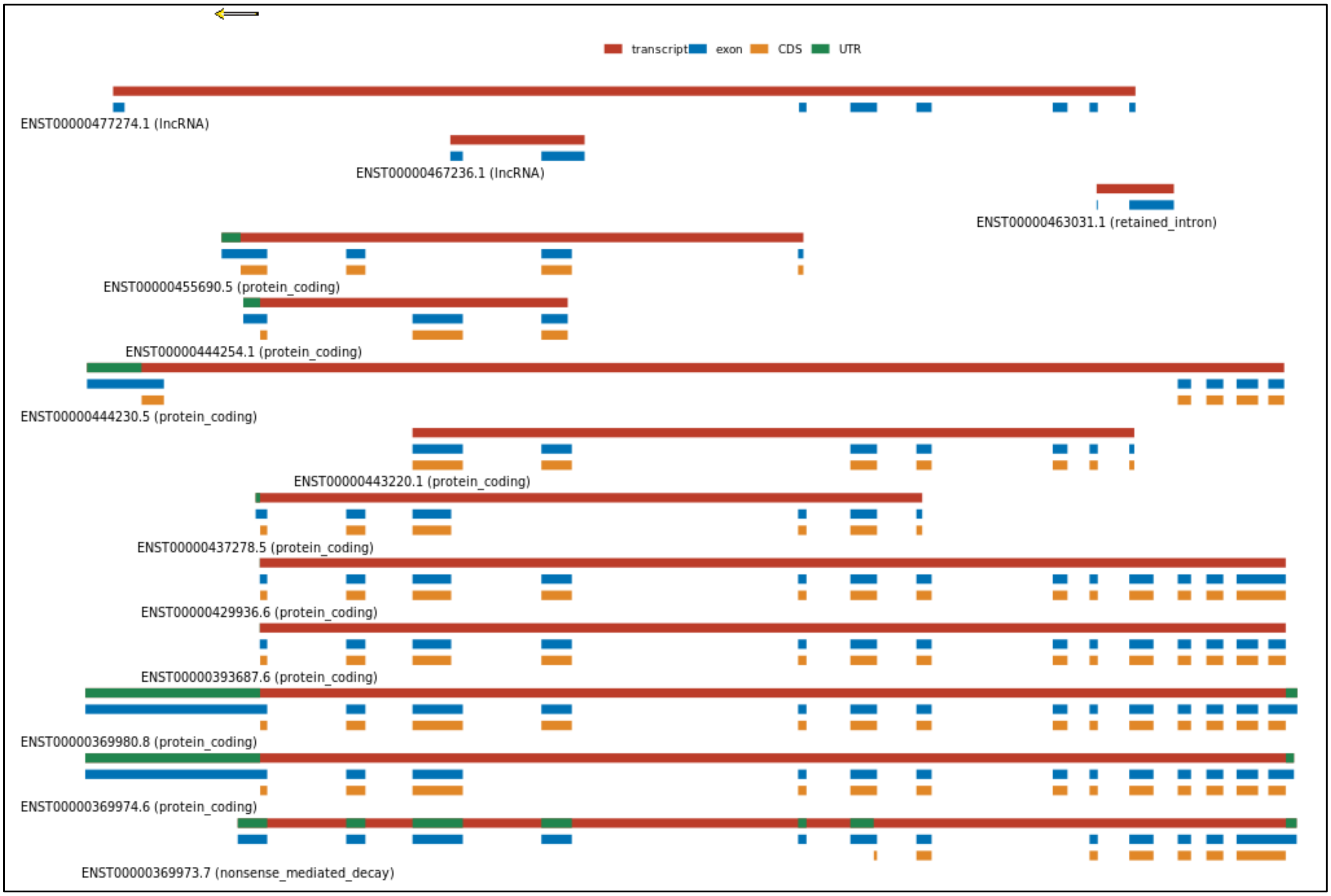


**Supplementary Figure S17:** A detailed genomic sketch of IRAK1 showing information on the 13 transcripts/isoforms identified as well as exons, gene coding sequence regions (CDS), and untranslated regions (UTR).

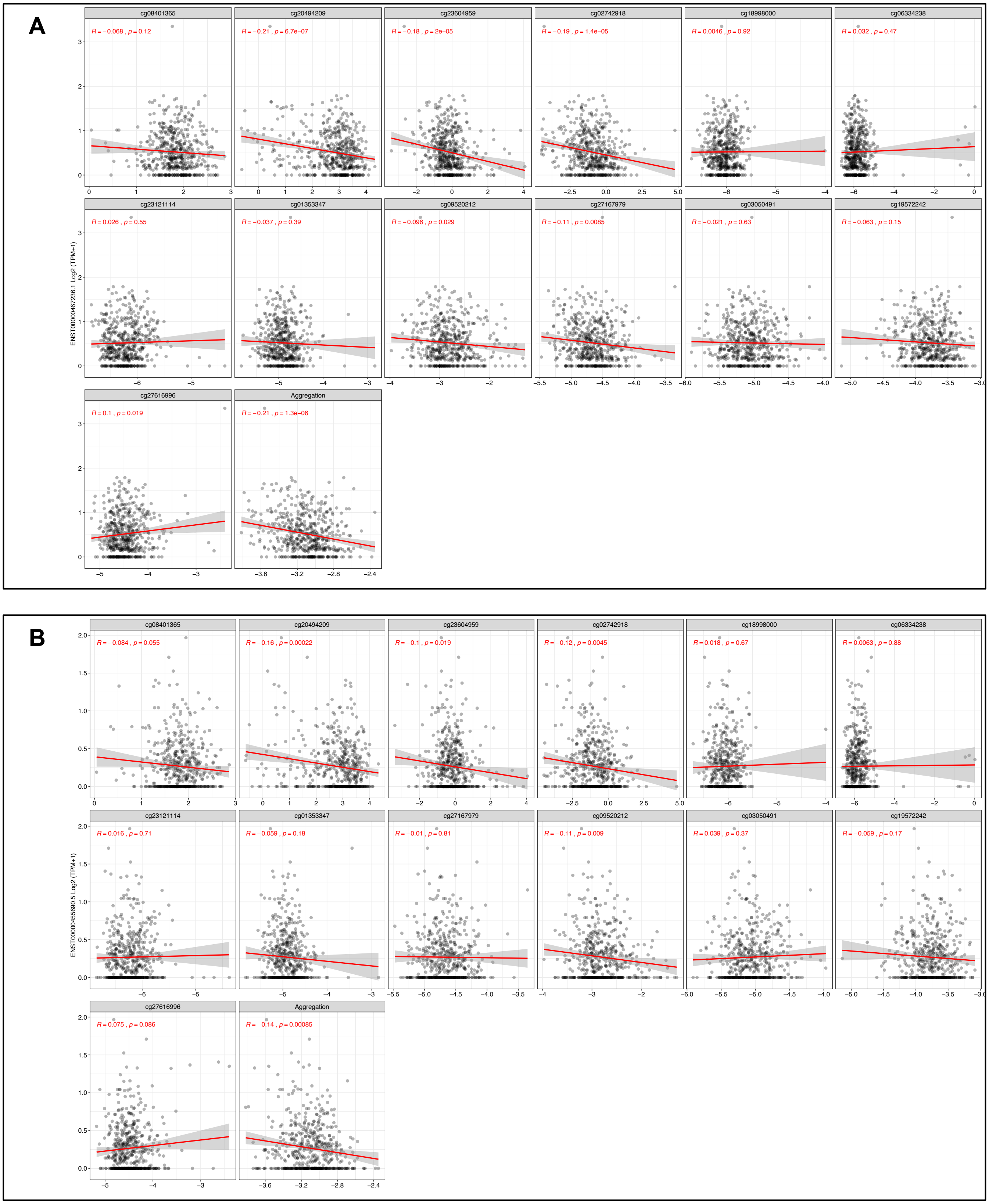
A
ENST00000467236.1 Log2 (TPM+1)
cg08401365
R = −0.068 , p = 0.12
cg20494209
R = −0.21 , p = 6.7e−07
cg23604959
R = −0.18 , p = 2e−05
cg02742918
R = −0.19 , p = 1.4e−05
cg18998000
R = 0.0046 , p = 0.92
cg06334238
R = 0.032 , p = 0.47
cg23121114
R = 0.026 , p = 0.55
cg01353347
R = −0.037 , p = 0.39
cg09520212
R = −0.096 , p = 0.029
cg27167979
R = −0.11 , p = 0.0085
cg03050491
R = −0.021 , p = 0.63
cg19572242
R = −0.063 , p = 0.15
cg27616996
R = 0.1 , p = 0.019
Aggregation
R = −0.21 , p = 1.3e−06
B
ENST00000455690.5 Log2 (TPM+1)
cg08401365
R = −0.084 , p = 0.055
cg20494209
R = −0.16 , p = 0.00022
cg23604959
R = −0.1 , p = 0.019
cg02742918
R = −0.12 , p = 0.0045
cg18998000
R = 0.018 , p = 0.67
cg06334238
R = 0.0063 , p = 0.88
cg23121114
R = 0.016 , p = 0.71
cg01353347
R = −0.059 , p = 0.18
cg27167979
R = −0.01 , p = 0.81
cg09520212
R = −0.11 , p = 0.009
cg03050491
R = 0.039 , p = 0.37
cg19572242
R = −0.059 , p = 0.17
cg27616996
R = 0.075 , p = 0.086
Aggregation
R = −0.14 , p = 0.00085

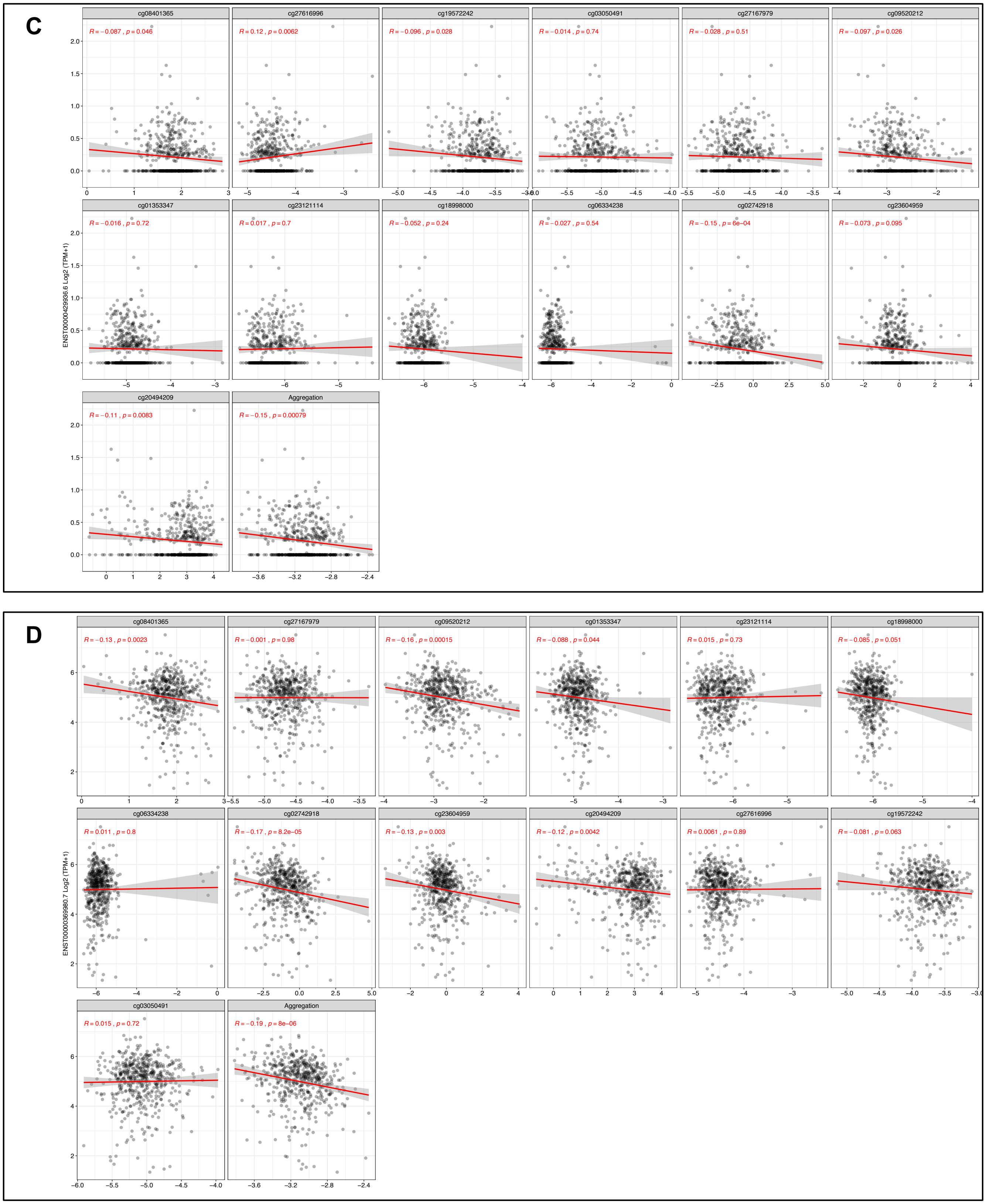
C
ENST00000429936.6 Log2 (TPM+1)
cg08401365
R = −0.087 , p = 0.046
cg27616996
R = 0.12 , p = 0.0062
cg19572242
R = −0.096 , p = 0.028
cg03050491
R = −0.014 , p = 0.74
cg27167979
R = −0.028 , p = 0.51
cg09520212
R = −0.097 , p = 0.026
cg01353347
R = −0.016 , p = 0.72
cg23121114
R = 0.017 , p = 0.7
cg18998000
R = −0.052 , p = 0.24
cg06334238
R = −0.027 , p = 0.54
cg02742918
R = −0.15 , p = 6e−04
cg23604959
R = −0.073 , p = 0.095
cg20494209
R = −0.11 , p = 0.0083
Aggregation
R = −0.15 , p = 0.00079
D
ENST00000369980.7 Log2 (TPM+1)
cg08401365
R = −0.13 , p = 0.0023
cg27167979
R = −0.001 , p = 0.98
cg09520212
R = −0.16 , p = 0.00015
cg01353347
R = −0.088 , p = 0.044
cg23121114
R = 0.015 , p = 0.73
cg18998000
R = −0.085 , p = 0.051
cg06334238
R = 0.011 , p = 0.8
cg02742918
R = −0.17 , p = 8.2e−05
cg23604959
R = −0.13 , p = 0.003
cg20494209
R = −0.12 , p = 0.0042
cg27616996
R = 0.0061 , p = 0.89
cg19572242
R = −0.081 , p = 0.063
cg03050491
R = 0.015 , p = 0.72
Aggregation
R = −0.19 , p = 8e−06

**E**

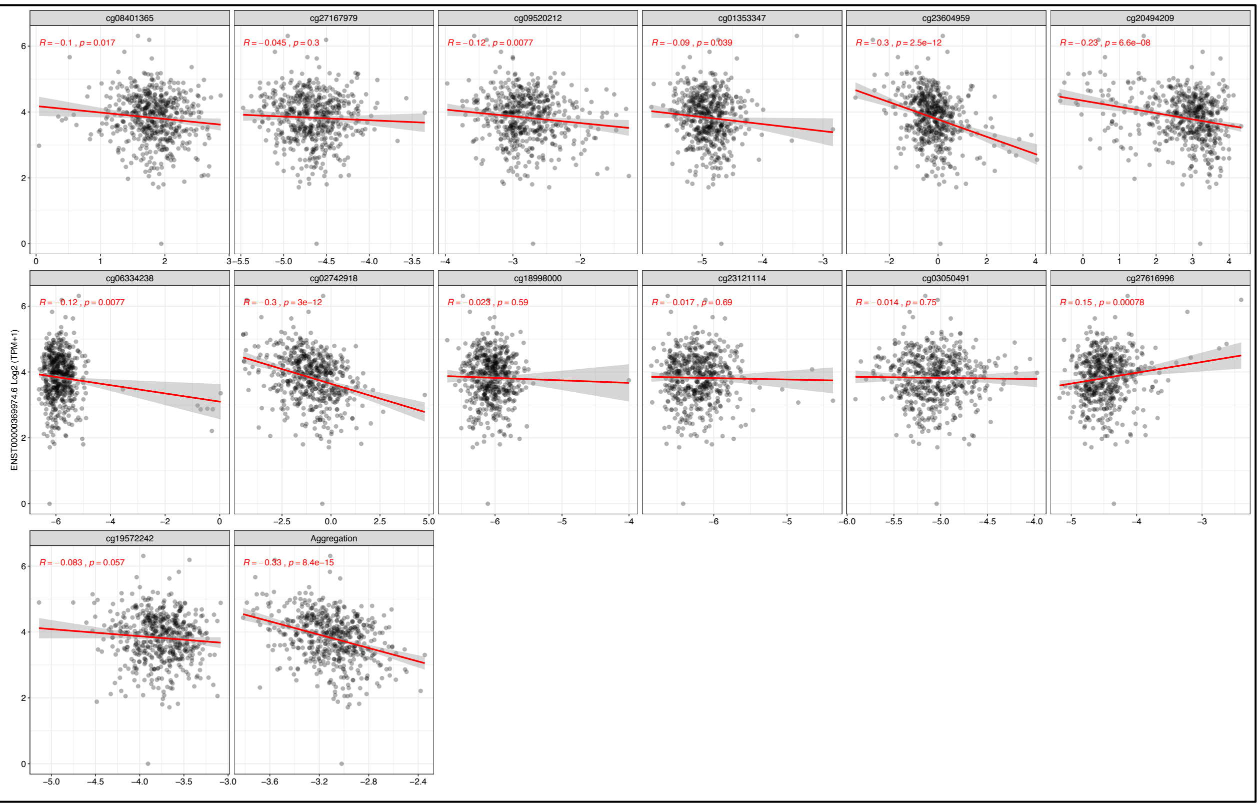
ENST00000369974.6 Log2 (TPM+1)
cg08401365
R = −0.1 , p = 0.017
cg27167979
R = −0.045 , p = 0.3
cg09520212
R = −0.12 , p = 0.0077
cg01353347
R = −0.09 , p = 0.039
cg23604959
R = −0.3 , p = 2.5e−12
cg20494209
R = −0.23 , p = 6.6e−08
cg06334238
R = −0.12 , p = 0.0077
cg02742918
R = −0.3 , p = 3e−12
cg18998000
R = −0.023 , p = 0.59
cg23121114
R = −0.017 , p = 0.69
cg03050491
R = −0.014 , p = 0.75
cg27616996
R = 0.15 , p = 0.00078
cg19572242
R = −0.083 , p = 0.057
Aggregation
R = −0.33 , p = 8.4e−15

**F**

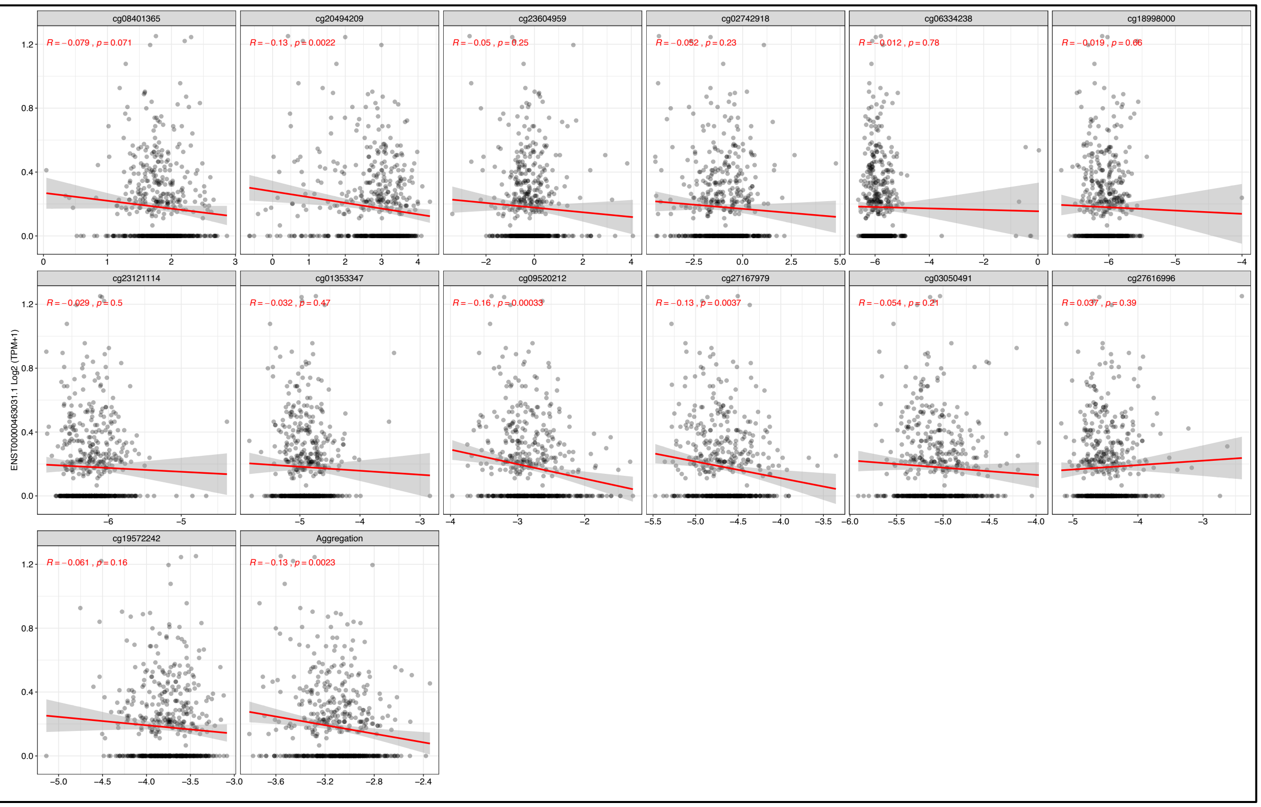
ENST00000463031.1 Log2 (TPM+1)
cg08401365
R = −0.079 , p = 0.071
cg20494209
R = −0.13 , p = 0.0022
cg23604959
R = −0.05 , p = 0.25
cg02742918
R = −0.052 , p = 0.23
cg06334238
R = −0.012 , p = 0.78
cg18998000
R = −0.019 , p = 0.66
cg23121114
R = −0.029 , p = 0.5
cg01353347
R = −0.032 , p = 0.47
cg09520212
R = −0.16 , p = 0.00033
cg27167979
R = −0.13 , p = 0.0037
cg03050491
R = −0.054 , p = 0.21
cg27616996
R = 0.037 , p = 0.39
cg19572242
R = −0.061 , p = 0.16
Aggregation
R = −0.13 , p = 0.0023

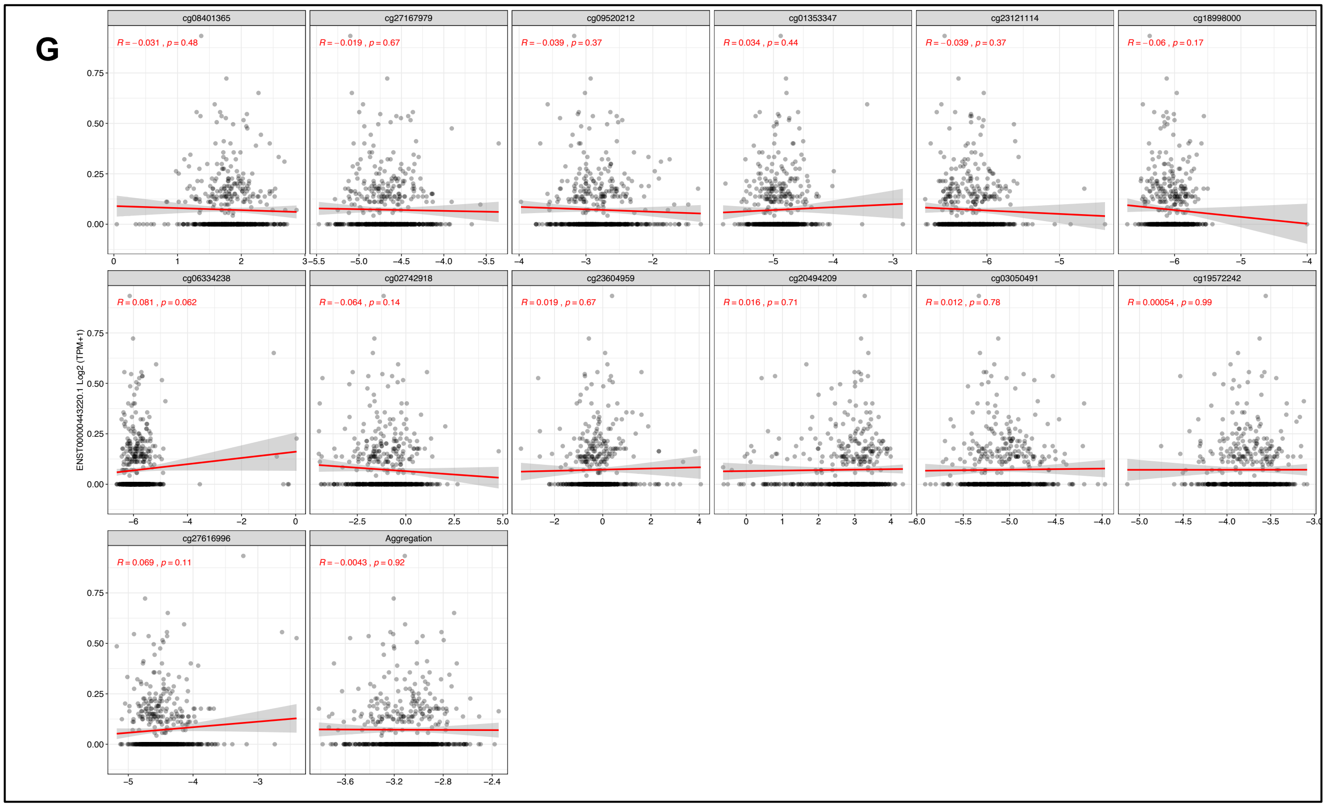

G
ENST00000443220.1 Log2 (TPM+1)
cg08401365
R = −0.031 , p = 0.48
cg27167979
R = −0.019 , p = 0.67
cg09520212
R = −0.039 , p = 0.37
cg01353347
R = 0.034 , p = 0.44
cg23121114
R = −0.039 , p = 0.37
cg18998000
R = −0.06 , p = 0.17
cg06334238
R = 0.081 , p = 0.062
cg02742918
R = −0.064 , p = 0.14
cg23604959
R = 0.019 , p = 0.67
cg20494209
R = 0.016 , p = 0.71
cg03050491
R = 0.012 , p = 0.78
cg19572242
R = 0.00054 , p = 0.99
cg27616996
R = 0.069 , p = 0.11
Aggregation
R = −0.0043 , p = 0.92


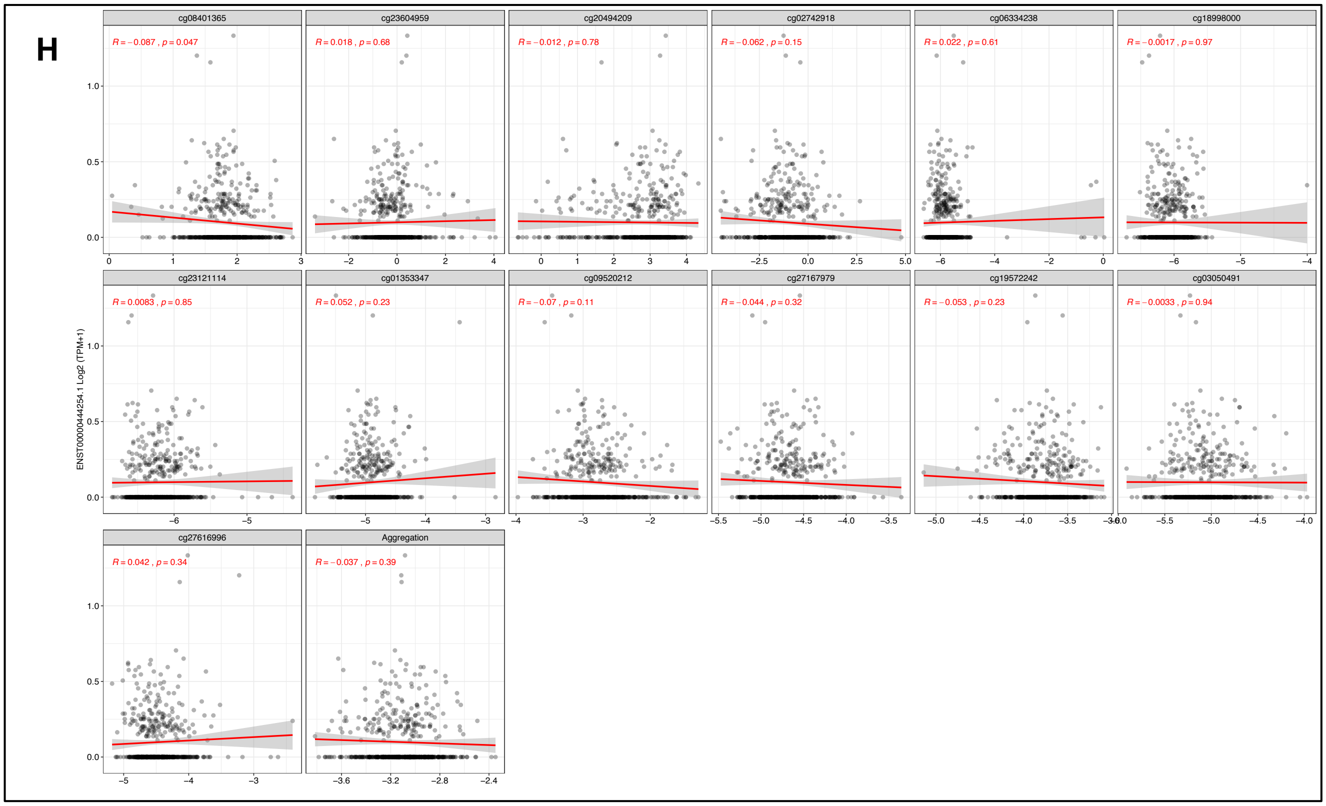

H
ENST00000444254.1 Log2 (TPM+1)
cg08401365
R = −0.087 , p = 0.047
cg23604959
R = 0.018 , p = 0.68
cg20494209
R = −0.012 , p = 0.78
cg02742918
R = −0.062 , p = 0.15
cg06334238
R = 0.022 , p = 0.61
cg18998000
R = −0.0017 , p = 0.97
cg23121114
R = 0.0083 , p = 0.85
cg01353347
R = 0.052 , p = 0.23
cg09520212
R = −0.07 , p = 0.11
cg27167979
R = −0.044 , p = 0.32
cg19572242
R = −0.053 , p = 0.23
cg03050491
R = −0.0033 , p = 0.94
cg27616996
R = 0.042 , p = 0.34
Aggregation
R = −0.037 , p = 0.39

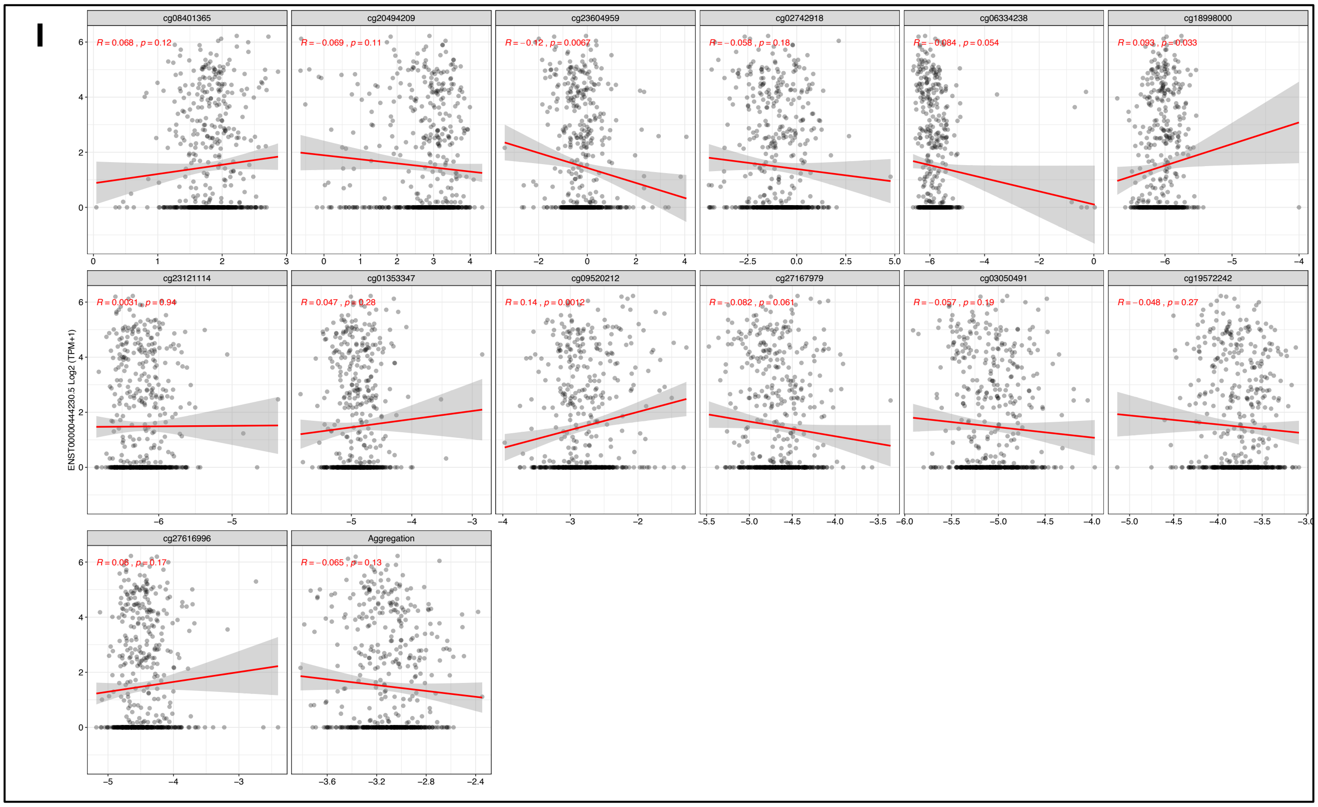
I
ENST00000444230.5 Log2 (TPM+1)
cg08401365
R = 0.068 , p = 0.12
cg20494209
R = −0.069 , p = 0.11
cg23604959
R = −0.12 , p = 0.0067
cg02742918
R = −0.058 , p = 0.18
cg06334238
R = −0.084 , p = 0.054
cg18998000
R = 0.093 , p = 0.033
cg23121114
R = 0.0031 , p = 0.94
cg01353347
R = 0.047 , p = 0.28
cg09520212
R = 0.14 , p = 0.0012
cg27167979
R = −0.082 , p = 0.061
cg03050491
R = −0.057 , p = 0.19
cg19572242
R = −0.048 , p = 0.27
cg27616996
R = 0.06 , p = 0.17
Aggregation
R = −0.065 , p = 0.13

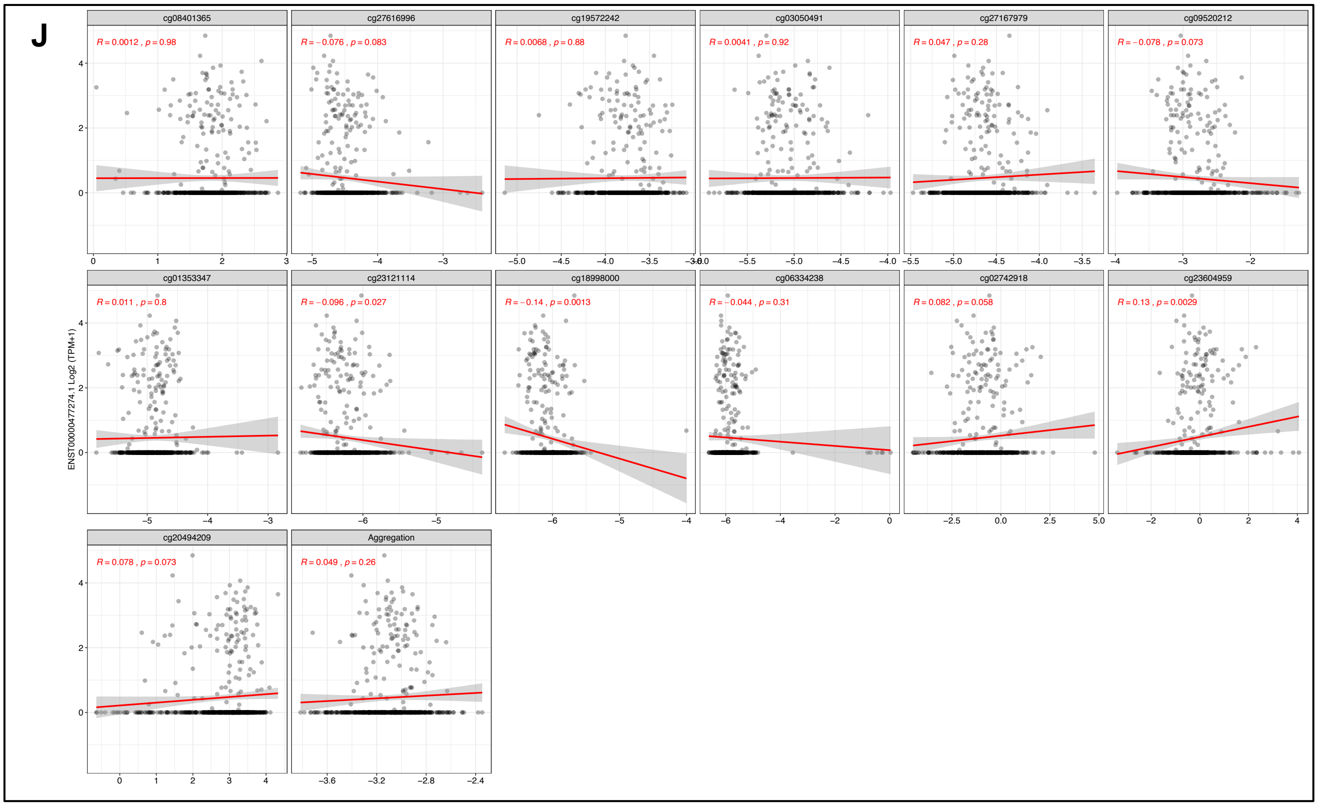
J
ENST00000477274.1 Log2 (TPM+1)
cg08401365
R = 0.0012 , p = 0.98
cg27616996
R = −0.076 , p = 0.083
cg19572242
R = 0.0068 , p = 0.88
cg03050491
R = 0.0041 , p = 0.92
cg27167979
R = 0.047 , p = 0.28
cg09520212
R = −0.078 , p = 0.073
cg01353347
R = 0.011 , p = 0.8
cg23121114
R = −0.096 , p = 0.027
cg18998000
R = −0.14 , p = 0.0013
cg06334238
R = −0.044 , p = 0.31
cg02742918
R = 0.082 , p = 0.058
cg23604959
R = 0.13 , p = 0.0029
cg20494209
R = 0.078 , p = 0.073
Aggregation
R = 0.049 , p = 0.26

K

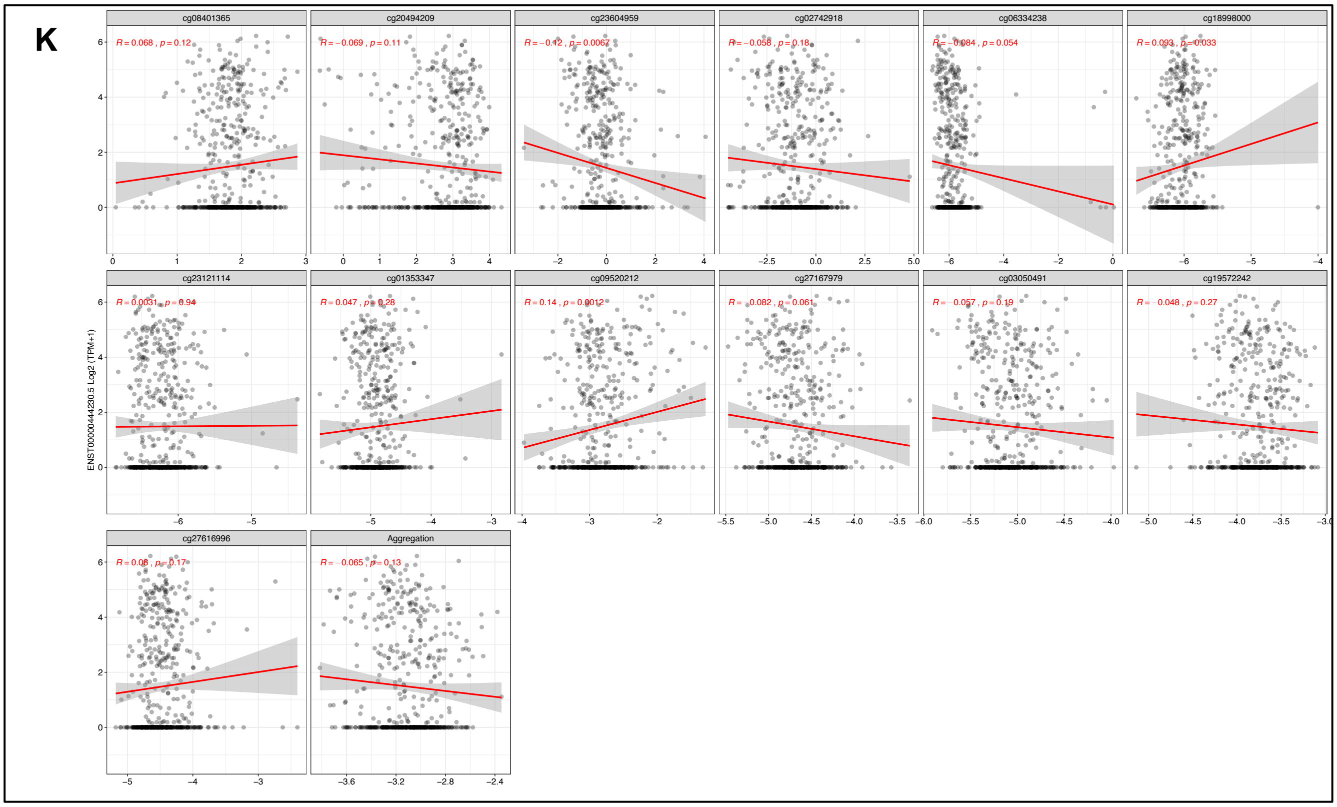
ENST00000444230.5 Log2 (TPM+1)
cg08401365
R = 0.068 , p = 0.12
cg20494209
R = −0.069 , p = 0.11
cg23604959
R = −0.12 , p = 0.0067
cg02742918
R = −0.058 , p = 0.18
cg06334238
R = −0.084 , p = 0.054
cg18998000
R = 0.093 , p = 0.033
cg23121114
R = 0.0031 , p = 0.94
cg01353347
R = 0.047 , p = 0.28
cg09520212
R = 0.14 , p = 0.0012
cg27167979
R = −0.082 , p = 0.061
cg03050491
R = −0.057 , p = 0.19
cg19572242
R = −0.048 , p = 0.27
cg27616996
R = 0.06 , p = 0.17
Aggregation
R = −0.065 , p = 0.13

L

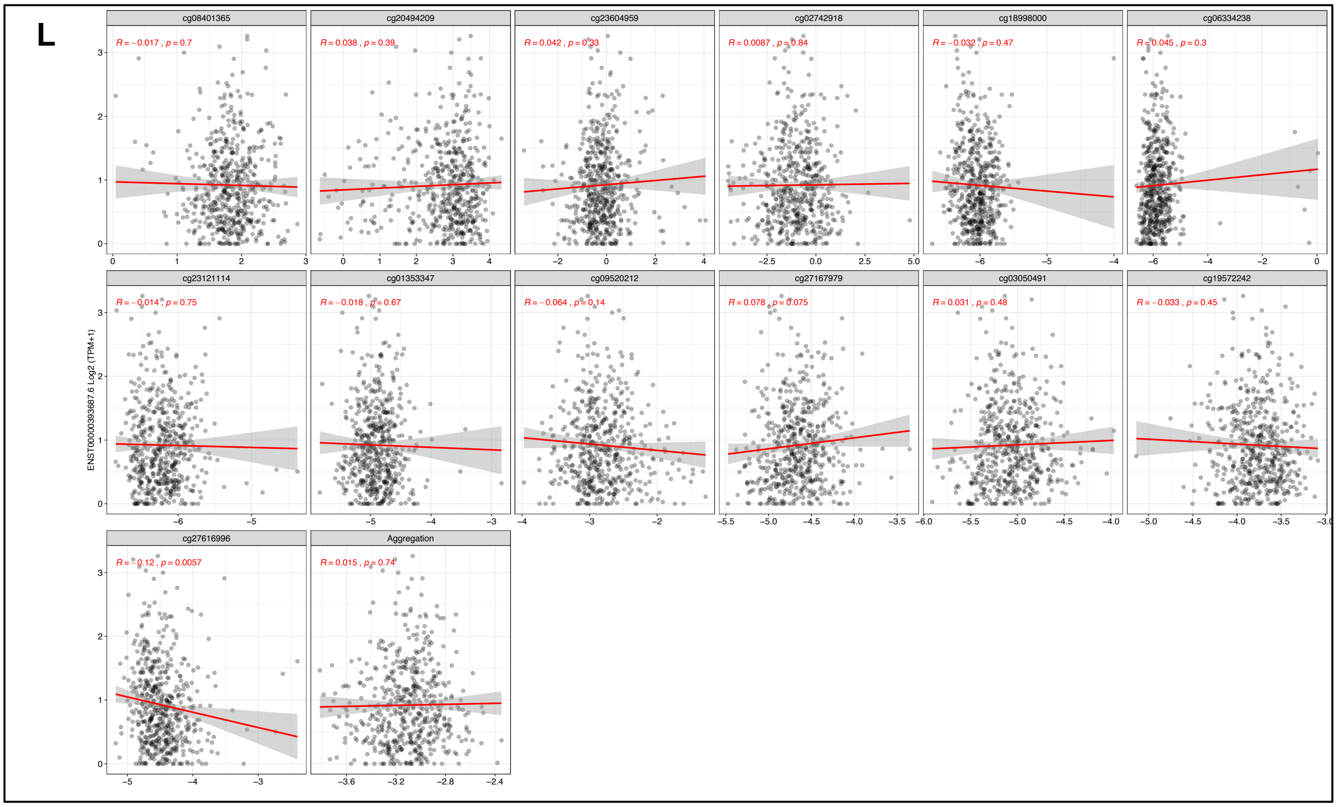
ENST00000393687.6 Log2 (TPM+1)
cg08401365
R = −0.017 , p = 0.7
cg20494209
R = 0.038 , p = 0.39
cg23604959
R = 0.042 , p = 0.33
cg02742918
R = 0.0087 , p = 0.84
cg18998000
R = −0.032 , p = 0.47
cg06334238
R = 0.045 , p = 0.3
cg23121114
R = −0.014 , p = 0.75
cg01353347
R = −0.018 , p = 0.67
cg09520212
R = −0.064 , p = 0.14
cg27167979
R = 0.078 , p = 0.075
cg03050491
R = 0.031 , p = 0.48
cg19572242
R = −0.033 , p = 0.45
cg27616996
R = −0.12 , p = 0.0057
Aggregation
R = 0.015 , p = 0.74

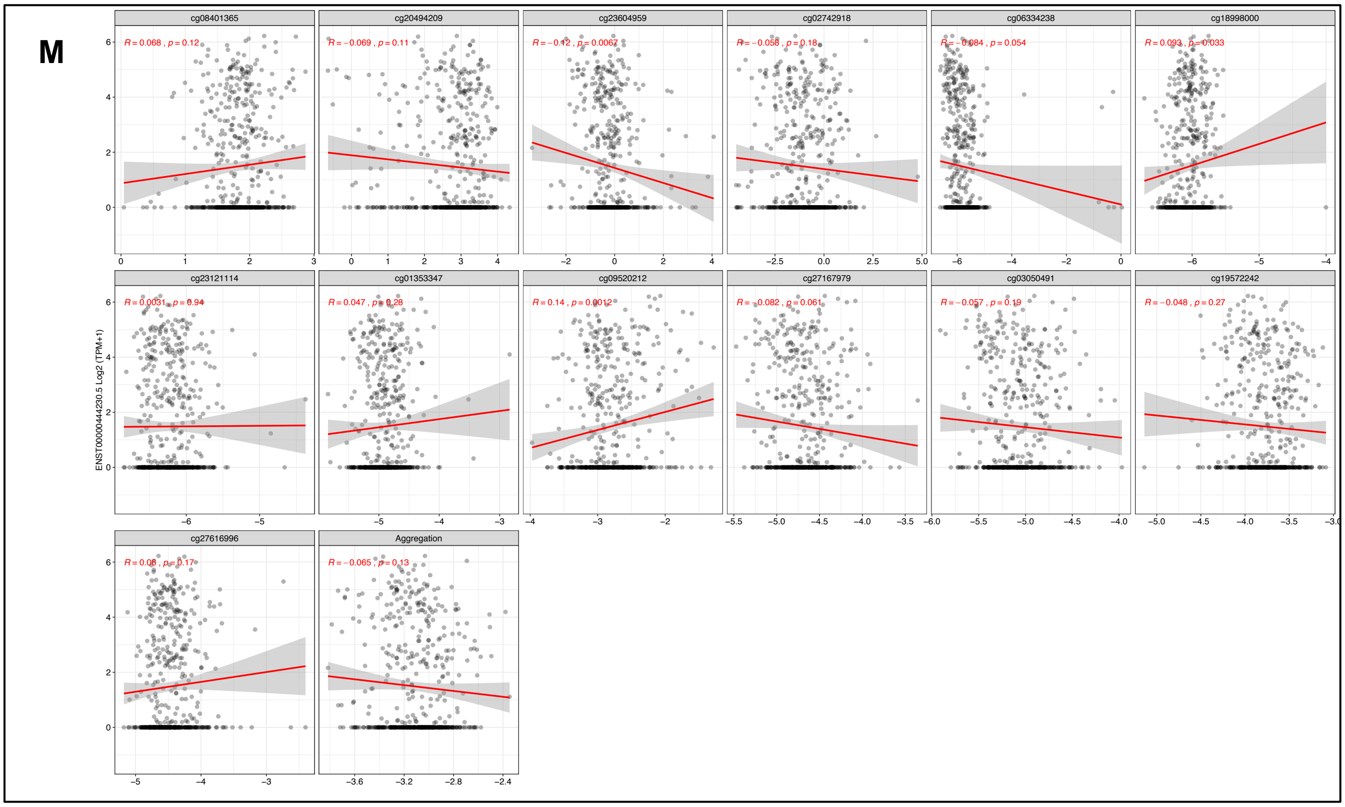


**Supplementary Figure S18 (A-M):** Correlation analysis plots using Pearson coefficient to identify the correlation between various IRAK1 CpG methylation values and the expression levels of IRAK1 isoforms. Only 6 (ENST00000369974.6 (r = -0.338; p = 8.4e-15), ENST00000369980.7 (r = -0.19; p = 8e-06), ENST00000429936.6 (r = -0.15; p = 0.00079), ENST00000455690.5 (r = -0.14; p = 0.00085), ENST00000463031.1 (r = -0.13; p = 0.0023), ENST00000467236.1 (r = -0.21; p = 1.36e-6)) of the 13 isoforms identified show negative correlation ($p < 0.05$) with the methylation values of IRAK1.
